# Chromosome axis protein SYCP2 recruits HORMAD2 to enable meiotic synapsis quality control in mice

**DOI:** 10.64898/2026.01.06.697704

**Authors:** Kavya Raveendran, Sarai Valerio-Cabrera, Arkasarathi Gope, Vladyslav Telychko, Geen George, Christin Richter, Tanja Scholte, Matthias Weigel, Anastasiia Bondarieva, Andreas Petzold, Andreas Dahl, Kevin D Corbett, Attila Tóth

## Abstract

Faithful chromosome segregation during meiosis depends on accurate recombination and synapsis of homologous chromosomes. These processes are monitored in mammals by checkpoint mechanisms involving the meiotic HORMA-domain proteins HORMAD1 and HORMAD2, which bind unsynapsed chromosome axes and promote activation of the DNA damage–response kinase ATR independently of DNA double-strand breaks (DSBs). However, no mechanism for axis recruitment of HORMAD1 or HORMAD2 had been demonstrated, nor had its role in checkpoint function been tested.

We establish that a putative HORMAD-interacting region—the closure motif (CM)—within the chromosome-axis component SYCP2 is selectively required for HORMAD2, but not HORMAD1, localization. Deletion of the SYCP2 CM disrupts SYCP2–HORMAD2 complexes and prevents HORMAD2 axis binding without affecting axis assembly or recombination. Consequently, ATR accumulation and signaling on unsynapsed axes are reduced, and the prophase checkpoint malfunctions in a sexually dimorphic manner—causing aberrant elimination of synapsis-proficient spermatocytes and persistence of asynaptic oocytes. The phenotypes of SYCP2-CM–deficient and HORMAD2-null mice are indistinguishable, establishing the requirement for HORMAD2 axis recruitment in synapsis surveillance.

We propose that axial recruitment generates a HORMAD2 scaffold that drives clustering-mediated ATR network activation independently of DSBs, thereby linking chromosome-axis architecture to synapsis quality control in mammalian meiosis.

## Introduction

Orderly segregation of chromosomes during meiosis requires that homologous copies of chromosomes (homologs) pair and form physical links with one another in meiotic prophase. In most taxa, including mammals, these physical linkages depend on inter-homolog crossovers generated by homologous recombination (reviewed in ^1^). Meiotic homologous recombination initiates with multiple programmed DNA double-stranded breaks (DSBs) along the proteinaceous axis of each chromosome. Axes assemble from cohesins and filamentous chromosome core proteins (SYCP2/SYCP3 in mammals, Red1 in budding yeast), and control both DSB formation and repair (reviewed in ^2^). DSB processing produces resected single-stranded DNA (ssDNA) ends that invade homologs, promoting synapsis of homolog axes and assembly of synaptonemal complexes (SCs). SCs, in turn, facilitate recombination-mediated repair of DSBs, yielding mostly non-crossover products and at least one crossover per homolog pair to ensure chromosome segregation in the first meiotic division (reviewed in ^1,2^).

Genome integrity is safeguarded by prophase checkpoints that eliminate meiocytes with persistent asynapsis and unrepaired DSBs (reviewed in ^3,4^). The DNA damage response (DDR) kinase ATR and auxiliary factors (e.g., ATRIP, TOPBP1, BRCA1, MDC1) are central to meiotic prophase checkpoints in mammals ^5–12^. ATR is activated by single-stranded DNA ends resulting from DSBs ^5,6^ and shows high activity on chromosome axes and chromatin in unsynapsed regions ^9,13,14^, where unrepaired DSBs persist longer than in synapsed regions. Elevated chromatin-associated ATR activity leads to meiotic silencing of unsynapsed chromatin (MSUC)^7,9,13,15–17^, which cooperates with canonical ATR signaling from DSBs in meiotic surveillance mechanisms for synapsis and recombination.

In males, restriction of MSUC to the heterologous unsynapsed X and Y chromosomes underlies meiotic sex chromosome inactivation (MSCI) and sex body formation during the pachytene stage of prophase ^18^. MSCI is central to spermatocyte survival beyond pachytene ^18–21^. It has been proposed either to prevent apoptosis-inducing gene expression from sex chromosomes ^17–20^ or to sequester and suppress persistent DDR signaling from autosomes ^21^. Conversely, failure to synapse homologs or repair DSBs on autosomes titrates ATR away from sex chromosomes, resulting in MSCI failure and persistent autosomal DDR signaling, either of which may trigger spermatocyte elimination ^18,21–23^. Whereas MSUC also operates in females ^24–26^, MSCI does not have a distinguished role as the two X chromosomes normally synapse. Instead, elimination of recombination-defective oocytes is triggered via persistent DDR signaling from unrepaired DSBs, ATR signaling from unsynapsed chromosomes, and/or resultant suppression of *ad hoc* sets of essential genes by MSUC ^12,24,27–31^. Thus, enrichment of ATR activity on unsynapsed chromosome sections is a common feature in meiotic prophase checkpoints in both sexes, yet, the underlying mechanisms remain incompletely understood.

Asynapsis is monitored by two meiotic HORMA (Hop1, Rev7, MAD2) domain proteins, HORMAD1 and HORMAD2, that preferentially associate with unsynapsed axes in mammalian meiocytes (Supplementary Fig. 1) ^32,33^. Whereas HORMAD1 has HORMAD2-independent roles in promoting SC and DSB formation ^34–40^, HORMAD1 and HORMAD2 collaborate in meiotic prophase checkpoints, a function proposed to involve enhancement of HORMAD2 binding to chromosomes by HORMAD1 ^35^. Via a poorly understood mechanism, HORMAD1 and HORMAD2 promote axis-binding of BRCA1 and TOPBP1, leading to ATR accumulation and activation on unsynapsed chromosome axes ^29,34–36,38^. In turn, axial ATR induces ATR spreading to associated chromatin loops by establishing a positive feedback circuit: ATR phosphorylates histone H2AX on serine 139 (γH2AX), followed by ATR recruitment to γH2AX-containing nucleosomes via MDC1-TOPBP1 complexes ^10^. This mechanism is thought to ultimately allow effective MSUC and DDR signaling on unsynapsed chromosomes.

A key untested tenet of the synapsis surveillance model is that ATR activation, axial DDR signaling, and MSUC are enabled by axis-bound HORMAD1 and HORMAD2, not by soluble HORMAD1 or HORMAD2. Axis-bound HORMADs are thought to enable ATR activation even in the absence of DSBs ^28,29^. However, it remains unclear how axis binding supports ATR activation and whether unbound HORMADs play a role in meiosis. The biological functions and mechanisms of HORMAD1 and HORMAD2 binding to chromosome axes are poorly understood, in part because genetic tools specifically disrupting HORMAD1 and HORMAD2 chromosomal localization in mammals have been lacking.

One way meiotic HORMA-domain proteins form protein complexes involves a “safety belt” mechanism, first described in the spindle assembly checkpoint protein MAD2 ^41,42^. In this mechanism, a ligand uses a closure motif (CM) to bind to an exposed surface on the N-terminal core of the HORMA domain (AlphaFold 3 predictions, amino acids 16-180 in HORMAD1 and 20-184 in HORMAD2). The ligand is then encircled by the C-terminal section of the HORMA domain (AlphaFold 3 predictions, amino acids 181-235 in HORMAD1 and 185-240 in HORMAD2), which folds back onto the N-terminal core like a safety belt (Supplementary Fig. 2) ^29,43,44^. This “closed” HORMA-domain conformation is thought to provide stable ligand binding that is regulatable by reversible opening and closing of the safety belt. Interactions of meiotic HORMA domain proteins with one another, cohesins, and Red1/SYCP2-related proteins have been proposed to enable assembly of HORMADs on chromosome axes in meiosis in diverse taxa ^43,45–48^. Potential CMs have been identified in the C-termini of HORMAD1 and HORMAD2 and in the central disordered region of SYCP2 ^43,45^. These CMs interact with HORMA domains of HORMAD1 and HORMAD2 in *in vitro* or exogenous interaction assays ^29,43,45^. Yet, their *in vivo* importance has not been reported, and the mechanism of HORMAD1 and HORMAD2 recruitment to chromosomes remains unclear.

## Results

### Predicted SYCP2 CM is required for male fertility but not axis and SC formation

To investigate the mechanism and function of axial HORMAD1 and HORMAD2 recruitment *in vivo,* we removed the postulated CM from SYCP2 (residues 395-414) by deleting the 16th exon of *Sycp2* (encoding residues 395-419) in mice (Fig. 1a-b, and Supplementary Fig. 3a-b, hereafter *Sycp2^e16^* allele). Levels of SYCP2 were comparable in testes of wild-type *Sycp2^+/+^* (hereafter WT) and *Sycp2^e16/e16^* mice (Supplementary Fig. 3c-d), indicating that the CM is not required for SYCP2 expression and stability.

**Figure 1.**
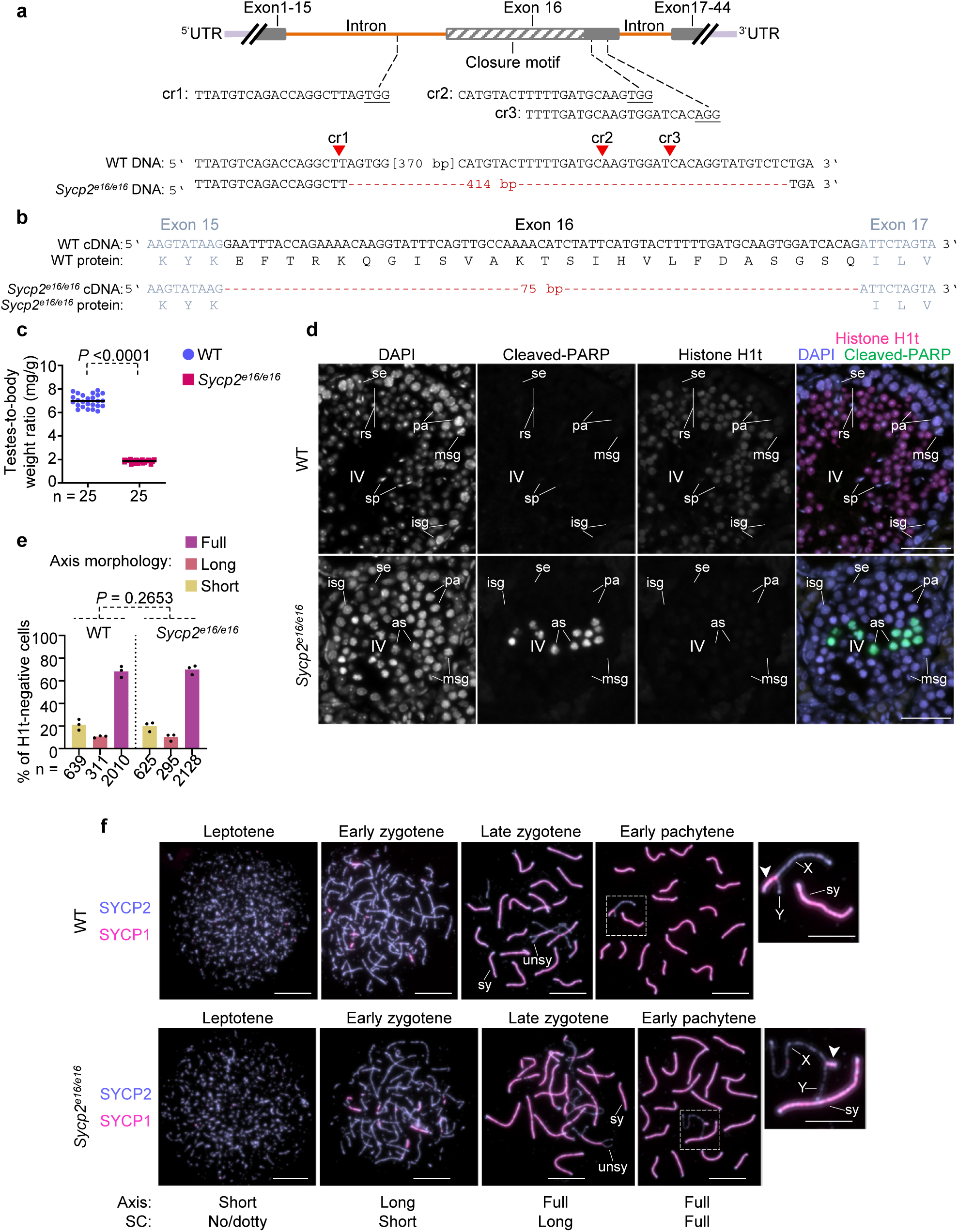
Defective spermatogenesis despite efficient axis formation in *Sycp2^e16/e16^* spermatocytes. **a** Targeting of *Sycp2* exon 16 by CRISPR/Cas9. The closure motif (diagonal stripes) is shown in the context of the *Sycp2* locus. Sequences of CRISPR RNAs (cr1-3, PAMs underlined), their predicted DNA cleavage sites (red arrowheads), and the resultant 414-bp deletion in genomic DNA are shown. **b** Sequences of *Sycp2* complementary DNA segments (cDNA) from testes. Inferred protein sequences are shown. **c** Testis-to-body weight ratios and their averages from n=25 mice of each genotype (*Sycp2^+/+^* (WT), 6.99 mg/g, *Sycp2^e16/e16^*, 1.86 mg/g). Statistical significance (P) was calculated by two-tailed unpaired t-test with Welch’s correction. **d** DNA staining (DAPI) and immunostaining of histone H1t (marker of mid pachytene-to-spermatid stages) and cleaved PARP (marker of apoptosis) in cryosections of testes. Cleaved PARP-positive spermatocytes are observed in the pachytene layer of *Sycp2^e16/e16^*, corresponding to apoptosis at epithelial cycle stage IV (see ‘Methods’ for staging). Sertoli cells (se), intermediate spermatogonia (isg), mitotic intermediate spermatogonia (msg), pachytene (pa) spermatocytes, round spermatids (rs), and sperm (sp) are marked. Scale bars, 50 µm. **e** Quantification of axis morphology categories in histone H1t-negative spermatocytes (see ‘Methods’, **f** and Supplementary Fig. 3f); Axis morphology was assessed via immunostaining of SYCP2 and SYCP3 in nuclear spreads preparations. Graph shows percentages of spermatocytes in each axis category from 3 experiments. Total number of analyzed cells (n) and weighted averages (block bars) are indicated. An analysis of deviance using the likelihood-ratio test based on the chi-squared distribution indicated no significant difference in the proportions of cell populations, P = 0.2653. **f** Immunostained nuclear spreads of spermatocytes from adult mice. Morphology categories for axes (relevant to **e**) and SCs (detected by SYCP1 staining, relevant to Fig. 4a) characterize the prophase stages indicated above each image. Unsynapsed (unsy) and synapsed (sy) regions of autosomes are marked in late zygotene. X and Y chromosomes and the pseudoautosomal region (PAR; white arrowheads) are marked in enlarged insets of early pachytene images. Scale bars, 10 µm (cell) and 5 µm (insets). Source data are provided as a Source Data file.

*Sycp2^e16/e16^* showed no obvious somatic abnormalities, but reproductive functions were severely compromised. Whereas females exhibited normal fertility (Supplementary Table 1), male *Sycp2^e16/e16^* mice were infertile (no pups after 26 breeding weeks, n = 3 males). Consistent with male infertility, *Sycp2^e16/e16^* mice had much lower testis weight than age-matched WT mice (Fig. 1c).

Examination of testis cross-sections revealed a spermatogenic block in *Sycp2^e16/e16^* mice (Fig. 1d and Supplementary Fig. 3e). We found no post-meiotic cells in the seminiferous tubules of *Sycp2^e16/e16^* mice (Fig. 1d and Supplementary Fig. 3e, > 500 tubules). Furthermore, spermatocytes expressing high levels of histone H1t—a marker of cells at and beyond the mid-pachytene stage, normally abundant after stage IV of the seminiferous epithelial cycle in WT—were absent (Supplementary Fig. 3e; see ‘Methods’ for epithelial cycle staging ^49^). Correspondingly, detection of the apoptosis marker cleaved PARP revealed high levels of spermatocyte death at stage IV of the seminiferous epithelial cycle (Fig. 1d). These findings suggest elimination of spermatocytes at mid pachytene as the cause of infertility in male *Sycp2^e16/e16^* mice. Because defects in meiotic axes, SC formation, recombination, or MSCI cause spermatocyte apoptosis at a mid-pachytene–equivalent stage (reviewed in ^3,4^), we investigated these structures and processes in *Sycp2^e16/e16^* mice.

Immunofluorescence analysis of spermatocytes showed that the core axis components SYCP2 and SYCP3 co-assembled into axes with similar efficiency and timing in WT and *Sycp2^e16/e16^* mice (Fig. 1e-f and Supplementary Fig. 3f). Axes also assembled efficiently in oocytes of *Sycp2^e16/e16^* mice (Supplementary Fig. 4b). Furthermore, staining of the SC transverse filament SYCP1 showed that axes supported SC formation in *Sycp2^e16/e16^* mice. This was evidenced by fully synapsed autosomes and synapsis between short homologous segments, the pseudoautosomal regions (PARs), of the X and Y chromosomes in early-to-mid pachytene, the most advanced prophase stage reached in *Sycp2^e16/e16^* spermatocytes (Fig. 1f). Thus, SYCP2 CM was not required for axis assembly or for the scaffolding function of axes in SC formation, which allowed direct comparison of prophase stages in WT and *Sycp2^e16/e16^* spermatocytes based on axis morphology.

### SYCP2 CM is crucial for HORMAD2 recruitment to chromosomal axes

To uncover the role of the predicted CM in SYCP2, we compared the behavior of HORMAD1 and HORMAD2 in WT and *Sycp2^e16/e16^* meiocytes. Both HORMAD1 and HORMAD2 were detected on unsynapsed chromosome axes in nuclear spreads of WT spermatocytes and oocytes ^32^ (Fig 2a-c, Supplementary Fig. 4a-b). In contrast, whereas *Sycp2^e16/e16^* meiocytes exhibited efficient HORMAD1 accumulation on unsynapsed chromosome axes, HORMAD2 was absent from axes (Fig. 2a-c, Supplementary Fig. 4a-b). Immunostainings of testis sections nevertheless showed that HORMAD2 was present at similar levels in spermatocyte nuclei from WT and *Sycp2^e16/e16^* mice (Supplementary Fig. 4c). Furthermore, Triton X-100 protein extraction showed comparable HORMAD2 levels in the soluble, chromatin-depleted fractions of WT and *Sycp2^e16/e16^* testis extracts, but a marked reduction in the chromatin-enriched insoluble fraction of *Sycp2^e16/e16^* extracts (Fig 2d). These observations indicate that CM loss specifically impairs axial recruitment of HORMAD2 without noticeably reducing its overall cellular abundance.

**Figure 2.**
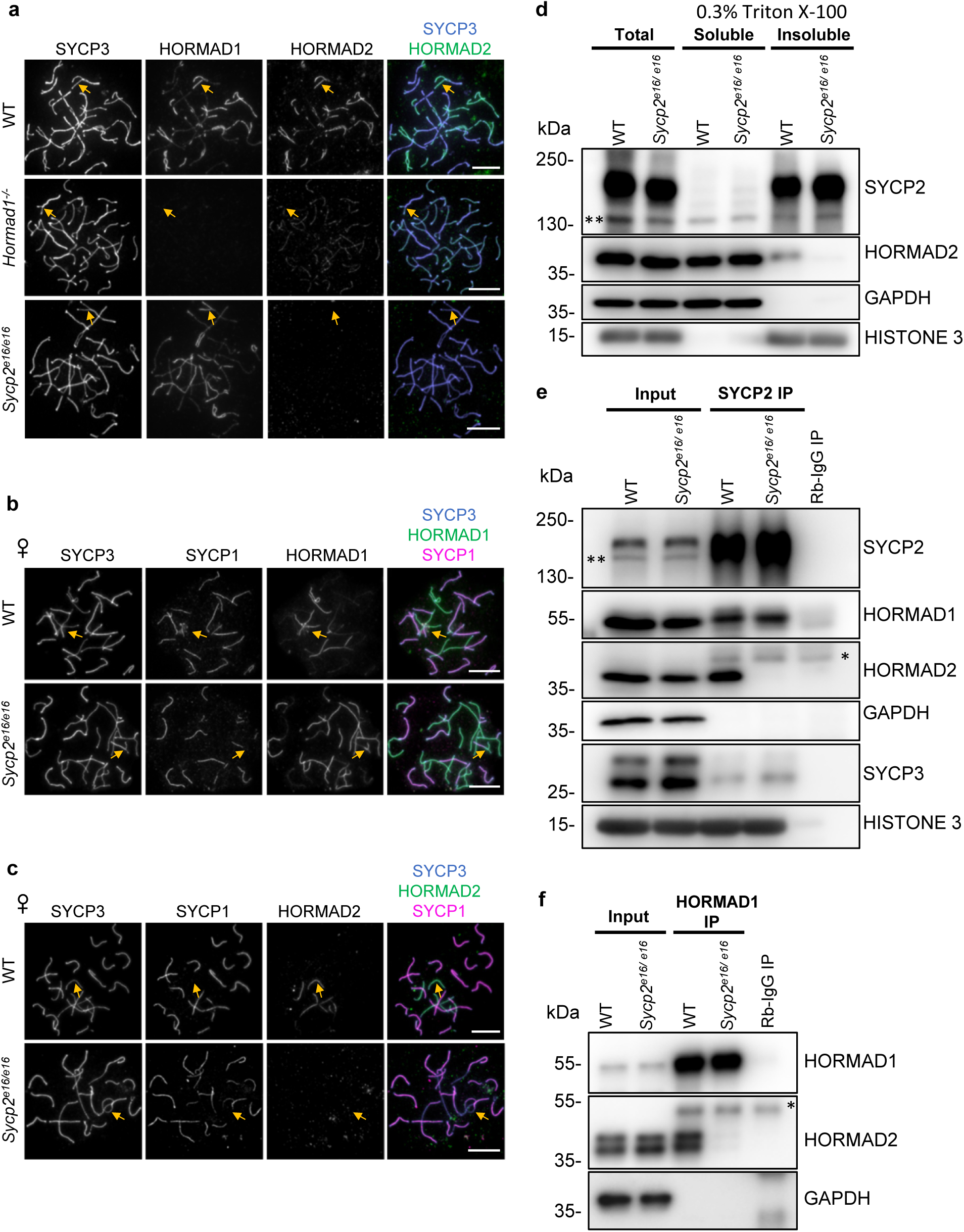
The 16^th^ exon of *Sycp2* is required for HORMAD2 recruitment to chromosomal axes. **a-c** Chromosome axis (SYCP3) was immunostained in combination with HORMAD1, HORMAD2 and/or SC marker SYCP1 in nuclear spreads of late zygotene spermatocytes from adult mice (**a**) or oocytes from fetuses at 17 days postcoitum (dpc) (b-c). Bars, 10 µm. Yellow arrows mark unsynapsed axes. **d-f** SDS-PAGE immunoblots of Total, Triton X-100-soluble (chromatin-poor) and -insoluble (chromatin-rich) protein extracts (**d**) or immunoprecipitation experiments of SYCP2 (**e**) or HORMAD1 (**f**) from testes of 13 dpp mice. SYCP2 and SYCP3 were resolved by 10% SDS–PAGE, whereas other proteins were resolved by 15% SDS–PAGE, and detected on separate blots in panels **d** and **e**. Unspecific bands in SYCP2 blots (** in **d, e**, see also Supplementary Fig. 5a) and heavy chain from precipitating antibodies in HORMAD2 blots (* in **e, f**) are marked. Depending on SDS-PAGE conditions HORMAD2 is detected as a single (**d, e,** 15% PAGE) or double band (**f,** 4-16% gradient). Source data are provided as a Source Data file.

This conclusion was further supported by SYCP2 immunoprecipitations from testis extracts, which showed that SYCP2 formed complexes with both HORMAD1 and HORMAD2 in WT, but only with HORMAD1 in *Sycp2^e16/e16^* mice (Fig. 2e, Supplementary Fig. 5a). Thus, the predicted CM in SYCP2 specifically promotes SYCP2-HORMAD2 complex formation, enabling HORMAD2 accumulation on unsynapsed chromosome axes.

Previous analysis found strong depletion of HORMAD2 from axes in *Hormad1^-/-^* mice and the presence of HORMAD1-HORMAD2 complexes in WT testis extracts ^35^. In addition, yeast two-hybrid assays showed that the HORMA domain of HORMAD1 interacted with both the HORMA domain of HORMAD2 and a putative CM in the C-terminus of HORMAD2 (Supplementary Fig. 5b-c), with the latter interaction also reconfirmed by an *in vitro* protein pull-down assay ^43^. These findings suggested that direct HORMAD1–HORMAD2 interactions play a key role in the axial recruitment of HORMAD2; hence, the observed loss of axial HORMAD2 in the presence of axis-associated HORMAD1 in *Sycp2^e16/e16^* meiocytes appears paradoxical. Notably, we found that HORMAD1 did not efficiently form complexes with HORMAD2 in *Sycp2^e16/e16^* testes (Fig. 2f). Thus, direct HORMAD1–HORMAD2 interactions observed in heterologous assays are insufficient for axial HORMAD2 recruitment and HORMAD1–HORMAD2 complex formation *in vivo*. Instead, our data suggest that the CM in SYCP2 provides the primary anchor for HORMAD2 axis binding, which in turn supports stable HORMAD1-HORMAD2 complex formation *in vivo*.

### SYCP2 CM loss does not markedly alter recombination dynamics

The specific loss of HORMAD2 from axes in *Sycp2^e16/e16^* meiocytes enabled us to test whether HORMAD2 axis association is essential for its meiotic functions. We therefore systematically compared meiotic phenotypes of *Hormad2^-/-^* and *Sycp2^e16/e16^* with WT. DSB formation and recombinationdynamics were monitoredby quantifying recombinationfoci between leptotene and early pachytene in nuclear spread spermatocytes (Fig. 3 and Supplementary Fig. 6). We detected foci of the recombinases RAD51 and DMC1, which bind resected ssDNA and promote homology search, and of RPA, which marks both early resected ssDNA and D-loops formed during strand invasions ^1,50^. To distinguish prophase stages, we used SYCP3 (to assess axis morphology as the primary staging criterion), IHO1 (present on unsynapsed axes in all prophase stages except early pachytene as shown in ^39,51^ and Supplementary Fig. 6 and 7), and histone H1t (present in and beyond mid pachytene ^52^) (for details see ‘Methods’).

**Figure 3.**
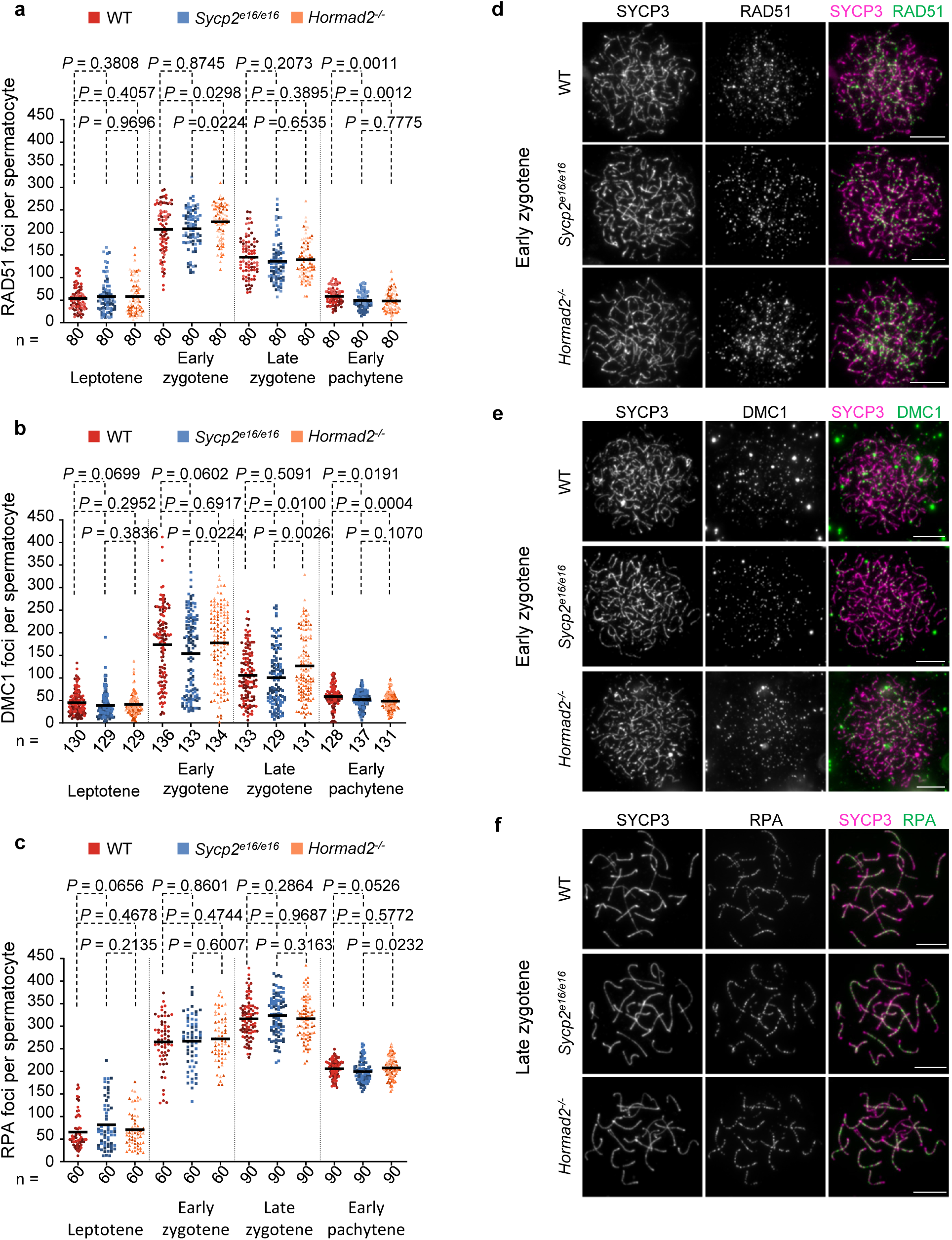
The 16^th^ exon of *Sycp2* is not essential for DSB repair. **a-c** Numbers of axis-associated DSB/recombination foci marked by RAD51 (**a**), DMC1 (**b**), and RPA (**c**) in spermatocytes at the indicated stages from adult mice. Focus quantification was performed in immunostained nuclear surface spreads which were staged using axis marker SYCP3 and stage markers IHO1 and histone H1t. Number of analyzed cells (n) and means (bars) are shown from four (**a**) or three (**b, c**) experimental replicates. Each replicate is shown with distinct shade of color. P values were calculated by unpaired t-test with Welch’s correction after pooling replicate data together. **d-f** Images show immunostained nuclear surface spreads of spermatocytes at stages where focus numbers of the corresponding recombination marker are the highest as shown in **a-c**. Images are shown with matched exposure and levelling for recombination markers, SYCP3 signal levels were adjusted to optimize visualization. Scale bars, 10 µm. See also Supplementary Fig. 6 for images of IHO1 and histone H1t channels corresponding to **d-f**, and images of spermatocytes at the other prophase stages corresponding to **a-c**. Source data are provided as a Source Data file.

Recombination dynamics were similar in all three genotypes, with median focus counts differing by less than 20% from WT levels, although some comparisons reached statistical significance of questionable biological relevance. These observations are consistent with a previous report showing that HORMAD2 loss has only a modest effect on DSB formation and repair kinetics during unperturbed meiosis ^35^.

### SYCP2 CM loss does not lead to major synapsis defects

Neither HORMAD2 ^35^ nor the SYCP2 CM (Fig. 1f, Supplementary Fig. 4a) is essential for SC formation, yet partial or delayed SC formation could contribute to the stage IV meiotic arrest in *Sycp2^e16/e16^* males. To test this possibility, and to identify potential SC defects that cannot be explained by HORMAD2 loss of function, we quantitatively assessed SC formation in nuclear spreads of *Sycp2^e16/e16^* spermatocytes (Fig. 4 and Supplementary Fig. 8). SC initiation and elongation dynamics were similar among WT, *Hormad2^-/-^* and *Sycp2^e16/e16^* spermatocytes between leptotene and early pachytene (Fig. 4a), but incomplete SCs and illegitimate chromosome associations were observed in both mutants (Fig. 4b-e and Supplementary Fig. 8). To quantify these defects, we focused on early pachytene, which in WT is marked by the completion of synapsis on all autosomes and the homologous PARs of X and Y chromosomes. Because synapsis completion could not be used as a staging criterion when evaluating synapsis defects, we instead used the unsynapsed-axis–binding protein IHO1, which disappears upon pachytene entry—independent of SC formation—for the duration of early pachytene ^39,51^. Fully formed continuous axes and absence of histone H1t characterize both late zygotene and early pachytene spermatocytes, with IHO1 detected only in late zygotene in WT. Across all three genotypes, IHO1 was present in similar proportions of spermatocytes that exhibited full-length axes and lacked histone H1t (Supplementary Fig. 7a-d), indicating comparable IHO1 dynamics and validating IHO1 loss as a marker for early pachytene.

**Figure 4.**
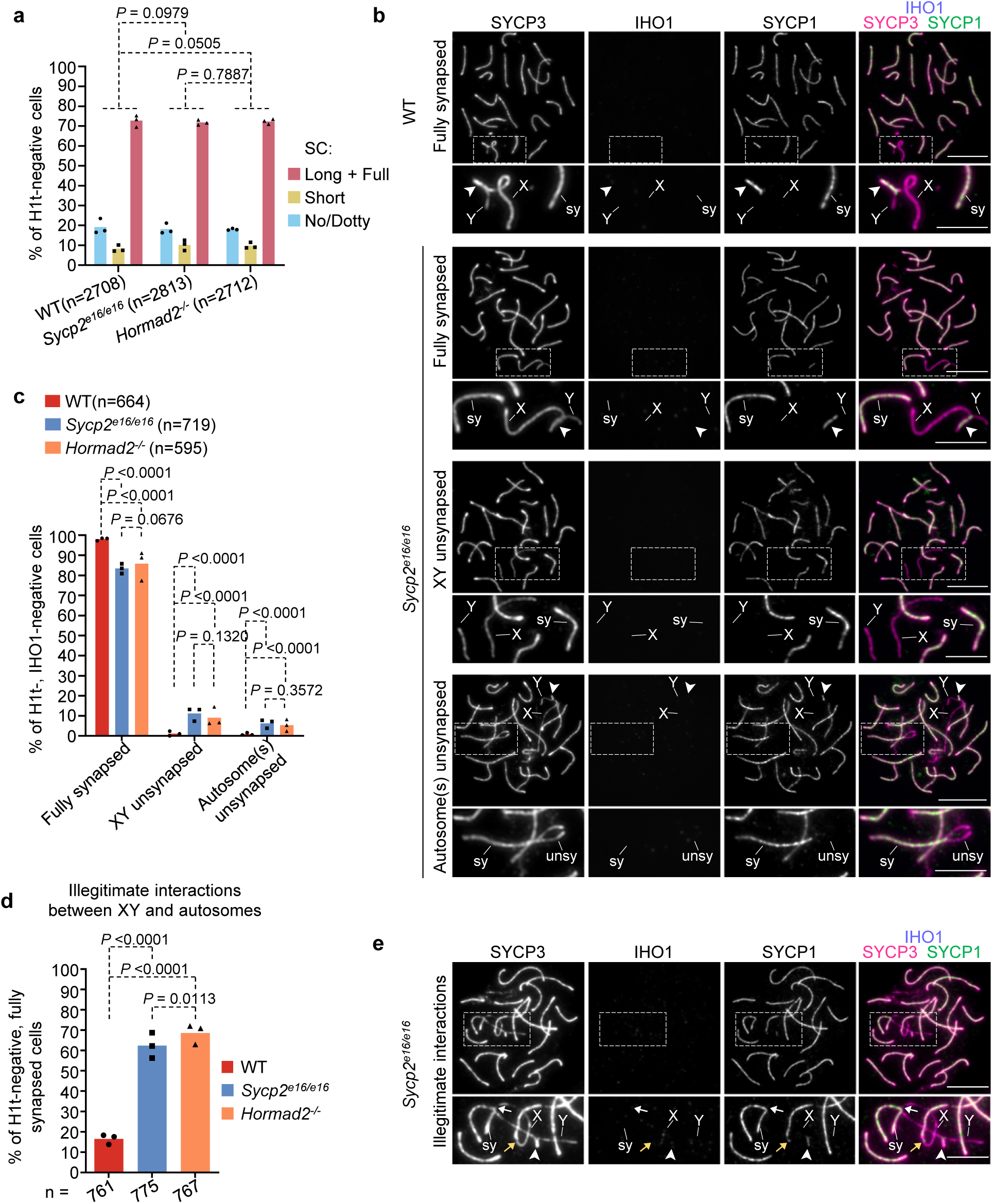
Most *Sycp2^e16/e16^* and *Hormad2^-/-^* spermatocytes are proficient in synapsis, but a minority exhibit mild synapsis defects. **a, c, d** Quantifications of synapsis formation (**a**) and SC defects (**c, d**) in spermatocytes from adult mice. Weighted averages of percentages of spermatocytes (block bars) and the total number of analyzed cells (n) from three experimental replicates are shown. The likelihood-ratio test was used to calculate the statistical difference in the proportions of spermatocytes in the indicated genotypes. **a** Degree of synapsis is quantified in histone H1t-negative spermatocytes (which corresponds to leptotene to early pachytene stages), corresponding to three categories of SC morphologies illustrated in Fig. 1f: no/dotty, short, and long+full SC. See ‘Methods’ for detailed explanation of categories. **c** Graph shows percentages of spermatocytes with indicated states of synapsis (illustrated in **b** by examples) in early pachytene stage as identified by fully formed axes and the absence of both histone H1t and IHO1 from cells. The “Autosomes unsynapsed” category represents spermatocytes where at least one autosome showed an unsynapsed axis fork. Cells displaying simultaneous XY and autosome asynapsis are counted in both “XY unsynapsed” and “Autosome(s) unsynapsed” categories. **d** Percentages of fully synapsed early pachytene cells that display illegitimate interactions between XY chromosomes and fully synapsed autosomes as illustrated by images in **e**. **b, e** Immunostained nuclear spreads of early pachytene spermatocytes illustrating synapsis categories quantified in **c**, **d**. Pseudoautosomal regions (PARs; white arrowheads in **b**, **e**), synapsed (sy; **b**, **e**) and unsynapsed (unsy; **b**) autosomal regions, and sites of illegitimate interactions between the X chromosome and autosomes (white arrow, axis entanglement; yellow arrow, end-to-end fusion) are indicated. Scale bars, 10 µm (cell) and 5 µm (inset). Source data are provided as a Source Data file.

Synapsis was therefore assessed in histone H1t-negative, IHO1-negative spermatocytes with fully formed axes (Fig. 4b-e, Supplementary Fig.8). Incomplete synapsis was detected in approximately 2% of WT, 17% of *Sycp2^e16/e16^*, and 14% of *Hormad2^-/-^* spermatocytes. Asynapsis of the XY PARs occurred about twice as frequently as incomplete autosomal synapsis in both *Sycp2^e16/e16^* (XY: 11%, autosomal :6%) and *Hormad2^-/-^* (XY: 9%, autosomal: 5%) cells. Thus, SC completion efficiency is only modestly reduced in *Sycp2^e16/e16^* spermatocytes, consistent with loss of HORMAD2 function, and is unlikely to explain the stage IV elimination of *Sycp2^e16/e16^* spermatocytes.

Beyond incomplete synapsis, we observed elevated illegitimate interactions between sex chromosomes and autosomes in *Sycp2^e16/e16^* (62%) and *Hormad2^-/-^* (69%) compared with WT (17%) early pachytene spermatocytes (Fig 4d-e, Supplementary Fig.8b). These aberrant associations included entanglements between X and autosomal axes or contacts between the non-PAR end of the X chromosome and termini of fully synapsed autosomes. Similar illegitimate interactions have been reported in mutants with disrupted sex-body formation and MSCI, potentially reflecting defective phase separation of sex chromosomes from transcriptionally active autosomes ^10,15,35,53^). Thus, the illegitimate sex chromosome-autosome associations might arise from impaired MSCI in *Sycp2^e16/e16^* meiocytes.

### SYCP2 CM is required for sex body formation and MSCI

To directly evaluate sex body formation in *Sycp2^e16/e16^* spermatocytes, we examined the localization of three key markers of HORMAD2-dependent ATR signaling that drives MSCI in early pachytene (Fig. 5 and Supplementary Fig. 9). These markers were (i) BRCA1, which promotes ATR recruitment and activation on unsynapsed sex-chromosome axes ^9^, (ii) ATR, which accumulates on the axes and, more diffusely, on associated chromatin loops ^9,54,55^, and (iii) γH2AX, which marks ATR activity and facilitates MSCI on sex-chromosome chromatin ^56^.

**Figure 5.**
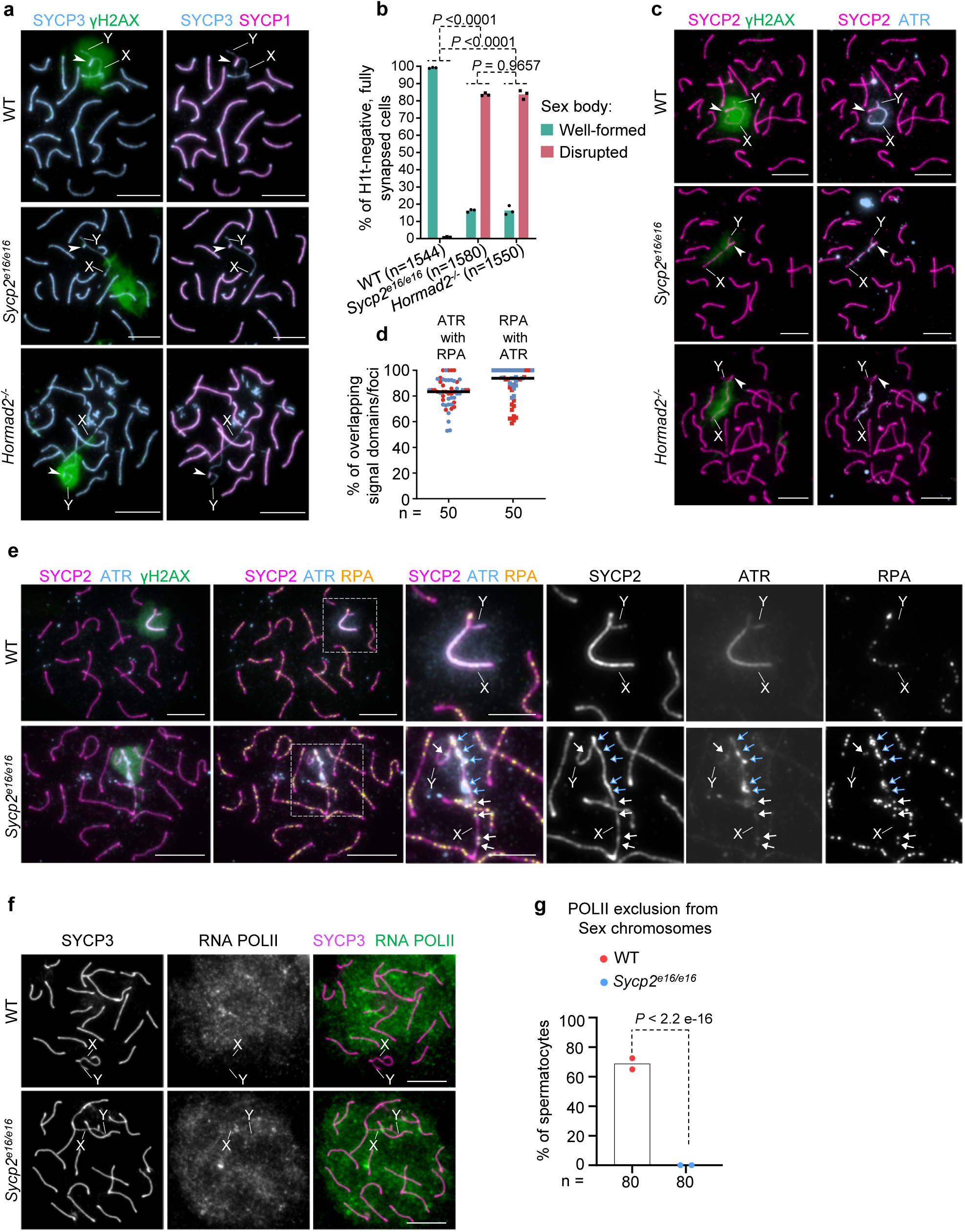
Mislocalization of silenced chromatin markers and ATR from sex chromosomes in *Sycp2^e16/e16^* spermatocytes. **a, c, e** Immunostained nuclear surface spreads of early pachytene spermatocytes from adult mice. Images with matched exposure and levelling are shown for γH2AX and ATR signals. X and Y chromosomes **(a, c, e)** and PAR (white arrowheads, **a** and **c**) are marked. In the enlarged inset of the *Sycp2^e16/e16^* spermatocyte image of panel **e**, blue and white arrows mark sites of sex-chromosome-axis-associated RPA foci within or outside of the γH2AX-chromatin domain, respectively. Scale bars, 10 µm (cell), 5 µm (insets, (**e**)). **b** Percentages of early spermatocytes with wild-type-looking (well-formed) or disrupted sex bodies. Total number of analyzed cells (n) and weighted averages (block bars) from three experimental replicates are shown. **d** Quantification of overlap between axis-associated RPA and ATR signals on sex chromosomes in *Sycp2^e16/e16^* spermatocytes. The numbers of analyzed cells (n) correspond to two experiments (25 cells each, differentiated by blue and red colors); medians (bars) are 83.30% (ATR colocalizing with RPA), and 93.80% (RPA colocalizing with ATR). **f** Immunostaining in nuclear spreads of pachytene spermatocytes of adult mice, showing RNA Pol II staining as a marker of chromatin regions with transcriptional activity. Bars, 10 µm. **g** Quantification of spermatocytes with clear exclusion of RNA Pol II from sex chromosomes. Total number of analyzed cells (n), means (block bars) from two experimental replicates and P value from Likelihood-ratio test (chi-squared distribution) are shown. Source data are provided as a Source Data file.

In nearly all wild-type early pachytene cells (99.2%, n = 1544), γH2AX was enriched in a well-defined chromatin domain encompassing the unsynapsed axes of the X and Y chromosomes along their entire length (Fig. 5a-c, Supplementary Fig. 9a). In contrast, most *Sycp2^e16/e16^* and *Hormad2^-/-^* ^35^ spermatocytes (83.7% for both genotypes) displayed spatially restricted or less distinct γH2AX-rich domains that did not fully coincide with X or Y axes (Fig. 5a-b). Consistent with these altered γH2AX patterns, ATR and BRCA1 accumulation on sex chromosomes was markedly reduced in *Sycp2^e16/e16^* spermatocytes (Fig. 5c-e, Supplementary Fig. 9a), mirroring reported *Hormad2^-/-^* phenotypes ^35,38^. Whereas both ATR and BRCA1 showed strong, continuous staining along X and Y axes in WT, their signals were weak and discontinuous in *Sycp2^e16/e16^* spermatocytes, with residual enrichment confined to regions marked by γH2AX (Fig. 5e, Supplementary Fig. 9a). In addition, the discontinuous accumulations of ATR and BRCA1, although not precisely co-localized with individual RPA foci or matching their signal intensities, broadly overlapped with RPA along sex-chromosome axes and followed a similar distribution pattern (Fig. 5d-e, Supplementary Fig. 9b-c).

Together, these observations indicate that *Sycp2^e16/e16^* and *Hormad2^-/-^* spermatocytesare similarly defective in the build-up of ATR activity on unsynapsed sex-chromosome axes, although low levels of BRCA1 and ATR accumulation still occur on the axes in response to unrepaired DSBs.

To test if disrupted sex body formation leads to defective MSCI, we analyzed the localization of RNA Pol II, as its exclusion from the sex body is a hallmark of silenced transcription on X and Y chromosomes ^57^. In WT, RNA Pol II appeared depleted from sex bodies in a significant fraction of mid pachytene cells. In contrast, RNA Pol II was consistently detected on the sex chromosomes in *Sycp2^e16/e16^* spermatocytes, suggesting disrupted MSCI (Fig. 5f-g).

To further test for MSCI failure in pachytene spermatocytes, we performed single-cell RNA sequencing (scRNA-seq) on whole-testis cell suspensions from adult WT, *Sycp2^e16/e16^*, and *Hormad2^-/-^* male mice (Fig. 6, Supplementary Figs. 10–13, Supplementary Table 2). Testis samples were obtained from two animals per genotype. WT samples were processed with and without dead cell removal; the former improved representation of early prophase spermatocytes through partial depletion of post-meiotic sperm populations, enabling comparison with mutant samples (Supplementary Fig. 12; see Methods).

**Figure 6.**
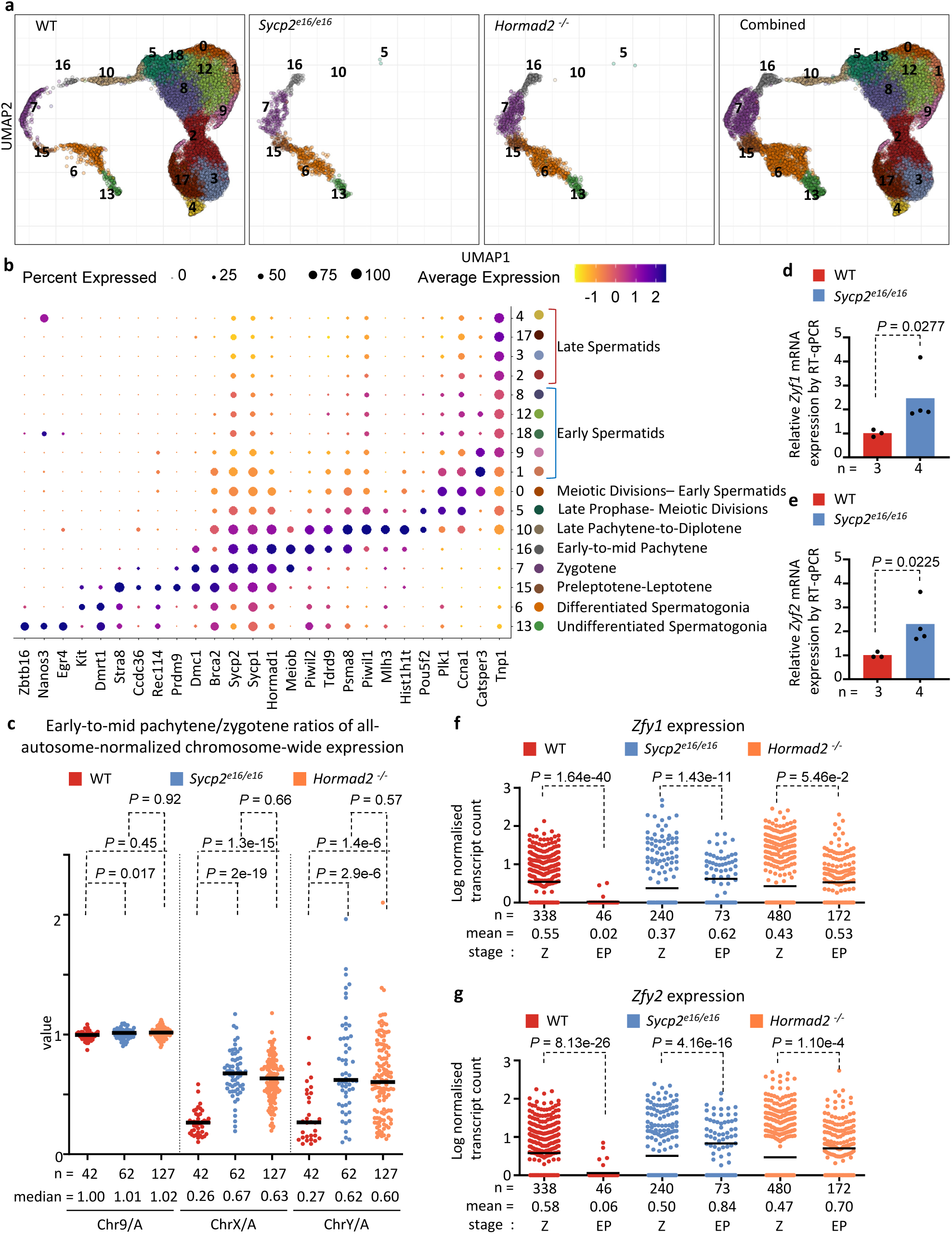
Single-Cell RNA-seq shows impaired sex chromosome silencing in *Hormad2^-/-^*and *Sycp2^e16/e16^* spermatocytes. **a** UMAP visualization of scRNA-seq data showing spermatogenic cell populations identified by supervised clustering and manual curation using stage-specific germline markers (see **b**, Supplementary Fig. 10; Supplementary Tables 3-5; ‘Methods’). Cluster numbers correspond to spermatogenic stages listed in **b**. UMAPs show pooled data from all experimental replicates for each genotype and combined data from all three genotypes. See Supplementary Fig. 12 for UMAPs of individual samples. **b** Expression dot plot of selected genes characterizing the spermatogenic cell clusters. The y-axis indicates cluster numbers and their dominant cell populations based on the expression of marker genes listed on the x-axis. The proportion of cells expressing each gene and the relative expression level (z-score ranging from -1 to 2) are represented by dot size and color, respectively. Clusters 6 and 7 each comprise two closely related subclusters. These subclusters displayed similar expression of dominant stage-defining germline marker genes (see in Supplementary Fig. 11), allowing their consolidation into single clusters corresponding to zygotene (cluster 6) and undifferentiated spermatogonia (cluster 7). **c** All-autosome-normalized chromosome-wide expression in early-to-mid pachytene cells relative to zygotene stage. Data points represent chromosome-wide expression in individual early-to-mid pachytene cells divided by the median normalized expression of the same chromosome in zygotene cells. Medians (bars) are shown*. P* values, Wilcoxon rank-sum test. **d-e** RT–qPCR quantification of *Zfy1* **(d)** and *Zfy2* **(e)** transcript levels in *Sycp2^e16/e16^* testes, shown as fold change relative to WT following normalization to ribosomal housekeeping genes. Total testicular RNA was isolated from 13-day-old juvenile mice. Bars, mean fold-change values from three (WT) or four (*Sycp2^e16/e16^*) biological replicates. *P* values, one-way ANOVA followed by Tukey’s HSD test. **f-g** Library size normalized transcript counts of *Zfy1* and *Zfy2* in zygotene (Z) and early-to-mid pachytene (EP) cells in scRNA-seq data. Individual cells and means (bars) are shown. *P* values, two-tailed Welch’s t test. All replicate samples per genotype were pooled for calculations; n=cell numbers in **c, f, g**. Source data are provided as a Source Data file.

Unsupervised clustering of expression profiles identified multiple transcriptionally distinct populations containing cells from all genotypes (Supplementary Fig. 10). However, this initial clustering did not resolve prophase stages with sufficient detail. To increase resolution, we first isolated germline clusters based on established germline markers (Supplementary Fig. 10b; Supplementary Tables 3 and 4), and then applied supervised clustering combined with manual curation using stage-specific prophase I marker genes (Fig. 6b and Supplementary Fig. 11; Supplementary Tables 3 and 5), guided by prior scRNA-seq studies^58,59^. This approach defined a continuous series of germ-cell populations spanning spermatogenesis, including distinct clusters corresponding to preleptotene–leptotene, zygotene, and early-to-mid pachytene spermatocytes (Fig. 6a, b).

Uniform manifold approximation and projections (UMAP) visualization revealed differential representation of the supervised clusters between WT and mutant testes (Fig. 6a and Supplementary Fig. 12). In WT, germ cells progressed from an early-to-mid pachytene (cluster 16) through a late pachytene transcriptional state (cluster 10) into spermatids. In contrast, *Sycp2^e16/e16^* and *Hormad2^-/-^* spermatocytes stalled after early-to-mid pachytene, consistent with the cytological stage IV arrest observed in these mutants (this study and ^35^).

We evaluated MSCI by quantifying chromosome-wide expression from our scRNA-seq data. For each cell, expression from X and Y chromosomes, as well as from individual autosomes was normalized to total autosomal expression (‘all-autosome-normalized’). Using these normalized values, we compared chromosome-specific expression in individual early-to-mid pachytene spermatocytes to the median normalized expression of the same chromosome in preleptotene-leptotene or zygotene cells (Fig. 6c and Supplementary Fig. 13).

All-autosome-normalized expression remained stable for representative autosomes of different lengths (chromosome 1 [long], chromosomes 9 and 10 [intermediate], and chromosome 19 [short]) as cells progressed from preleptotene-leptotene and zygotene to pachytene in WT, *Sycp2^e16/e16^*, and *Hormad2^-/-^*. Median expression ratios between stages ranged from 0.95 to 1.09. In WT spermatocytes, X- and Y-linked transcripts were reduced by 3.8-fold and 3.7-fold, respectively, in early-to-mid pachytene compared to earlier prophase stages, consistent with the onset of MSCI at pachytene entry. In contrast, sex-chromosomal expression ratios between early-to-mid pachytene and zygotene were significantly higher in both *Sycp2^e16/e16^* and *Hormad2^-/-^*spermatocytes, indicating that transcriptional silencing in early-to-mid pachytene is 2.4-2.6-fold less efficient for the X chromosome and ∼2.3-fold less efficient for the Y chromosome compared with WT. These data indicate a comparable, though incomplete, impairment of MSCI in both mutants, consistent with the partial cytological loss of γH2AX from sex chromosomes.

Failure of MSCI has been proposed to trigger spermatocyte elimination through the toxic overexpression of a subset of sex-chromosome-linked genes, including the Y-linked *Zfy1* and *Zfy2*^19^. Using quantitative real-time PCR, we detected more than twofold higher expression of *Zfy1* and *Zfy2* in testes from *Sycp2^e16/e16^* compared with WT mice at 13 days postpartum (dpp), when most spermatocytes in the first meiotic wave had reached early pachytene (Fig. 6d-e). Consistent with these results, scRNA-seq data showed that whereas *Zfy1* and *Zfy2* expression significantly dropped in WT spermatocytes upon progression from zygotene to early-to-mid pachytene (mean decrease: 9.8-to-22.4-fold), expression persisted at comparable overall levels in *Sycp2^e16/e16^* and *Hormad2^-/-^* spermatocytes (mean change: 1.2-to-1.7-fold) (Fig. 6f-g). Together, these data indicate MSCI impairment in both *Sycp2^e16/e16^* and *Hormad2^-/-^* spermatocytes, likely explaining mid-pachytene spermatocyte apoptosis and male infertility in both mutants.

### SYCP2 CM is crucial for the synapsis checkpoint in oocytes

Having established that the SYCP2 CM is required for HORMAD2-dependent checkpoint signaling and MSCI in spermatocytes, we also investigated its role in meiotic quality control in oocytes. Quality control of recombination and synapsis in oocytes is thought to depend on two genetically separable, yet interconnected, signaling pathways (^60,61^ and reviewed in ^3,4^). On one hand, unrepaired DSBs likely elicit a DNA damage response(DDR), which may involve ATM, ATR, and/or DNA-PK kinases. On the other hand, unsynapsed chromosome axes initiate BRCA1-dependent ATR signaling, which requires the synergistic functions of HORMAD1 and HORMAD2 ^12,24,28,29,34–38^. Both pathways contribute to the elimination of oocytes with recombination or synapsis defects at, or soon after, diplotene exit during the perinatal period in mice ^60^.

Although HORMAD2 has been suggested to delay DSB repair in unsynapsed regions ^27^, it is not essential for DDR signaling from unrepaired DSBs ^35^. This likely explains the dispensability of HORMAD2 for elimination of *Dmc1^-/-^* oocytes ^35^, which fail in DNA strand invasion and therefore accumulate both unrepaired DSBs and extensive asynapsis ^62,63^. In contrast, HORMAD2 is essential for efficient BRCA1- and ATR-mediated DDR signaling from asynaptic axes when DSBs are scarce ^35,38^, explaining the requirement for HORMAD2 in perinatal elimination of *Spo11^-/-^*oocytes, which display extensive asynapsis due to the absence of programmed DSBs ^64,65^.

We tested whether loss of the SYCP2 CM, and the consequent failure of axial HORMAD2 recruitment, results in checkpoint defects similar to those previously reported for *Hormad2^-/-^* mice. To this end, we quantified oocyte numbers in 6 week-old females carrying the *Sycp2^e16/e16^* mutation in combination with either *Dmc1^-/-^* ^62^, which disrupts DSB repair, or *Spo11^-/-^* ^64^ and *Iho1^-/-^* ^39^, which abolish programmed DSB formation and cause widespread asynapsis (Fig. 7a-b).

**Figure 7.**
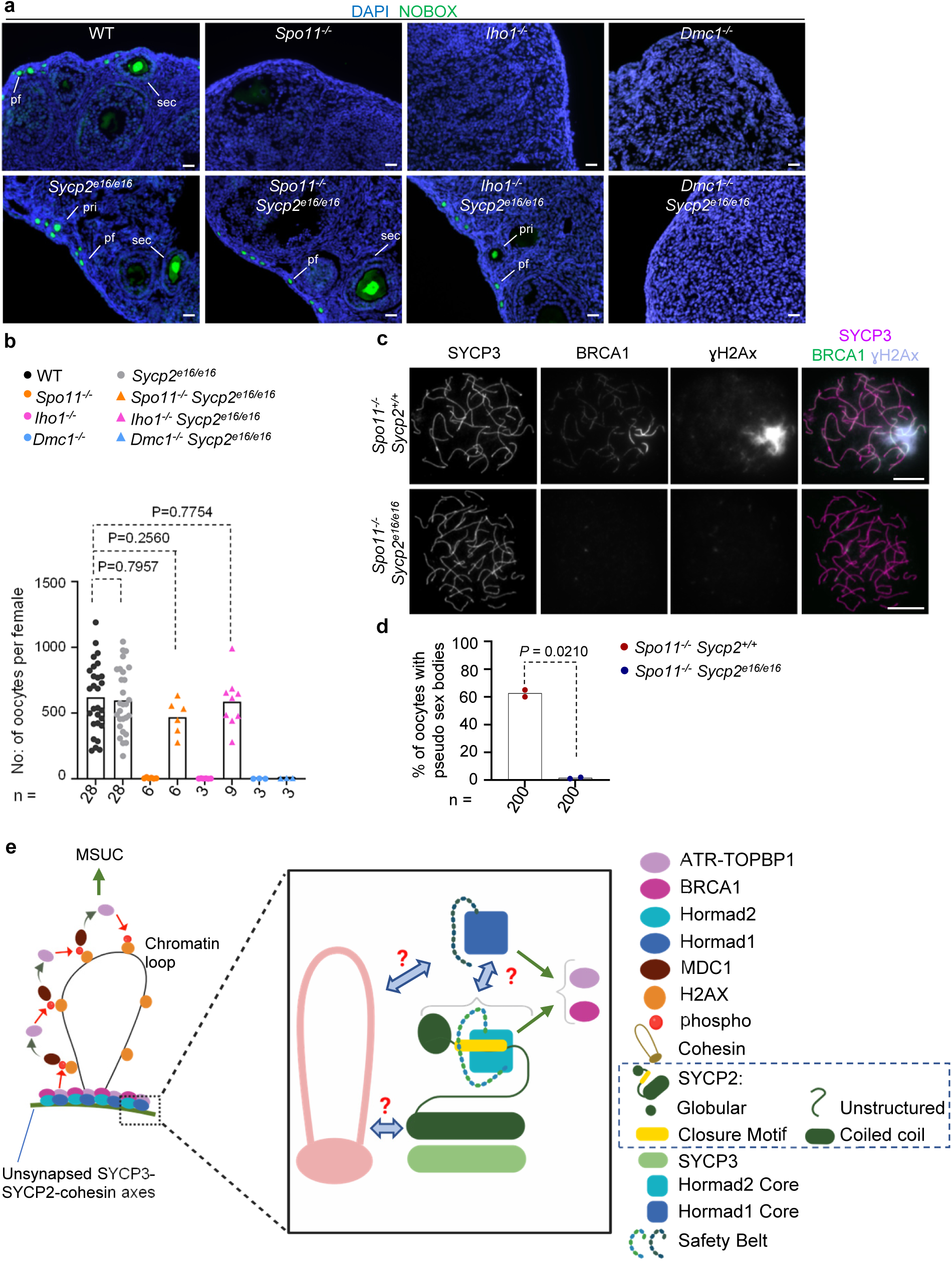
Axial HORMAD2 is essential for DSB-independent synapsis checkpoint signaling and regulates DNA repair turnover in oocytes. **a** DNA was labeled by DAPI and NOBOX (postnatal oocyte nucleus marker) immunostained in cryosections of ovaries from 6-week-old mice. Primordial (pf), primary (pri) and secondary (sec) follicles are indicated. Bars, 25 µm. **b** Quantification of oocyte numbers in ovarian sections from 6-week-old females of indicated genotypes. Data points represent sums of oocytes from every sixth section of both ovaries per mouse. Numbers of analyzed mice (n), means (bars) of oocyte numbers, and P values from two-tailed Welch t-test are shown. **c** Chromosome axis (SYCP3), BRCA1 and ɣH2Ax were detected by immunostaining in surface spreads of oocytes from one day old mice. Bars, 10 µm. **d** Quantification of pseudo-sex body formation in pachytene-early diplotene stage oocytes (as indicated by fully formed axes) collected from mice at 1dpp. Means (bar height) of two biological replicates, the numbers of assessed oocytes (n), and P values from two-tailed Welch t-test are shown. **e** Model for HORMAD1 and HORMAD2 recruitment to unsynapsed chromosome axes leading to ATR activation and MSUC. Green arrows indicate promoting effects; red arrows indicate phosphorylation; blue double-headed arrows indicate potential physical interactions between axial proteins. The model proposes undefined physical contacts between the SYCP2–SYCP3 complex and cohesins, enabling axis assembly, and between HORMAD1 and either cohesins or SYCP2, facilitating axial HORMAD1 recruitment. The HORMAD2 safety belt captures the closure motif (CM) encoded by exon 16 of *Sycp2*, providing the primary mechanism for HORMAD2 recruitment to axes. Axial HORMAD1 and HORMAD2 together promote recruitment of BRCA1, TOPBP1, and ATR, leading to phosphorylation of histone H2AX on chromatin flanking asynaptic axes (this study and ^34–38^). MDC1 binding to γH2AX recruits additional TOPBP1–ATR complexes, promoting spreading of ATR activity into chromatin loops ^10^. This cascade results in meiotic silencing of unsynapsed chromatin (MSUC) and sustained ATR signaling that underlies the synapsis checkpoint in both sexes. Source data are provided as a Source Data file.

Similar to *Hormad2^-/-^* mice, *Sycp2^e16/e16^* single mutants contained wild-type numbers of oocytes. Furthermore, *Sycp2^e16/e16^* rescued oocyte numbers to near wild-type levels in *Spo11^-/-^* and *Iho1^-/-^*backgrounds, but oocytes were efficiently eliminated in *Sycp2^e16/e16^ Dmc1^-/-^* double mutants. These results show that *Sycp2^e16/e16^* and *Hormad2^-/-^* genotypes comparably disable the synapsis checkpoint without disrupting DDR-mediated elimination of oocytes in response to persistent DSBs.

We next examined the mechanism underlying the synapsis checkpoint defect in *Sycp2^e16/e16^* oocytes (Fig. 7c-d). In the DSB-deficient *Spo11^-/-^* oocytes, BRCA1 and ATR accumulate on unsynapsed axes, enabling γH2AX accumulation both along these axes and within distinct chromatin domains encompassing subsets of unsynapsed chromosomes ^12,28,34^. Spermatocytes of meiotic DSB-deficient mutants also exhibit large γH2AX domains that are transcriptionally silenced, resembling sex bodies ^18^. However, these domains are not restricted to sex chromosomes and are therefore referred to as pseudo–sex bodies (PSBs) ^14,66^. Silencing of essential genes within PSBs, or ATR signaling emanating from PSBs and unsynapsed axes outside them, have been proposed as key drivers of perinatal oocyte elimination in *Spo11^-/-^* mice ^12,24,28^. Consistent with these hypotheses, loss of HORMAD2 diminishes perinatal oocyte elimination, BRCA1 and ATR accumulation in unsynapsed chromosomal regions, and PSB formation in the *Spo11^-/-^* background ^35,38^.

We therefore tested whether loss of the SYCP2 CM disrupts HORMAD2-dependent BRCA1 recruitment and ATR activation, as detected by γH2AX accumulation at unsynapsed chromatin regions, which would be expected to rescue *Spo11^-/-^* oocytes (Fig. 7c-d). In most perinatal *Spo11^-/-^* oocytes corresponding to late pachytene or diplotene stages, PSBs were present, and BRCA1 accumulated strongly within them, in addition to lower-level binding along unsynapsed axes outside PSBs. Notably, both BRCA1 and γH2AX were undetectable along unsynapsed axes, and PSBs were absent in *Sycp2^e16/e16^ Spo11^-/-^* oocytes. Together, these observations suggest that loss of axial HORMAD2 binding in the absence of the SYCP2 CM disrupts the meiotic synapsis checkpoint in oocytes by preventing HORMAD2-mediated recruitment of BRCA1 and BRCA1-dependent ATR activity to unsynapsed regions.

## Discussion

Whereas HORMAD1 and HORMAD2 have critical roles in safeguarding germline genome integrity during meiosis, it has not been reported how these proteins are recruited to chromosome axes, whether a predicted CM in the axis protein SYCP2 contributes to their recruitment, and whether axial localization is essential for HORMAD1 and HORMAD2 functions.

Here, we show that the region encoded by exon 16 of *Sycp2*, encompassing the predicted CM, is essential for HORMAD2—but not HORMAD1—recruitment to chromosome axes. *Sycp2^e16/e16^* mice phenocopy *Hormad2^-/-^* mice. Because axis assembly and SYCP2 abundance are highly similar in WT and *Sycp2^e16/e16^* mice, the most parsimonious interpretation is that loss of HORMAD2 from axes disrupts HORMAD2-dependent meiotic synapsis checkpoint mechanisms in *Sycp2^e16/e16^* mice. The off-axis pool of HORMAD2 therefore cannot substitute for its essential axial functions in promoting BRCA1 and ATR recruitment, which underpin synapsis checkpoint activity in both sexes.

### Mechanisms enabling HORMAD assembly on chromosome axes

Our *in vivo* data, together with previous interaction assays showing binding between HORMAD2 and the SYCP2 CM ^43^, strongly support the model that the SYCP2 CM acts as the primary recruiter of HORMAD2 to chromosome axes (Fig. 7E). Nevertheless, a paradox remains: HORMAD2 is lost from axes in *Sycp2^e16/e16^* meiocytes, yet low levels of both HORMAD1 and HORMAD2 associate with fragmented cohesin cores in *Sycp2^-/-^* spermatocytes ^46^. Thus, cohesins and/or HORMAD1 may recruit HORMAD2 through an SYCP2-independent mechanism that is normally suppressed when SYCP2 co-assembles with cohesins into intact axes.

Although HORMAD1 might also contact the exon 16-encoded CM in SYCP2, the high level of axial HORMAD1 in *Sycp2^e16/e16^* meiocytes demonstrates that alternative, efficient recruitment mechanisms exist. The weak HORMAD1 signal on cohesin cores in *Sycp2^-/-^* spermatocytes, however, indicates that SYCP2 contributes significantly to HORMAD1 loading ^46^. HORMAD1 might not rely on CM-binding for its axis association, or it may use as-yet-unidentified CMs in SYCP2 or cohesin subunits as redundant anchors.

Notably, the high axial accumulation of HORMAD1 in *Sycp2^e16/e16^* meiocytes is insufficient to recruit HORMAD2, despite earlier evidence that HORMAD1 enhances axial HORMAD2 accumulation ^35^. The HORMAD1 HORMA domain binds both the HORMA domain and the putative CM of HORMAD2 in heterologous assays, but these interactions are insufficient for stable *in vivo* HORMAD1–HORMAD2 complex formation or HORMAD2 recruitment in the absence of the SYCP2 CM. Axis binding may raise local HORMAD concentrations and enable direct HORMAD1–HORMAD2 contacts that stabilize HORMAD2 on axes. For example, HORMAD1 binding to the HORMAD2 C-terminal CM might prevent competition between the HORMAD2 and SYCP2 CMs for the HORMAD2 HORMA domain, reinforcing HORMAD2 association with the axis. Notably, a conceptually related regulatory logic has been proposed for the meiotic HORMA-domain protein ASY1 in *Arabidopsis thaliana*, where ASY1 binding to the closure motif of the axis protein ASY3 competes with intramolecular closure of the ASY1 HORMA domain, controlling ASY1 residence on chromosome axes ^67^.

### How does SYCP2 CM–mediated HORMAD2 recruitment enable checkpoint signaling?

HORMAD1 and HORMAD2 have essential, non-redundant roles in the synapsis checkpoint. We hypothesize that they act as axial anchors for the ATR-activating protein network, although direct physical interactions with the ATR network components remain undefined. Both HORMADs are required for efficient recruitment of BRCA1, TOPBP1, and ATR to unsynapsed chromosome axes ^34–38^, thereby establishing a foundation for ATR activity to spread into adjacent chromatin loops and drive MSUC ^10^(see models in Fig. 7e, Supplementary Fig. 1). A hierarchy exists among these factors: HORMAD1 and HORMAD2 act upstream of BRCA1 and TOPBP1, which in turn promote ATR recruitment and activation.

In *Sycp2^e16/e16^* meiocytes, ATR and BRCA1 are detected at low levels on some unsynapsed segments, indicating that axis-bound HORMAD1 can support limited BRCA1 and ATR recruitment even without HORMAD2. However, this residual activity is insufficient for a functional synapsis checkpoint, highlighting the importance of SYCP2 CM-mediated HORMAD2 recruitment in ATR activation. We envisage several not-mutually-exclusive mechanisms to explain this requirement:

#### Allosteric activation

SYCP2 CM binding may prime HORMAD2 for ATR-network engagement, either directly via allosteric activation or indirectly through enabling HORMAD1–HORMAD2 contacts that allosterically enhance HORMAD2’s signaling-competent conformation. In the latter scenario, HORMAD1 both stabilizes and activates HORMAD2 on axes while providing a secondary, lower-affinity docking site for ATR or its partners.

#### Clustering and scaffold formation

Axial HORMADs may serve as a structural scaffold for ATR signaling. In the canonical DNA damage response, ATR localizes to RPA-coated ssDNA ^6^ and undergoes autophosphorylation *in trans* among neighboring ATR molecules, which enables engagement with the ATR activator TOPBP1 ^68^. By analogy, ATR activation in meiosis might originate at DSB-associated ssDNA and then propagate along unsynapsed axes through sequential activation among adjacent ATR–auxiliary complexes stabilized by HORMADs.

HORMADs and TOPBP1 undergo ATR-dependent phosphorylation ^7,69,70^, suggesting positive feedback in the axial accumulation of HORMADs, ATR, and their partners. Indeed, partial dependence of BRCA1 and TOPBP1 recruitment on ATR has been reported, though not consistently across studies ^7,16^.

Axis recruitment of ATR and its cofactors likely facilitates not only propagation but also initiation of ATR signaling. Forced clustering of ATR and its partners can trigger ATR activation even in the absence of DNA damage ^70–73^. Axis-bound HORMAD1 and HORMAD2 may provide a repetitive binding lattice that concentrates ATR, BRCA1, and TOPBP1, promoting trans-activation analogous to ssDNA-induced clustering. Since HORMAD1 and HORMAD2 may engage distinct components of the ATR network, their simultaneous presence on axes could be required to assemble the full complement of ATR cofactors necessary for efficient activation.

Consistent with axis-driven clustering activating ATR, ATR activity—as indicated by γH2AX and S271-phosphorylated HORMAD2 ^7^—occurs along asynaptic axes largely devoid of DSBs in *Spo11^⁻/⁻^* meiocytes ^18,28^, which demonstrates that ATR can be activated independently of DNA breaks. Nonetheless, residual DSBs in these cells frequently coincide with PSBs, which are ATR-rich chromatin domains ^28,74^. BRCA1 (Fig. 7d) ^18^ and HORMADs ^32^ often hyper accumulate within PSBs compared with other unsynapsed regions, suggesting that initial ATR activation near DSBs promotes further recruitment of HORMADs and ATR cofactors in a self-amplifying manner. In *Hormad2^-/-^* and *Sycp2^e16/e16^* spermatocytes, a similar DSB-initiated mechanism may underlie residual ATR and BRCA1 build-up on sex-chromosome axes, resulting in localized ATR activation. Such positive feedback mechanisms may also provide a form of molecular memory of synapsis defects, which could be particularly relevant in female meiosis, where elimination of asynaptic oocytes mostly occurs after chromosomes desynapse at diplotene (see discussions in refs. ^24,28^).

Together, our data indicate that SYCP2 CM–mediated recruitment of HORMAD2 to unsynapsed axes enables efficient ATR activation. Reinforced by positive feedback loops within the ATR-activating network, this mechanism sustains high chromatin-bound ATR activity, which ensures elimination of asynaptic meiocytes—a key safeguard against aneuploidy in mammalian meiosis. More broadly, our findings indicate that closure motif–mediated recruitment of HORMA-domain proteins to the structures they monitor is a conserved regulatory logic shared between the spindle assembly checkpoint ^41,42,75^ and the meiotic synapsis checkpoint, despite their distinct cellular contexts and molecular targets.

## Methods

### Animal Experiments

Mice were kept in a C57BL/6J background (obtained from The Charles River Laboratories). Gonads were collected from mice after euthanasia. Experiments of spermatocytes and testes were carried out on samples from adult mice unless indicated otherwise. In the case of ovaries and oocytes, the experiments were performed in 6 weeks old and newborn mice. All animals were used and maintained in accordance with the German Animal Welfare legislation (‘‘Tierschutzgesetz’’). The mice were kept in the barrier facility in individually ventilated cages at 22–24◦C and 50–55% air humidity with a 14-h light/10-h dark cycle. The feed was a rat-mouse standard diet in the form of pellets. The stocking density in the used cage type IIL was a maximum of five mice. Hygiene monitoring was carried out according to FELASA guidelines. All procedures pertaining to animal experiments were approved by the Governmental IACUC (‘‘Landesdirektion Sachsen’’) and overseen by the animal ethics committee of the Technische Universität Dresden. The license number concerned with the present experiments with mice is TV A 8/2017 DD24.1-5131/395/10.

In addition to the newly generated *Sycp2* mutant mice, we used previously published mutants mouse strains *Hormad2* ^35^, *Dmc1* ^62^, *Iho1^deletion^* ^39^, and *Spo11* ^64^.

### Generation of *Sycp2*-mutant mice

The *Sycp2* mutant line was generated using the commercially available Alt-R™ CRISPR-Cas9 System (Integrated DNA technologies), targeting exon 16 of *Sycp2*. The web-based tools used for the prediction of target sequences were CHOPCHOP version 2 ^76,77^ and GT-Scan ^78^. For genome editing, a crRNA:tracrRNA duplex (crispr RNA:trans-activating crispr RNA) was prepared by mixing equimolar amounts of tracrRNA (an universal 67mer) and 6 different crRNAs (cr1: TTATGTCAGACCAGGCTTAG, cr2: CATGTACTTTTTGATGCAAG, cr3: TTTTGATGCAAGTGGATCAC, cr4: AACTGAAATACCTTGTTTTC, cr5: GCAGAATTTGCTTACAAACA, cr6: AGGTGTAAACAGTTGACCTA). The mixture was hybridized by heating to 95 °C for 5 min followed by slowly cooling to room temperature. A mixture of 1 µM tracrRNA:crRNA complex and 333 ng/ul Cas9-NLS (MPI-CBG, Dresden) in Cas9 Protein Buffer 1X (20 mM HEPES pH 7.5, 150 mM KCl) was microinjected into pronuclei and cytoplasm of fertilized oocytes from C57Bl/6NCrl mice, which were transferred to pseudopregnant mice. Microinjections and embryo transfer were performed by the CRISPR/Cas9 Facility at the Experimentelles Zentrum of the Medical Faculty Carl Gustav Carus (Dresden, Germany). Out of 96 transferred embryos, 17 pups were born and 9 of them had alterations in the targeted genomic locus. 3 out of the 9 mutants had a deletion encompassing the exon 16 of *Sycp2*, and had indistinguishable phenotypes in initial analysis, hence all subsequent analysis was carried out only with one of the lines. A founder female with a deletion of 414 bp was bred with a C57BL/6J male to establish the *Sycp2* mutant mouse line. All the experiments reported here were performed on samples from mice derived from F0 after at least five backcrosses.

### Genotyping of *Sycp2*-mutant mice

Tail biopsies were used to generate genomic DNA by overnight protease K digestion (0.1 mg/mL) at 55 °C in lysis buffer (200 mM NaCl, 100 mM Tris-HCl pH 8, 5 mM EDTA, 0.1% SDS). Following heat inactivation for 10 min at 95 °C, the genomic preparations where diluted and used for PCR. *Sycp2*-targeted F0 mice were genotyped by PCR amplification followed by agarose electrophoresis and DNA sequencing (Microsynth Seqlab). A combination of two primers, TGCAAGGCTTCTCTGTTCCT and TGCCTTTACAGTGGCTGCTT (Eurofins), was used to generate a PCR product of 1057 bp for wild-type and 643 bp for the *Sycp2* mutant mouse line.

### Sequencing of Sycp2 transcript

Total RNA was extracted from testes from 6-week males using the RNeasy Plus Kit (Qiagen). Complementary DNA (cDNA) was generated using the Invitrogen SuperScript™ III First-Strand Synthesis System (Invitrogen) and oligo dT (20) primers. cDNA was used as a template to amplify the *Sycp2* transcript from exon 11 to 19 (primers: AACTTGGTAAATGGCATTCTTGGTG and GGTTCCCTACTTCTTCGCGG); the product sizes were 713 bp for wild-type and 638 bp for the mutant allele. The samples were sequenced from exons 14 to 18 (primers: AATGGGAAGCTGTCACTGTAC and TCTTCGCGGTGTTGATGTG).

### Generation of antibodies against SYCP2, histone H1t, NOBOX, BRCA1 and DMC1

Polyclonal antibodies were raised against recombinant protein fragments of SYCP2, histone H1t, NOBOX, BRCA1 and DMC1. The immunogens corresponded to the following regions: SYCP2 C-terminal fragment (246 amino acids, Val¹²⁵⁵–Ala¹⁵⁰⁰), histone H1t C-terminal fragment (99 amino acids, Ser¹¹⁰–Lys²⁰⁸), NOBOX fragment (224 amino acids, Gly²⁸¹–Leu⁵⁰⁴), BRCA1 fragment (249 amino acids, Ser⁸¹⁰–Leu¹⁰⁵⁸), and DMC1 fragment (101 amino acids, Met¹–Val¹⁰¹). Coding sequences corresponding to these fragments were cloned into the pDEST17 bacterial expression vector to generate recombinant proteins containing an N-terminal 6×His tag.

Recombinant protein fragments were expressed in E. coli strains BL21(DE3)pLysS (SYCP2), BL21 tRNA (H1t), Rosetta2(DE3)pLysS (NOBOX), BL21 tRNA (BRCA1) and Rosetta2(DE3)pLysS (DMC1) and subsequently purified by Ni–Sepharose beads (Amersham, GE Healthcare) and SDS–PAGE (∼29 kDa). Purified proteins, diluted in PBS (phosphate-buffered saline) plus Freund’s complete adjuvant, were used for the first immunization of rabbits and guinea pigs. Subsequent immunizations used incomplete Freund’s adjuvant and homogenized SDS–PAGE gel fragments containing the purified proteins. SYCP2 C-terminal fragment, NOBOX, BRCA1 and DMC1 fragments coupled to NHS-activated Sepharose 4 Fast Flow beads (Amersham, GE Healthcare) were used to affinity-purify the corresponding polyclonal antibodies following standard procedures. Anti-histone H1t polyclonal antibodies were used as sera. Immunostaining of spermatocyte nuclear surface spreads and testis sections from wild-type mice and mice with stage IV spermatogenic arrest was used to validate the antibody.

### Protein extracts, immunoprecipitation and western blotting

#### Total protein extract preparation and western blot

Testes (10–15 mg of tissue) from 12–13 dpp animals were homogenized using a disposable tissue grinder pestle in lysis buffer at a final tissue concentration of 0.1 mg/µL. The lysis buffer contained 50 mM Tris-HCl (pH 7.5), 500 mM NaCl, 1% Triton X-100, and 5 mM MgCl₂, supplemented with protease and phosphatase inhibitors: 1 mM Phenylmethylsulfonyl Fluoride (PMSF), complete EDTA-free Protease Inhibitor Cocktail tablets (Roche, 11873580001), 0.5 mM Sodium orthovanadate, Phosphatase Inhibitor Cocktail 1 (Sigma, P2850), and Phosphatase Inhibitor Cocktail 2 (Sigma, P5726). Protease and phosphatase inhibitor cocktails were used at the concentrations recommended by the manufacturers. Testis homogenates were lysed using an overhead rotator for 60 minutes at 4°C in the presence of 1 µL benzonase (Merck Millipore) to digest DNA during the lysis of 10-30 mg of tissue. The homogenates were then mixed 1:1 with dilution buffer (50 mM Tris-HCl (pH 7.5), protease and phosphatase inhibitors as described above to reduce the concentration of salts and detergent. After dilution, total testis lysates were mixed 1:1 with 2×Laemmli sample buffer (containing 10% β-mercaptoethanol) and incubated for 7 minutes at 95°C. Samples were then run on 10%,15% (homemade) or gradient 4–16% SDS-PAGE gels (Bio-Rad) and transferred onto a PVDF membrane at 300mA for 2 hours. After transfer, membranes were blocked with 5% skimmed milk and incubated overnight at 4°C with primary antibodies under rotation in a 50 ml Falcon tube. After detection using the Amersham imager, band intensities were measured using ImageJ.

#### Fractionation of testis extracts based on Triton X-100 solubility

To prepare Triton X-100-soluble and -insoluble testis extract fractions, we followed the protocol previously described in^40^. Testes (10-15 mg) were homogenized using a disposable tissue grinder pestle in resuspension buffer at a final tissue concentration of 0.07 mg/µl (50mM Tris-HCl pH 7.5, 150mM NaCl and supplemented with phosphatase and protease inhibitors). Larger tissue fragments were further disrupted using a 200 µL pipette tip with a cut end. The homogenate was then divided into two portions: ¼ for total extract preparation and ¾ for fractionation. Both portions were mixed at a 1:1 ratio with 2× Triton X-100 buffer (50 mM Tris, pH 7.4; 150 mM NaCl; 0.6% Triton X-100; supplemented with protease and phosphatase inhibitors) and incubated at 4°C for 30 minutes with continuous mixing using an overhead rotator. After incubation, homogenates designated for total extracts were kept on ice until further processing. Meanwhile, samples intended for fractionation were centrifuged at 16,000g for 10 minutes at 4°C. The supernatants, representing the soluble fraction, were transferred to a fresh tube. The pellet, containing the insoluble fraction, was resuspended in an equal volume of the same buffer as the supernatant. Both unfractionated extracts and the Triton X-100 soluble and insoluble fractions were then mixed with 2× Lysis buffer (850 mM NaCl, 50 mM Tris-HCl, pH 7.5; 2% Triton X-100; 5 mM MgCl₂; supplemented with protease inhibitors and benzonase). After 1 hour of incubation, all samples were mixed with 2× Laemmli sample buffer (containing 10% β-Mercaptoethanol) at a 1:1 ratio and incubated at 95°C for 7 minutes. Finally, the samples were aliquoted and analyzed by Western blotting like described in the section above.

#### SYCP2 and HORMAD1 immunoprecipitation (IP)

Testes extracts from 12–13 dpp animals were prepared as described above for total protein extraction. Tissues were lysed by preparing homogenates in 0.1 mg tissue /µL lysis buffer concentration. Following lysis, samples were centrifuged at 16,000 g for 10 minutes at 4°C, and the supernatant was collected while avoiding debris. The lysate was diluted 1:1 with dilution buffer (the same as described above for total protein extraction), resulting in a final tissue concentration of 0.05 mg/µL. The diluted supernatant was used as the IP input sample. For each IP reaction, 40 µL of Protein A magnetic beads were washed twice with PBS (pH 7.4) containing 0.02% Tween-20. The beads were then resuspended in 600 µL of PBS (pH 7.4) with 0.02% Tween-20, and 2 µg of anti-SYCP2 rabbit antibody was added (or 2 µg of rabbit anti-HORMAD1 antibody for HORMAD1 IP). Beads were incubated for a minimum of 2 hours at 4°C with continuous rotation. After incubation, beads were washed once with 600 µL wash buffer (50 mM Tris-HCl (pH 7.5), 250 mM NaCl, 0.5% Triton X-100, protease and phosphatase inhibitors), resuspended in 40 µL wash buffer, and used immediately for immunoprecipitation. To initiate IP, the wash buffer was removed from the antibody-conjugated beads, and 200–250 µL of the input sample was added. Beads were resuspended and incubated for 1-1.5 h at 4°C with continuous rotation. After incubation, beads were washed three times with the wash buffer. Immunoprecipitated proteins were eluted by adding 40 µL of 1× SDS Laemmli buffer containing 5% β-mercaptoethanol to the beads and incubating for 10 minutes at room temperature. The eluate was then separated from the beads using a magnetic rack and boiled for 7 minutes at 95°C. The resulting samples were subsequently analyzed by Western blotting.

### Antibodies used for detection of western blots

The following primary antibodies were diluted in 5% BSA containing 0.05% Tween-20 and used for detection of western blots: guinea pig anti-HORMAD1 (1:3000, ^32^), guinea pig anti-SYCP2 (1:1000), guinea pig anti-HORMAD2 (1:2500, ^32^), rabbit anti-Histone H3 (1:200 000; Abcam, ab1791), mouse anti-GAPDH (1:1000; Santa Cruz Biotechnology, sc-32233), and chicken anti-SYCP3 (1:1000;^79^).

### Yeast two-hybrid (Y2H) assay

Yeast two-hybrid experiments with mouse proteins were performed as described previously with minor modifications ^39^ . Pairwise interactions were carried out in the Y2HGold Yeast strain (Cat. no. 630498, Clontech). To transform the Y2HGold yeast strain with bait and prey vectors, yeasts were grown in 2xYPDA medium overnight at 30°C, 200 rpm shaking. Afterward, yeast cells were diluted to 0.4 optical density (OD, measured at 600 nm) and incubated in 2x YPDA for 5 h at 30°C, 200 rpm shaking. Cells were harvested, washed with water and re-suspended in 2 mL of 100 mM lithium acetate (LiAc). 50 µL of this cell suspension was used for each transformation. Transformation mix included 1 µg of each vector (bait and prey), 60 µL of polyethylene glycol 50% (w/v in water), 9 µL of 1.0M LiAc, 12.5 µL of boiled single-strand DNA from salmon sperm (AM9680, Ambion), and water up to 90 µL in total. The transformation mix was incubated at 30°C for 30 min, and then at 42°C for 30 min for the heat shock. The transformation mix was removed following centrifugation at 1000 g for 10 min, and then cells were resuspended in water, and plated first on -Leu -Trp plates to allow selective growth of transformants. After 2-3 days of growth, the transformants were cultured overnight in –Leu –Trp minimal medium till 3-7 OD₆₀₀. The next day, cells were diluted to an OD₆₀₀ of 1 in water and 10µl spotted onto both -Leu -Trp and -Leu -Trp -Ade -His plates. Plates were incubated for 2–6 days to test for interactions. We followed the manufacturer’s instructions for media and plate preparation.

### Immunofluorescence microscopy

Nuclear surface spreading of meiocytes and their immunostaining was carried out according to earlier described protocols ^80^ with minor modification.

#### Preparation of spermatocyte spreads

Detunicated testes were minced with scalpel in PBS pH 7.4, and filtered through a 70 µm sieve. The suspension was mixed with hypotonic extraction buffer (30 mM Tris-HCl pH 8.2, 50 mM sucrose pH 8.2, 17 mM sodium citrate pH 8.2, 5 mM EDTA pH 8.2, 0.5 mM DTT) in 1:1 ratio and incubated for 6 min at room temperature. Following incubation, the cell suspension was diluted five times in PBS pH 7.4, and centrifuged for 5 min at 1000 *g*. The pellet was resuspended in 100 mM sucrose solution and filtered through a 20 µm sieve. The cell suspension was added onto diagnostic slides, along with 5-7 times the volume of filtered (0.2 μm) fixative (1% paraformaldehyde, 0.15% Triton X-100, 1 mM sodium borate pH 9.2). The slides were incubated for 60–90 min at room temperature in wet chambers. After the incubation period, the spreads were air-dried under a fume hood, which could take from 30 min to 1 hr. Once dry, the slides were washed in 0.4% Photo-Flo (Kodak) and distilled water, and air-dried at room temperature. For the quantification of axis morphology and synaptonemal complex (SC) categories, a variation of the method omitting the centrifugation steps was employed to preserve native proportions of distinct spermatogenic cell types. Specifically, the testes were minced in a minimal volume of PBS pH 7.4 (1 µl per each mg) and diluted five times with 100 mM sucrose and 0.25 mM DTT. The resulting cell suspension was sequentially passed through 70 µm and 20 µm sieves, and then diluted with an equal volume of 100 mM sucrose and 0.25 mM DTT. The fixation step was performed as previously described.

#### Preparation of oocyte spreads

To prepare nuclear surface spread oocytes, two ovaries from each mouse were incubated in 20 µL hypotonic extraction buffer at room temperature for 15 min (Hypotonic Extraction Buffer/HEB: 30 mM Tris-HCl, 17 mM Trisodium citrate dihydrate, 5 mM EDTA, 50 mM sucrose, 0.5 mM DTT, 0.5 mM PMSF, 1x Protease Inhibitor Cocktail). After incubation, HEB solution was removed and 16 µL of 100 mM sucrose in 5 mM sodium borate buffer pH 8.5 was added. Ovaries were punctured by hypodermic needles (27Gx1/2“) to release oocytes. Big pieces of tissue were removed. 9 µL of 65 mM sucrose in 5 mM sodium borate buffer pH 8.5 was added to the cell suspension and incubated for 3 min. After mixing, 1.5 µL of the cell suspension was added in a well containing 20 µL of fixative (1% paraformaldehyde, 1 mM borate buffer pH: 9.2, 0.15% Triton X-100) on an uncoated glass slide. Cells were fixed for 45 min in humid chambers, and then slides were air-dried. Upon completion of drying slides were washed with 0.4% Photo-Flo 200 solution (Kodak, MFR # 1464510) for 5 min and afterward they were rinsed with distilled water temperature before proceeding to immunostaining.

#### Immunostaining of nuclear surface spreads

For immunostaining nuclear surface spreads of meiocytes, slides were blocked for 1 h at room temperature. The following blocking solutions were used: (1) 5% Normal Goat Serum (NGS) in PBS-TT (PBS, 0.05% Triton X-100, 0.05% Tween 20); (2) 2.5% BSA (Bovine serum albumin) in PBS-TT. Details of the optimal blocking solution for each combination of antibodies are available upon request. After blocking, slides were incubated overnight or for 3 h at room temperature with following primary antibodies diluted in blocking solution: rabbit anti-SYCP2 (this study, 1:5000), guinea pig anti-SYCP2 (this study, 1:3000), chicken anti-SYCP3 (1:800) ^79^, mouse anti-SYCP3 (gift from R. Jessberger, 1:2), chicken anti-SYCP1 (1:600) ^81^ , rabbit anti-SYCP1 (ab15090, Abcam, 1:800), guinea pig anti-H1t (this study, 1:2500), guinea pig anti-H1t (1:20000)^81^, guinea pig anti-HORMAD2 (1:800)^32^, chicken anti-IHO1 (1:600)^39^, rabbit anti-IHO1 (1:2000)^39^, rabbit anti-RAD51 (ab176458, Abcam, 1:800), mouse anti-DMC1 (ab11054, Abcam, 1:150), guinea pig anti-DMC1 (this study, 1:250), rabbit anti-RPA32/RPA2 (ab76420, Abcam, 1:1000), mouse anti-γH2AX (05-636, Millipore, 1:6000), goat anti-ATR (sc-1887, Santa Cruz, 1:50). Slides were then washed three times with PBS-TT and incubated for 1 h at room temperature with goat, donkey or bovine secondary antibodies conjugated with Alexa Fluor™ (AF) 647 (A21245, A21450, A21449 Invitrogen/Thermo Fisher Scientific; 711-495-152, 706-605-148, Jackson ImmunoResearch), AF 594 (703-585-155, Jackson ImmunoResearch), AF 568 (A11036, A11075, A11031, A11057, A11041, Invitrogen/Thermo Fisher Scientific), AF 488 (A11034, A11073, A11029, A11055, A11039, Invitrogen/Thermo Fisher Scientific; 706-545-148, 703-545-155, Jackson ImmunoResearch), AF 405 (A31553, Invitrogen; ab175675, Abcam), 350 (A10035, Invitrogen/Thermo Fisher Scientific), DyLight™ 488 (711-485-152, Jackson ImmunoResearch), DyLight™ 405 (706-475-148, Jackson ImmunoResearch), or Rhodamine Red™-X (805-295-180, Jackson ImmunoResearch). Secondary antibodies were diluted in the corresponding blocking solution 1:600, with very few exceptions: 706-605-148, A10035, 711-485-152, and 805-295-180 diluted 1:300; A31553, diluted 1:200. Details for immunostaining with each combination of antibodies are available upon request. After incubation, Slides were washed 3 times with PBS and embedded in SlowFade™ Gold Antifade Mountant (Invitrogen). All incubations were performed in closed wet chambers.

#### Immunofluorescence in gonad sections

Mouse testes were dissected and tunica albuginea was gently pricked with a needle. Testes were fixed with 4% formaldehyde, 0.1% Triton-X in 100 mM Sodium Phosphate Buffer pH 7.4, for 1 h at room temperature. In the case of females, we assessed oocyte numbers in ovaries collected from young adults (6 weeks old). Ovaries were dissected together with surrounding fat and a piece of oviduct and then cleaned from surrounding tissue inside a PBS pH 7.4 droplet. Clean ovaries were fixed in 4% Paraformaldehyde in 100 mM Sodium Phosphate Buffer pH 7.4 for 20-30 min at room temperature. Following fixation, the gonads were washed three times with PBS pH 7.4 and incubated in a solution of 30% Sucrose and 0.02% sodium Azide overnight at 4 °C. Afterwards, gonads were placed into an embedding mold, frozen on dry ice in Tissue-Tek® O.C.T. Compound (Sakura Finetek Europe) and stored at -80°C until sectioned.

For the testes, 5 µm thick sections were cut onto diagnostic slides and air-dried. The sections were washed with PBS pH 7.4, 0,05% Tween 20, 0,05% Triton X-100 to eliminate the O.C.T. compound and immediately used for immunofluorescence staining. The blocking solution used was 5% NGS in PBS-TT. To detect apoptosis, rabbit anti-cleaved PARP (9544, Cell Signalling, 1:250) was used along anti-histone H1t, followed by DAPI (5 µg/ml) staining. Histone H1t immunostaining was performed sequentially in order to avoid cross-reactivity. To detect HORMAD2, guinea pig anti-HORMAD2 (1:500)^32^ was used, followed by DAPI staining. Testis sections were blocked and incubated as described for spreads, and embedded in SlowFade™ Gold Antifade Mountant with DAPI (Invitrogen).

In the case of ovaries, 8 µm thick sections were cut using a Leica CM1850 cryostat, and the sections were dried onto glass slides at room temperature. Slides were washed several times with distilled water to remove OCT and then air dried. The dried slides were blocked with 2.5% w/v BSA in PBS for 1 hour and then stained (see immunostaining of spreads). NOBOX was detected in the oocyte sections, and oocytes were counted in every tenth section to determine the total number of oocytes.

#### Staging of meiotic prophase in nuclear spreads

As previously described ^39,81^, the staging of nuclear spreads was performed by evaluating axis development through detection of SYCP3, in combination with SYCP1 (SC) and histone H1t. The first meiotic prophase can be subdivided into preleptotene, leptotene, early zygotene, late zygotene, early pachytene (which are all histone H1t-negative), late pachytene, diplotene and diakinesis (characterized by histone H1t presence). A punctate staining pattern of SYCP3 throughout the nucleus characterizes the preleptotene stage. Leptotene is characterized by short stretches of axes and no or dotty SC.

During early zygotene, the axes are long but fragmented. In wild-type spermatocytes, short stretches of the SC are already observable at this stage. Late zygotene is characterized by the complete formation of axes for all chromosomes. In wild-type spermatocytes, the SC is typically long but remains incomplete at this stage. Cells reach pachytene when SC formation is completed in wild-type. In pachytene oocytes, complete synapsis is observed for all chromosomes. However, in spermatocytes, full synapsis is limited to autosomes, while heterologous sex chromosomes only synapse within their pseudoautosomal regions (PARs). Additionally, early pachytene can be distinguished by IHO1 detection. IHO1 accumulates on unsynapsed axes in late zygotene, gets restricted to the PAR region during zygotene-to-pachytene transition, transiently disappears from chromosomes for the duration of early pachytene, and reappears on the unsynapsed axes of sex chromosomes from mid-pachytene as shown earlier ^39^. During early pachytene, histone H1t is either absent or present at low levels in spermatocytes. As pachytene progresses, histone H1t levels gradually increase and reach an intermediate level during mid-pachytene. In late pachytene, Histone H1t levels are significantly elevated. The diplotene stage is characterized by axis desynapsis and fragmentation of the SC, along with high levels of histone H1t levels in spermatocytes. While similar stages exist in oocytes, histone H1t cannot be utilized as a reliable marker for staging in oocytes.

#### Staging of mouse seminiferous tubule cross sections

Meiosis is initiated in cohorts of synchronous spermatogenic cells inside seminiferous tubules in a cyclic process. These cohorts migrate from the basal membrane towards the lumen as they mature, forming layers of pre-meiotic, meiotic, and post-meiotic cells populations, whose distinct associations distinguishes 12 separate stages of the seminiferous epithelial cycle ^49^. The criteria we used to stage the epithelial cycle of mouse seminiferous tubules was described earlier ^81,82^. DAPI, which labels DNA, was used to detect nuclear morphology and chromatin condensation patterns. Histone H1t detection aid in staging, as it is a marker of spermatocytes after mid pachytene. **Stages I-IV:** the basal layer consists of large spermatogonia A and intermediate spermatogonia, which increase in numbers as tubules progress to stage IV. Chromatin in intermediate spermatogonia has a low DAPI intensity with occasional small and bright heterochromatic regions. The second layer contains early pachytene cells which are negative or very low for histone H1t, but histone H1t signal start to be noticeable in mid pachytene cells by the end of stage IV. The intermediate layer contains early round spermatids and advanced elongated spermatids are observed in the luminal layer. Stage IV can feature intermediate spermatogonia undergoing mitosis. **Stages V-VI:** the basal layer contains several spermatogonia B, and have oval-shaped nuclei with more and larger round-shaped heterochromatic regions than those in intermediate spermatogonia.

The second layer consisting of mid-pachytene cells, which exhibit intermediate levels of histone H1t. Stage VI sometimes has mitotic spermatogonia B. **Stages VII–VIII:** the basal cell layer primarily consists of preleptotene spermatocytes, which exhibit smaller and more round nuclei compared to spermatogonia B. The chromatin in preleptotene cells appears brighter, with larger and rounder heterochromatin aggregates compared to spermatogonia B. The number of preleptotene cells is approximately double that of spermatogonia B, resulting in an almost continuous basal layer. In the second layer, the late pachytene cells exhibit high levels of histone H1t. Stage VIII still has round spermatids but elongated spermatids are often absent from the luminal layer due to spermiation. **Stage IX:** the basal cell layer contains leptotene cells, which have even brighter chromatin than pre-leptotene cells and their heterochromatin aggregates are bigger and fewer. The second cell layer consists of late pachytene/diplotene cells which are strongly H1t-positive. Elongating spermatids are found in the lumen. **Stage X-XII:** the basal cell layer contains zygotene cells that display bright DAPI staining throughout the nucleus, with 2-3 bright heterochromatin aggregates in their periphery. The second layer consists of diplotene cells followed by third layer of elongated spermatids. In stage XII tubules, the diplotene cells progress into the first and second meiotic divisions, resulting in the presence of metaphase cells and secondary spermatocytes. Chromatin appears similar in late zygotene and early pachytene cells.

### Image analysis and quantification

Immunofluorescence images were captured by a microscope Axio Imager.Z2 (objectives EC Plan-Neofluar 20x/0.50 M27, EC Plan-Neofluar 40x/1.30 Oil DIC M27, and Plan-Apochromat 63x/1.40 Oil DIC M27) and ZEN imaging software (Carl Zeiss Microscopy). Images for figure panels were prepared with Adobe Photoshop CC19.

#### Quantification of axis morphology

Chromosome axis development was assessed by categorizing spermatocytes into three groups based on the length of axis stretches, identified by SYCP3 and/or SYCP2. The categories were defined as: short (punctate-looking to stretches shorter than 15 times their widths), long (at least one axis longer than 15 times its width, most axes fragmented), and full (all chromosome axes continuous and identifiable).

#### Quantification of SC morphology

SC initiation was assessed by categories of stretches detected through SYCP1. The categories were defined as: no/dotty (undetectable to strong dots of SYCP1), short (stretches longer than 2 but shorter than 15-20 times their widths), and long+full (at least one axis stretch per cell longer than 15-20 times its width, up to fully formed synapsis). In case of ambiguity regarding the length of the longest SC stretch, SC was classified as long if the remaining stretches were estimated to cover more than 40% of the total axis lengths. Importantly, SC was quantified independently from the axis, although they were co-stained. SC completion was evaluated in spermatocytes exhibiting fully formed axes that lacked IHO1 and were histone H1t-negative, corresponding to the early pachytene stage. The categories quantified were: fully synapsed, XY unsynapsed (but autosomes fully synapsed), and autosomes unsynapsed (at least one autosome showing a fork). Cells with asynapsis of both XY and autosomes were counted twice, once in each category. Illegitimate interactions between X/Y chromosomes and fully synapsed autosomes were quantified in H1t-negative, fully synapsed cells.

#### Quantification of meiosis markers and focus counts

Similar to a previously described strategy ^81,82^, DSB repair foci (RAD51, DMC1, RPA) were counted manually using the Count Tool in Adobe Photoshop and only foci that overlapped or closely associated with chromosome axes (identified by SYCP3) were considered in the counts. All DSB markers were co-stained with IHO1 and histone H1t for staging. The co-localization of RPA foci and ATR/BRCA1 foci/blobs was quantified only in S*ycp2^e16/e16^*, as ATR/BRCA1 has continuous distribution on the sex chromosomes in wild-type. To analyze the proportion of spermatocytes with a disrupted sex body, only fully synapsed spermatocytes were included. This approach aimed to minimize the impact of synapsis defects on the results.

### RNA isolation and Quantitative Reverse Transcription PCR (RT-qPCR)

RNA was extracted from whole testis of 13dpp old *Sycp2* mutant and wild-type mice using the NucleoSpin RNA kit (Macherey-Nagel, Germany) and reverse-transcribed with First Strand cDNA Synthesis Kit (Thermo Fisher Scientific, USA). For qPCR, the samples were prepared with iTaq Universal SYBR Green Supermix (Bio-Rad, USA) and analyzed on a CFX Opus 384 Real-Time PCR Detection System. Relative mRNA expression of *Zfy1* and *Zfy2* was calculated by comparative CT method using *S9*, *S12* and *Rsp16* for normalization and significance levels were calculated by one-way ANOVA followed by pairwise comparison of the mean expression between biological groups via Tukey’s HSD test using CFX Maestro software (Bio-Rad, USA). Primers are listed in Table.

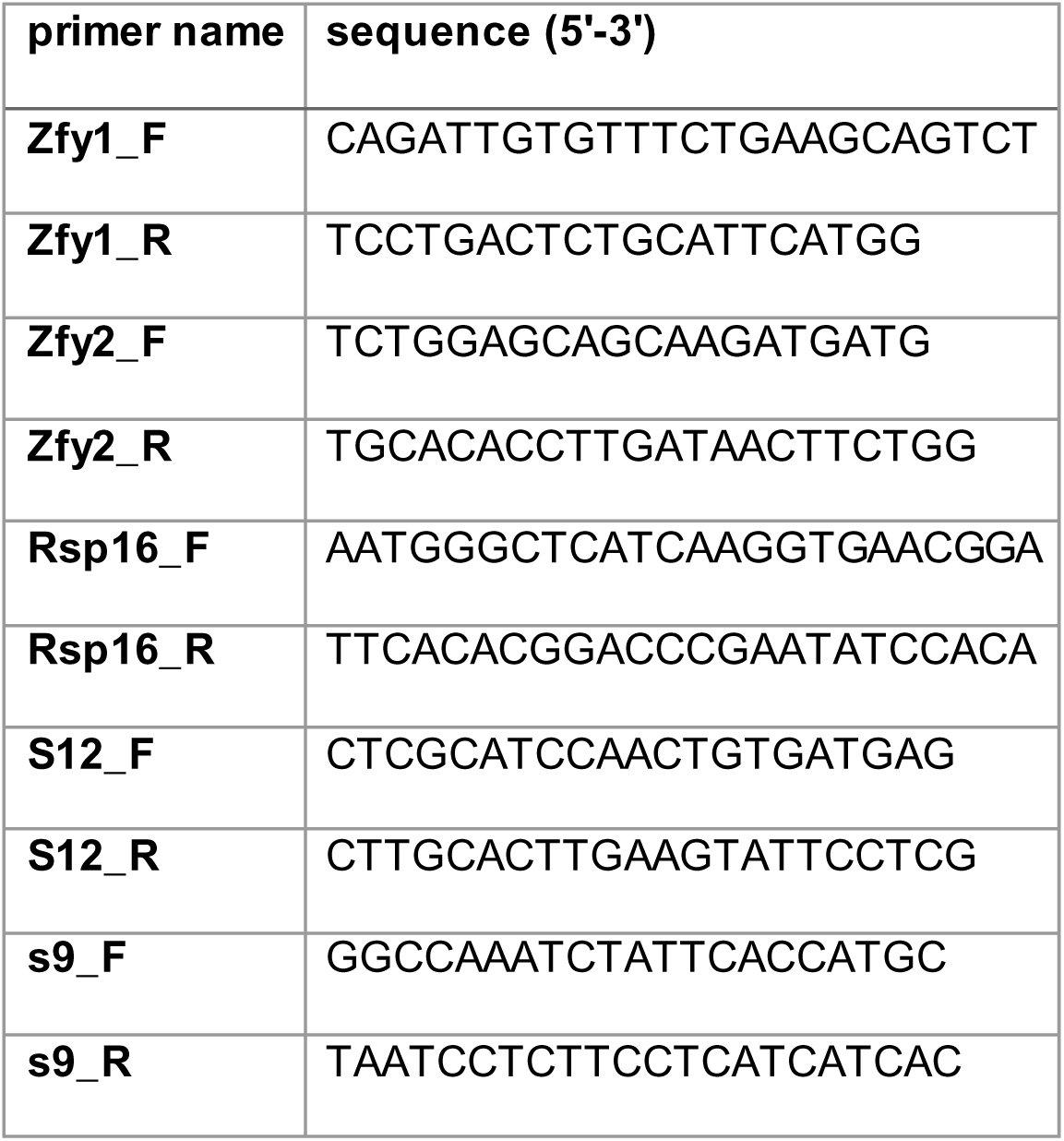

### Preparation of single-cell RNA sequencing samples

Testes of adult mice were detunicated, and seminiferous tubules were enzymatically dissociated in 1 mL HBSS containing Collagenase A (25 mg/mL), Dispase II (25 mg/mL), and DNase I (2.5 mg/mL) for 30 min at 37 °C with intermittent mixing. The digestion was quenched with 4 mL DMEM supplemented with 10% FCS, followed by centrifugation (400 × g, 5 min, RT). Cell pellets were washed in DPBS-S (PBS containing 10% knockout serum) and passed through a 40 µm mesh filter.

Testes were collected from two biological replicates per genotype. Testes from each WT mouse were split to generate samples processed with or without dead cell removal. Dead cell removal partially depleted post-meiotic sperm populations, thereby increasing the representation of prophase-stage spermatocytes, which are otherwise underrepresented in scRNA-seq libraries from adult WT testes and complicate direct comparison with mutants lacking post-meiotic cells. Mutant samples were processed without dead cell removal to retain spermatocytes prior to mid-pachytene apoptosis.

Dead-cell removal was performed using the Miltenyi Dead Cell Removal Kit (Miltenyi Biotec, Cat. #130-090-101) according to the manufacturer’s instructions. Viable cells were subsequently enriched by differential-speed centrifugation (400/200/100 × g).

In all samples, single cells were resuspended in PBS containing 0.04% BSA and labeled with Cell Multiplexing Oligos (CMOs) from 10x Genomics 3’CellPlex Kit Set A (Part number 1000261) according to the manufacturer’s protocol. WT samples were pooled separately from a pool comprising *Sycp2^e16/e16^* and *Hormad2^-/-^* samples. Sequencing libraries were prepared following 10x Genomics Single Cell 3’ v3.1 (Dual Index) protocols with Feature Barcode technology (CG000388). WT and mutant pools were sequenced separately on Illumina NovaSeq 6000/NovaSeq S4 v1.5 (200cyc) flow cells. Separate sequencing of WT and mutant pools was implemented to avoid interference from abundant post-meiotic cells in WT samples and to ensure adequate representation of early meiotic prophase cells in mutant datasets.

### Processing and clustering of single-cell RNA sequencing data

Single-cell RNA sequencing (scRNA-seq) data was generated using the 10x Genomics Chromium platform. The raw sequencing data were processed through the Cell Ranger pipeline (v7.1.0) ^83^ using default parameters to generate gene-expression matrices, using the *GRCm38_mm10* genome build from Ensembl 98. Further analysis were carried out in R4.4.1 (https://www.r-project.org/) using the Seurat package (v5.0) ^84^. Quality control measures included the exclusion of cells with high mitochondrial RNA content (>10%) and low feature counts (<200 genes). Genes expressed in fewer than three cells across the dataset were also filtered out to ensure robust downstream analyses. Normalization was performed using the LogNormalize method with a scaling factor of 10,000, correcting for sequencing depth and technical variation. Principal Component Analysis (PCA) was applied to the top 2000 variable features and eight principal components were used to reduce dimensionality, followed by Uniform Manifold Approximation and Projection (UMAP) for two-dimensional visualization, using a minimum distance of 0.3 and a spread of 1.0. Cell clustering was conducted using the Louvain algorithm with resolution parameters optimized between 0.4 and 1.2, and a neighborhood size (k) of 20 to identify distinct cell populations. Unsupervised clustering was performed, using clustering resolution of 0.5, twenty-one clusters were identified, which contained meiotic and somatic cells. Notably, prophase I cells at the zygotene and early pachytene stages co-localized within a single cluster. To achieve a better separation, supervised clustering was performed by subsetting specific clusters of interest (Supplementary Table 4) from the Seurat object generated from unsupervised clustering. Clusters were selected based on predefined identifiers (Supplementary Table 3) using the “subset” function in Seurat.

After identifying germline populations, we performed supervised re-clustering of these cells using stage-specific germline marker genes (Supplementary Tables 3 and 5) and a clustering resolution of 1 to increase the resolution of meiotic prophase I subpopulations. This procedure initially yielded 19 clusters. Examination of marker gene expression profiles revealed that some clusters represented closely related cell populations with very similar transcriptional identities but differing representation across genotypes. Specifically, two pairs of clusters displayed highly similar expression signatures characteristic of zygotene-stage spermatocytes or undifferentiated spermatogonia, respectively. Because these paired clusters were not distinguished based on dominantly expressed stage-defining marker genes, each pair was consolidated into a single cluster: clusters “6a” (WT enriched) and “6b” (mutant enriched) were merged and retained as cluster 6 (zygotene), and clusters “7a” (WT enriched) and “7b” (mutant enriched) were merged and retained as cluster 7 (undifferentiated spermatogonia). All remaining clusters displayed distinct and robust marker gene expression patterns and were retained as separate populations. Cell-type annotations were assigned based on expression of curated stage-specific marker genes (Fig. 6b; Supplementary Fig. 11; Supplementary Tables 3–5).

### All-autosome-normalized chromosome-wide expression calculations

To quantify meiotic sex chromosome inactivation (MSCI) at single-cell resolution, we adapted the X:A framework of ^85^ to our Seurat-based scRNA-seq dataset. Ensembl gene identifiers were mapped to Seurat feature names (rowname) using a lookup table. Genes were assigned to individual chromosomes and autosomal genes were pooled for all-autosome expression calculation (chromosomes 1–19, hereafter A). For each genotype (WT, *Hormad2⁻^/⁻^*, *Sycp2^e16/e16^*), cells were grouped by cluster identity and genotype. Log-normalized expression values were retrieved from the RNA assay (layer). To reduce noise from sparsely detected genes, we required each gene to be expressed (expression > 0) in at least 30% of cells within WT reference cell populations (min_expressed = 0.3). This filtering was applied to each genotype to define gene sets for each chromosome and the pooled autosomes that were shared by all three genotypes (WT, *Hormad2^⁻/⁻^*, and *Sycp2^e16/e16^*). This approach was used to maintain identical gene composition across genotypes and avoid biases from genotype-specific expression dropout. For each cell, we calculated chromosome-wide expression values by computing the mean expression of all genes on each chromosome. In case of zero expression value for a chromosome, the value was replaced with NA and the corresponding cell was excluded from downstream ratio-based analyses. To obtain all-autosome-normalized ratios, the mean expression of genes on individual chromosomes was divided by the mean expression of genes on all autosomes (chromosomes 1–19).

These all-autosome-normalized expression ratios for individual early-to-mid pachytene cells were divided by the median all-autosome-normalized expression of the same chromosome in preleptotene–leptotene or zygotene cells of the same genotype. The results from these ratio calculations were compared between genotypes using the Wilcoxon rank-sum test, followed by Benjamini–Hochberg correction for multiple testing. Data visualization was performed using GraphPad Prism. This analysis followed the computational framework described by previously ^59^, with minor modifications tailored to the present dataset.

### *Zfy1* and *Zfy2* expression quantification in scRNA-seq data

*Zfy1* and *Zfy2* expression was quantified using log-normalized expression values from the Seurat RNA assay (data layer; log1p-normalized). Distinct stages were statistically compared using two-tailed unpaired t-test with Welch’s correction. Data visualization was performed using GraphPad Prism.

### Reagents and resources

Mice, antibodies, software and datasets used in this study are listed with references in the “Reagents and resources” section of the supplementary information.

### Biological materials availability

Transgenic mouse strains and plasmids produced in this study are available from the authors upon request.

### Statistics and Reproducibility

Graphs were plotted by Graphpad Prism 10. Statistical test were calculated by Graphpad Prism 10 or R version 4.1.3 ^86^ for likelihood ratio tests and R version 4.4.1 ^86^ for scRNAseq data. The likelihood ratio tests were implemented in the R packages lmerTest and lme4 ^87,88^. The R code was published previously ^82^. Details of statistical tests for quantitative reverse transcription PCR and scRNA-seq analysis are described in the relevant sections of the methods. We used two-tailed statistics if this option was available in the statistical tests (Wilcoxon rank-sum test and t tests but not the Likelihood ratio test ^82^). There were no adjustments for multiple comparisons with the exception of the scRNA-seq and RT-qPCR analysis.

Specific randomization methods were not used. The study relied on comparison of wild type and various mutant mice that were generated by random segregation of alleles during sexual reproduction. Where control versus mutant mice were compared, samples were processed in parallel to eliminate batch effects. No formal sample-size calculations were performed and investigators were not blinded to the origin of samples during the assessment of results. All phenotypes were observed in at least two animals of each genotype and quantifications represent analysis of at least two independent animals. Sample sizes, statistical tests, and p-values are indicated in the figure legends and the source data file.

Reproducibility of representative experiments: images of immunoprecipitations, immunoblots, immunofluorescence in histological sections, nuclear surface-spread meiocytes or oocytes and Y2H assays represent at least two independent (biological) repetitions of experiments.

### Data and Code availability

The authors declare that all data supporting the findings of this study are available within the paper, its Supplementary Information, and provided source data files. Raw data underlying figures and large-scale datasets have been deposited in public repositories, including BioStudies ^89,90^ and ArrayExpress. Custom code used for analysis is available via Zenodo. All resources will be made publicly available upon peer-reviewed publication; however, they are available from the corresponding author upon reasonable request prior to that time.

## ACKNOWLEDGMENTS

We thank D. Schade from the Facility for Reproductive and Transgenic Technologies in Dresden for the generation of *Sycp2^e16^* mice, M. Munzig and V. Hantzsch for lab support, R. Jessberger for departmental support and sharing ideas and reagents (anti-SYCP3 antibody), J. Weir and E. Selezneva for sharing ideas about the structural basis of HORMAD-SYCP2 interactions, J.M.A. Turner for insightful discussions about the theoretical framework on meiotic checkpoint control.

K. R., S.V.C., A.G., V.T., G.G., C.R., A.B., T.S., M.W. and A. T. were supported by the Deutsche Forschungsgemeinschaft (DFG; grants: TO421/6-2/3, TO421/10-1, TO421/11-1, TO421/12-1 and TO421/14-1), and core funding from the Faculty of Medicine at the TU Dresden. A.P. and A.D were supported by the DFG Research Infrastructure NGS_CC (INST 269/768, project 407482635, DRESDEN-concept Genome Center). NGS library preparation, data production and analyses were carried out at DcGC Dresden-concept Genome Center - a core facility of the CMCB and Technology Platform of the TUD Dresden University of Technology.

## AUTHOR CONTRIBUTIONS

K. R., S.V.C.: conceptualization, methodology, investigation, formal analysis, visualization, writing–original draft preparation, data curation and project administration.

A.G.: methodology, investigation, formal analysis, visualization, data curation V.T., G.G., C.R.: methodology, investigation, formal analysis, visualization A.B., T.S., M.W.: investigation

A.P.: methodology, formal analysis, data curation

A.T.: conceptualization, writing–original draft preparation, review and editing, supervision, project administration, funding acquisition.

