## Supplementary Information for "Chromosome axis protein SYCP2 recruits HORMAD2 to enable meiotic synapsis quality control in mice"

### Chromosomal synapsis surveillance requires HORMAD2 recruitment by the chromosome axis protein SYCP2 in mouse meiosis

#### SUPPLEMENTARY FIGURE AND TABLE LEGENDS

**Supplementary Figure 1. Current model and outstanding questions regarding the molecular network underlying asynaptic axis–triggered ATR activation and the resultant prophase checkpoint in mammalian meiosis.**

Green arrows indicate promoting effects; the blue double-headed arrow denotes inferred but undefined physical interactions that enable recruitment of HORMAD1 and HORMAD2 to unsynapsed axes. The prevailing model of the synapsis checkpoint proposes that preferential enrichment of HORMAD1 and HORMAD2 on unsynapsed chromosome axes forms the basis of asynapsis sensing [1–7]. Axis-bound HORMADs are thought to establish an ATR-mediated signalling cascade that is reinforced by positive feedback and feedforward mechanisms [8–18], resulting in robust ATR activity on both axes and associated chromatin loops within unsynapsed regions. This signalling drives meiotic silencing of unsynapsed chromatin (MSUC) and DNA damage response (DDR)–like signalling, both of which may contribute to the checkpoint mechanisms that eliminate persistently asynaptic meiocytes [8–10,13,15,17–23]. Key outstanding questions include: How are HORMAD1 and HORMAD2 recruited to chromosome axes? What is the functional significance of their axial localization? And do soluble, off-axis pools of HORMADs contribute to ATR activation and DDR-like responses during asynaptic meiosis?

**Supplementary Figure 2. AlphaFold 3 models for HORMAD1 and HORMAD2 interactions with a predicted closure motif in SYCP2.**

**a-b** AlphaFold 3 model predictions of complexes between a fragment of mouse SYCP2 (amino acids (aa) 1-500) and full-length HORMAD1 (aa 1-392) **(a)** or HORMAD2 (aa 1-306) **(b)**. Colors: SYCP2 is yellow, HORMA domain safety belts (HORMAD1, aa 180-230; HORMAD2, aa 184-235) are pink, and the remaining HORMAD regions are blue. Insets of the full models (far left) are enlarged (middle), showing the predicted closure motif of SYCP2 (aa 400-406) wrapped by the safety belts on the surface of the HORMA domains. Right panels show predicted aligned error (PAE) plots with ipTM and pTM scores.

**Supplementary Figure 3. Spermatogenic failure despite efficient axis formation in *Sycp2*$^{e16/e16}$.**

**a** Agarose gel electrophoresis of PCRs using genomic DNA as a template and primers annealing to genomic loci flanking the 16th exon of *Sycp2*. Predicted PCR product sizes: 1057 bp in WT, and 643 bp in *Sycp2*$^{e16/e16}$. **b** Agarose gel electrophoresis of reverse transcription PCRs using total testis RNA as a template from 13 days post-partum (dpp) mice, amplifying sequences between the 11th and 19th exons of *Sycp2*. Predicted PCR product sizes: 713 bp in WT, and 638 bp in *Sycp2*$^{e16/e16}$. **a-b** No-template PCRs (NT) are shown. **c** SDS-PAGE immunoblots of total protein extracts from testes of WT and *Sycp2*$^{e16/e16}$ mice at 13 dpp. GAPDH (cytoplasmic marker) and histone H3 (chromatin marker) serve as loading controls. **d** Quantification of total SYCP2 protein abundance in testes of 13 dpp mice. SYCP2 immunoblot signals from testis extracts of *Sycp2*$^{e16/e16}$ mice were normalized to corresponding signals from wild-type controls. Bar indicates mean = 1.29 from three experiments; two-tailed one-sample *t*-test, $P = 0.0111$. **e** DNA staining (DAPI) and immunostaining of cleaved PARP and histone H1t in testis cryosections from adult mice, showing seminiferous tubules in epithelial cycle stages V-VI (see Methods for staging). The images show the presence of pachytene spermatocytes and post-meiotic cells in the WT testis section and their absence in the *Sycp2*$^{e16/e16}$ testis section, consistent with spermatocyte elimination at epithelial cycle stage IV in the mutant, as shown in Fig. 1d. Sertoli cells (Se), type B spermatogonia (Bsg), late pachytene spermatocytes (Lpa), round spermatids (rs), and sperm (sp) are marked. Scale bars, 50 µm. **f** Immunostained nuclear spreads of spermatocytes from adult mice. Morphology categories for axes (relevant to Fig. 1e) indicated at the bottom of the panel characterize the prophase stages indicated above each image. Unsynapsed (unsy) and synapsed (sy) regions of autosomes are marked in late zygotene. X and Y chromosomes and the pseudoautosomal region (PAR; white arrowheads) are marked in enlarged insets of early pachytene images. Scale bars, 10 µm (cell) and 5 µm (insets). Source data are provided as a Source Data file.

**Supplementary Figure 4. HORMAD2 is present in *Sycp2*$^{e16/e16}$ spermatocytes but depleted from the chromosome axes.**

**a-b** Immunostaining in nuclear spreads of spermatocytes **(a)** from adult mice or oocytes **(b)** from 17 dpc foetuses. Scale bars, 10 µm (cell) and 5 µm (inset). Images with matched exposure and levelling are shown within each stage. The channels were differentially levelled between different stages to optimize viewing. **a** X and Y chromosomes, the pseudoautosomal region (PAR; white arrowheads), and examples of synapsed (sy) and unsynapsed (unsy) regions of autosomes are marked in enlarged insets. **b** Unsynapsed axes are marked by yellow arrows**. c** DNA staining (DAPI) and immunostaining of HORMAD2 in cryosections of testes from adult mice. Stages of the seminiferous epithelial cycle are indicated (see 'Methods' for staging). Sertoli cells (se), intermediate spermatogonia (isg), pachytene (pa) spermatocytes, and round spermatids (rs), are marked in the insets. White arrowheads mark the sex body, identified by enriched HORMAD2 signal in WT spermatocytes**.** Note the absence of this enrichment despite presence of nuclear HORMAD2 in pachytene spermatocytes in the *Sycp2*$^{e16/e16}$ testis. Scale bars, 25 µm (section) and 10 µm (inset).

**Supplementary Figure 5. Y2H interactions between HORMADs and complex formation between SYCP2 and HORMAD2.**

**a** SDS-PAGE immunoblot analysis of protein extracts from testes of WT juvenile mice (11 dpp). Total lysate (Input), and immunoprecipitates with anti-SYCP2 (SYCP2 IP) or non-specific rabbit IgG (Rb-IgG IP) antibodies are shown. Asterisks (**) mark an aspecific band that is observed in total protein extracts when analysed in 10% PAGE (see also Fig. 2d-e) but undetectable in the immunoprecipitated product. **b** Yeast two-hybrid interaction assays testing interactions between fragments corresponding to distinct domains of HORMAD1 and HORMAD2. Amino acid (aa) positions of fragment ends are indicated. Yeast cultures are shown after 3 and 5 days of growth on dropout plates. For negative control, proteins of interest were tested in transformations where either the Gal4-binding domain (Gal4-BD) or the Gal4-activation domain (Gal4-AD) vectors were empty. **c** Schematics of HORMAD1 and HORMAD2 domain structures and summary of Y2H interactions between their HORMA domains and depicted protein fragments. Boxes mark the positions of HORMA domains and predicted closure motifs (CMs) as previously described [24]. Numbers represent amino acid positions. Strength of Y2H interactions is graded from no interaction (−) to very strong interaction (++++).

**Supplementary Figure 6. Recombination foci in WT**, ***Sycp2***$^{e16/e16}$ **and** ***Hormad2***$^{-/-}$ **spermatocytes.**

Immunostaining of nuclear surface spreads of spermatocytes from adult mice. For recombination markers (RAD51, DMC1, and RPA), images are shown with matched exposure and levelling across all genotypes and prophase stages (**d-f**). For stage markers (IHO1 and histone H1t), exposure and levelling are matched across genotypes for late zygotene and early pachytene; at other stages, exposure and levelling are matched across genotypes but differentially adjusted between stages for optimal viewing. SYCP3 was differentially levelled to optimize visualization. Histone H1t is shown in miniaturized images in the bottom right corners of full-size cell images. Panels **a-c** show IHO1 and histone H1t images in cells corresponding to Fig. 3d-f. Scale bars, 10 µm.

**Supplementary Figure 7. IHO1 localization in WT, *Sycp2*$^{e16/e16}$ and *Hormad2*$^{-/-}$ spermatocytes. a-c** Immunostaining of nuclear surface spreads of spermatocytes from adult mice. Images with matched exposure and levelling are shown for stage markers IHO1 and histone H1t (miniaturized images in bottom right corner of full-size cell images) across genotypes and prophase stages. To illustrate IHO1 dynamics, early zygotene, late zygotene, and early pachytene are compared between WT **(a)**, *Sycp2*$^{e16/e16}$ **(b)**, and *Hormad2*$^{-/-}$ **(c)** genotypes; late pachytene is only shown in WT **(c)** because the mutants lack spermatocytes beyond mid-pachytene. SYCP3 was differentially levelled to optimize viewing. Scale bars, 10 µm. **d** Quantification of IHO1 localization on axes of spermatocytes that have fully developed axes and are negative for histone H1t (corresponding to late zygotene and early pachytene in wild-type spermatocytes). Graph shows data points and weighted averages (bars) of percentages of IHO1-positive spermatocytes (WT = 48.60%, *Sycp2*$^{e16/e16}$ = 47.90%, and *Hormad2*$^{-/-}$ = 50.33%) from three biological replicates. Number of analyzed cells (*n*) per genotype is indicated. Likelihood-ratio test (chi-squared distribution) was used to determine whether the proportion of IHO1-positive cells among histone H1t-negative, fully-axis-displaying spermatocytes differed significantly between mutants and wild type. Source data are provided as a Source Data file.

**Supplementary Figure 8. *Hormad2*$^{-/-}$ spermatocytes display mild synapsis defects.**

**a-b** Immunostained nuclear spreads of early pachytene *Hormad2*$^{-/-}$ spermatocytes from adult mice illustrating synapsis categories quantified in Fig. 4c-d. SYCP1 marks synapsed regions; fully formed axes (SYCP3) and the absence of IHO1 identify the cells as early pachytene. Pseudoautosomal regions (PARs; white arrowheads in **a-b**), synapsed (sy) and unsynapsed (unsy) autosomal regions (**a**), and illegitimate interactions between the ends of the X or Y chromosomes and fully synapsed autosomes (white arrows in **a-b**) are indicated. Scale bars, 10 µm (whole cell) and 5 µm (insets).

**Supplementary Figure 9. The 16th exon of *Sycp2* is required for efficient BRCA1 accumulation on sex chromosomes.**

**a-b** Immunostained nuclear surface spreads of early pachytene spermatocytes from adult mice. Images with matched exposure and levelling are shown for γH2AX, RPA and BRCA1 signals. X and Y chromosomes, PARs (white arrowheads in **a-b**), an illegitimate end-to-end association between the X chromosome and an autosome (yellow arrow in **a**), and sites of sex-chromosome-axis-associated RPA foci (arrows in **b**) are marked in enlarged insets. Scale bars, 10 µm (cells), 5 µm (insets). **c** Quantification of overlap between axis-associated RPA and BRCA1 signals on sex chromosomes in *Sycp2*$^{e16/e16}$ spermatocytes. The numbers of analysed cells (n) correspond to two experiments (differentiated by blue and red colors); medians (bars) are 80% (BRCA1 colocalizing with RPA), and 90% (RPA colocalizing with BRCA1).

**Supplementary Figure 10. Unsupervised clustering of pooled scRNAseq data from testes of WT, *Sycp2*$^{e16/e16}$, and *Hormad2*$^{-/-}$ mice.**

**a** UMAP visualization of whole testis scRNA-seq data, showing cell populations identified by unsupervised Seurat-based clustering (cluster resolution parameter = 0.5). Pooled data are shown from testes of WT (four samples from two biological replicates), *Sycp2*$^{e16/e16}$ (two biological replicates), *Hormad2*$^{-/-}$ (two biological replicates) mice. The numbers represent clusters of distinct testicular cell populations as listed in **b**. **b** Dot plot heatmaps showing expression of selected marker genes. The x-axis lists representative marker genes and the cell types that primarily express them; the y-axis lists cluster numbers corresponding to **a**. Expression levels are represented as z-scores ranging from –1 to 2. The size and color intensity of each dot indicate the proportion of cells expressing the gene and the relative expression level, respectively. The full list of marker genes used for identifying spermatogenic and somatic clusters, and the distinguishing characteristics of the cell clusters are listed in Supplementary Table 3 and 4.

**Supplementary Figure 11. Expression of marker genes in spermatogenic populations identified by supervised clustering.**

Dot plot showing scRNA-seq–derived expression levels of all the marker genes used for supervised clustering and subsequent annotation of spermatogenic cell populations. Genes used exclusively for cell-type annotation are boxed. The x-axis lists marker genes; the y-axis lists cluster numbers corresponding to those in Fig. 6a–b. Clusters 6 and 7 are shown both as their constituent subclusters (6a/6b and 7a/7b) and as pooled clusters. In both cases, WT cells are preferentially enriched in subcluster “a” and mutant cells in subcluster “b,” despite highly similar expression profiles of stage-defining markers. Expression values are represented as z-scores ranging from –1 to 2. Dot size and color intensity indicate the proportion of cells expressing each gene and the relative expression level, respectively. Colored bars below the x-axis denote the cell populations predominantly expressing the corresponding markers. A complete list of marker genes used for clustering and their defining characteristics is provided in Supplementary Tables 3 and 5.

**Supplementary Figure 12. scRNA-seq UMAPs of spermatogenic populations from individual WT, *Sycp2$^{e16/e16}$*, and *Hormad2$^{-/-}$* samples.**

UMAP visualization of single-cell RNA sequencing data showing spermatogenic cell populations identified by supervised clustering using germline marker genes listed in Supplementary Fig. 10 (see also methods). Cluster numbers correspond to spermatogenic stages listed in Fig. 6b. **a** UMAPs for two biological replicates (left and right columns) of WT samples processed either without (top row) or with (bottom row) sperm and dead cell depletion. UMAPs for two biological replicates (left and right columns) of *Sycp2$^{e16/e16}$* (top row), and *Hormad2 $^{-/-}$* (bottom row) samples.

**Supplementary Figure 13. scRNA-seq reveals impaired sex chromosome silencing in *Hormad2$^{-/-}$* and *Sycp2$^{e16/e16}$* spermatocytes.**

**a** All-autosome-normalized chromosome-wide expression in early-to-mid pachytene cells relative to preleptotene-leptotene stages. Each data point represents normalized expression for chromosome 9 (Chr9/A), the X chromosome (ChrX/A), or the Y chromosome (ChrY/A) in early-mid pachytene cells divided by the median normalized expression of the same chromosome in preleptotene-leptotene cells (see Methods for details). *P* values were determined using the Wilcoxon rank-sum test. **b-c** All-autosome-normalized chromosome-wide expression in early-to-mid pachytene cells relative to zygotene (**b**) and preleptotene-leptotene (**c**) stages. Each data point represents all-autosome-normalized expression for chromosome 1 (Chr1/A), chromosome 10 (Chr10/A), or chromosome 19 (Chr19/A) in individual early-mid pachytene cells divided by the median normalized expression of the same chromosome in zygotene (**b**) or preleptotene-leptotene (**c**) cells. Data points represent individual cells; medians (black bars), and *P* values from the Wilcoxon rank-sum test are indicated. Source data are provided as a Source Data file.

**Supplementary Table 1. Fertility in female *Sycp2$^{e16/e16}$* and *Hormad2$^{-/-}$* mice**

Quantification of pup numbers from crosses of wild-type (WT) male mice with wild-type, *Sycp2*$^{e16/e16}$ or *Hormad2*$^{-/-}$ female mice. Statistical significances were calculated by two-tailed unpaired t-test with Welch correction.

**Supplementary Table 2. Summary of single-cell RNA-seq quality metrics in WT and mutant testes samples.**

Key sequencing and mapping statistics for four WT, two *Hormad2*$^{-/-}$, and two *Sycp2*$^{e16/e16}$ samples. Two biological replicates of WT samples were processed either without (1 and 2) or with (3 and 4) sperm and dead cell depletion.

**Supplementary Table 3. Germline and somatic marker genes used for cluster identification.**

Two sets of marker genes used to identify and separate germline and somatic cell populations in the single-cell RNA-seq data. Somatic markers (top section) were used to identify somatic cell-enriched clusters, which were excluded from further analysis. Germline markers (bottom section) were used to identify spermatogenic clusters and to perform supervised re-clustering of spermatogenic subpopulations.

**Supplementary Table 4. Summary of unsupervised clusters and marker-based cell-type annotations.**

Clusters identified by unsupervised analysis of single-cell RNA-seq data and annotated based on expression of somatic and germline marker genes. Identified cell types include spermatogonia, spermatocytes, spermatids, Leydig cells, Sertoli cells, macrophages, and endothelial cells. Clusters with germline marker enrichment were subsequently selected for supervised re-clustering (see Supplementary Table 5). Key marker genes identifying dominant cell types in each cluster are listed.

**Supplementary table 5. Summary of supervised clusters and marker-based cell-type annotations.**

Clusters identified by supervised analysis of spermatogenic cells of single-cell RNA-seq data and annotated based on expression of germline marker genes. Key marker genes identifying dominant cell types in each cluster are listed.

**Table of reagents and resources**

| REAGENT or RESOURCE | SOURCE | IDENTIFIER |
|---|---|---|
| Antibodies | | |
| Chicken polyclonal anti-SYCP3 | [25] | N/A |
| Mouse monoclonal anti-SYCP3 | [26] | N/A |
| Chicken polyclonal anti-IHO1 | [27] | N/A |
| Rabbit polyclonal anti-IHO1 | [27] | N/A |
| Guinea polyclonal pig anti-HORMAD1 | [3] | N/A |
| Rabbit polyclonal anti-HORMAD1 | [3] | N/A |
| Guinea pig polyclonal anti-HORMAD2 | [1] | N/A |
| Guinea pig polyclonal anti-Histone H1t | [28] | N/A |
| Guinea pig polyclonal anti-Histone H1t | This study | N/A |
| Chicken polyclonal anti-SYCP1 | [28] | N/A |
| Rabbit polyconal anti-BRCA1 | This study | N/A |
| Rabbit polyclonal anti-NOBOX | This study | N/A |
| Rabbit polyclonal anti-SYCP1 | Abcam | Cat# ab15090 RRID: AB_301636 |
| Rabbit polyclonal anti-SYCP2 | This study | N/A |
| Guinea pig polyclonal anti-SYCP2 | This study | N/A |
| Rabbit polyclonal anti-cleaved PARP (Asp214) | Cell signaling | Cat# 9544; RRID: AB_2160724 |
| Mouse monoclonal anti-GAPDH | Santa Cruz | Cat# sc-32233 RRID: AB_627679 |
| Rabbit polyclonal anti-Histone H3 | Abcam | Cat# ab1791 RRID: AB_302613 |
| Mouse monoclonal anti-α-TUBULIN | Sigma-Aldrich | Cat# T6199 RRID: AB_477583 |

| | | |
|---|---|---|
| Mouse monoclonal anti-DMC1 (2H12/4) | Abcam | Cat# ab11054; RRID: AB_297706 |
| Guinea pig polyclonal anti-DMC1 | This study | N/A |
| Rabbit polyclonal anti-Rad51 | Abcam | Cat# ab176458; RRID: AB_266540 5 |
| Rabbit monoclonal anti-RPA32/RPA2 (EPR2877Y) | Abcam | Cat# ab76420; RRID: AB_15243 36 |
| Mouse monoclonal anti-phospho-Histone H2A.X (Ser139) | Millipore | Cat# 05-636; RRID: AB_309864 |
| Goat polyclonal anti-ATR (N-19) | Santa Cruz | Cat# sc-1887; RRID: AB_630893 |
| Mouse monoclonal anti-Pol II | Santa Cruz | Cat# sc-56767 RRID:AB_785522 |
| Mouse monoclonal anti-SYCP3 | Abcam | Cat# ab 97672 RRID:AB_1067884 1 |
| Goat anti-rabbit IgG-HRP | Jackson ImmunoResearch | Cat# 111-035-003; RRID: AB_2313567 |
| Goat anti-guinea pig IgG-HRP | Jackson ImmunoResearch | Cat# 706-035-148; RRID: AB_2340447 |
| Goat anti-mouse IgG-HRP | Jackson ImmunoResearch | Cat# 115-035-003; RRID: AB_10015289 |
| Goat anti-Rabbit IgG-AF405 | Thermo Fisher Scientific | Cat# A-31556; RRID: AB_221605 |
| Goat anti-rabbit IgG-AF488 | Thermo Fisher Scientific | Cat# A-11034; RRID: AB_2576217 |

| | | |
|---|---|---|
| Goat anti-rabbit IgG- AF568 | Thermo Fisher Scientific | Cat# A-11036; RRID: AB_10563566 |
| Goat anti-Rabbit IgG- AF647 | Thermo Fisher Scientific | Cat# A-21244; RRID: AB_2535812 |
| Donkey anti-guinea pig IgG- DyLight™405 | Jackson ImmunoResearch | Cat# 706-475-148; RRID: AB_2340470 |
| Goat anti-guinea pig IgG-AF488 | Thermo Fisher Scientific | Cat# A-11073; RRID: AB_2534117 |
| Goat anti-guinea pig IgG-AF568 | Thermo Fisher Scientific | Cat# A-11075; RRID: AB_2534119 |
| Goat anti-guinea pig IgG-AF647 | Thermo Fisher Scientific | Cat# A-21450 RRID: AB_2735091 |
| Donkey anti-guinea pig IgG-AF647 | Jackson ImmunoResearch | Cat# 706-605-148 RRID: AB_2340476 |
| Goat anti-mouse IgG-AF405 | Thermo Fisher Scientific | Cat# A-31553; RRID: AB_221604 |
| Goat anti-mouse IgG-AF488 | Thermo Fisher Scientific | Cat# A-11029; RRID: AB_2534088 |
| Goat anti-mouse IgG-AF568 | Thermo Fisher Scientific | Cat# A-11031; RRID: AB_144696 |
| Goat anti-chicken IgY-AF405 | Abcam | Cat# ab175675 RRID: AB_2810980 |

| | | |
|---|---|---|
| Goat anti-chicken IgY-AF488 | Thermo Fisher Scientific | Cat# A-11039; RRID: AB_2534096 |
| Goat anti-chicken IgY-AF568 | Thermo Fisher Scientific | Cat# A-11041; RRID: AB_2534098 |
| Goat anti-rat IgG-AF488 | Thermo Fisher Scientific | Cat# A-11006; RRID: AB_2534074 |
| Goat anti-rabbit IgG-AF647 | Thermo Fisher Scientific | Cat# A-21245; RRID: AB_2535813 |
| Goat anti-chicken IgY-AF647 | Thermo Fisher Scientific | Cat# A-21449; RRID: AB_2535866 |
| Donkey anti-rabbit IgG-AF647 | Jackson ImmunoResearch | Cat# 711-495-152; RRID: AB_2315775 |
| Bovine anti-goat IgG-RRX | Jackson ImmunoResearch | Cat# 805-295-180; RRID: AB_2340881 |
| Donkey anti-mouse IgG-AF350 | Thermo Fisher Scientific | Cat# A-10035; RRID: AB_2534011 |
| Donkey anti-guinea pig IgG-AF488 | Jackson ImmunoResearch | Cat# 706-545-148; RRID: AB_2340472 |
| Donkey anti-rabbit IgG-DyLight™ 488 | Jackson ImmunoResearch | Cat# 711-485-152; RRID: AB_2492289 |
| Chemicals, peptides, and recombinant proteins | | |

| | | |
|---|---|---|
| | | Cat#<br>Cas# |
| 6xHis-tagged SYCP2 C-terminus | This study | N/A |
| 6xHis-tagged H1t C-terminus | This study | N/A |
| 6xHis-tagged BRCA1 C-terminus | This study | N/A |
| 6xHis-tagged NOBOX | This study | N/A |
| 6xHis-tagged DMC1 | This study | N/A |
| Critical commercial assays | | |
| NHS-activated Sepharose 4 Fast Flow beads | Cytiva/GE Healthcare | Cat# 17-0906-01 |
| Kits | | |
| NucleoSpin RNA kit | Macherey-Nagel | Cat# 740955.50 |
| First Strand cDNA Synthesis Kit | Thermo Fisher Scientific | Cat# K1612 |
| iTaq Universal SYBR Green Supermix | Bio-Rad | Cat# 1725120 |
| Experimental models: Organisms/strains | | |
| Y2HGold Yeast strain | Clontech | Cat# 630498 |
| *E. coli* strain BL21(DE3) pLysS | | |
| *E. coli* strain BL21 *tRNA* | | |
| *E. coli* strain Rosetta2(DE3)pLysS | | |
| Mouse/ *Sycp2*$^{+/+}$ and *Sycp2*$^{e16/e16}$ | This study | |
| Mouse/ *Hormad2*$^{-/-}$ | [2] | |
| Mouse/ *Iho1*$^{+/+}$ and *Iho1*$^{-/-}$ | [27] | |
| Mouse/ *Hormad1*$^{+/+}$ and *Hormad1*$^{-/-}$ | [3] | |
| Mouse/ *Spo11*$^{+/+}$ and *Spo11*$^{-/-}$ | [29] | |
| Mouse/ *Dmc1*$^{+/+}$ and *Dmc1*$^{-/-}$ | [30] | |
| Mouse/ *Sycp1*$^{+/+}$ and *Sycp1*$^{-/-}$ | [31] | |
| Mouse/ *Spo11*$^{-/-}$ *Sycp2*$^{+/+}$ and *Spo11*$^{-/-}$ *Sycp2*$^{e16/e16}$ | | |
| Mouse/ *Dmc1*$^{-/-}$ *Sycp2*$^{e16/e16}$ | This study | |
| Mouse/ *Iho1*$^{-/-}$ *Sycp2*$^{e16/e16}$ | This study | |

| | | |
|---|---|---|
| Mouse/ *Sycp1*$^{+/+}$ *Sycp2*$^{+/+}$, *Sycp1*$^{-/-}$ *Sycp2*$^{+/+}$ and *Sycp1*$^{-/-}$ *Sycp2*$^{e16/e16}$ | This study | |
| Oligonucleotides | | |
| Sycp2_Fwd: TGCAAGGCTTCTCTGTTCCT | Eurofins Genomics | N/A |
| Sycp2_Rvs: TGCCTTTACAGTGGCTGCTT | Eurofins Genomics | N/A |
| crRNA1: TTATGTCAGACCAGGCTTAG | Integrated DNA Technologies | N/A |
| crRNA2: CATGTACTTTTTGATGCAAG | Integrated DNA Technologies | N/A |
| crRNA3: TTTTGATGCAAGTGGATCAC | Integrated DNA Technologies | N/A |
| crRNA4: AACTGAAATACCTTGTTTTC | Integrated DNA Technologies | N/A |
| crRNA5: GCAGAATTTGCTTACAAACA | Integrated DNA Technologies | N/A |
| crRNA6: AGGTGTAAACAGTTGACCTA | Integrated DNA Technologies | N/A |
| Trans-activating CRISPR RNA | Integrated DNA Technologies | N/A |
| To amplify C-terminus of SYCP2, SYCP2C_LICfwd: CACCACCACCACAGGGTATCAGG TCCCAGTCAACGTGG | Eurofins Genomics | N/A |
| To amplify C-terminus of SYCP2, SYCP2C_LICrvs: TGAGGAGAAGGCGCGTCATGCAT CATTTCCTTCATGAGCC | Eurofins Genomics | N/A |

| | | |
|---|---|---|
| To amplify C-terminus of H1t, H1tcterm_fwd: ATCACCACCACCACA GG CTCAGTAAGAAGGCGGCTTCTG | Eurofins Genomics | N/A |
| To amplify C-terminus of H1t, H1tcterm_rvs: TGAGGAGAAGGCGCGTCACTTCC TCCCTGCTGCC | Eurofins Genomics | N/A |
| Software and algorithms | | |
| ImageJ | [32] | |
| Fiji Suite for ImageJ | [33] | |
| Adobe Photoshop CC 19 | Adobe | |
| Cell Ranger V6.1 | [34] | |
| R | [35] | |
| Seurat v5 | [36] | |
| scCustomize | [37] | |
| Ggplo2 | [38] | |
| ImerTest Package | [39] | |
| Lme4 package | [40] | |
| CHOPCHOP version 2 | [41,42] | |
| GT-Scan | [43] | |
| CFX Maestro software | | Cat# 12013758 |
| Alphafold 3 | [44] | |
| Other | | |
| salmon sperm | Thermo Fisher Scientific | Cat# AM9680 |

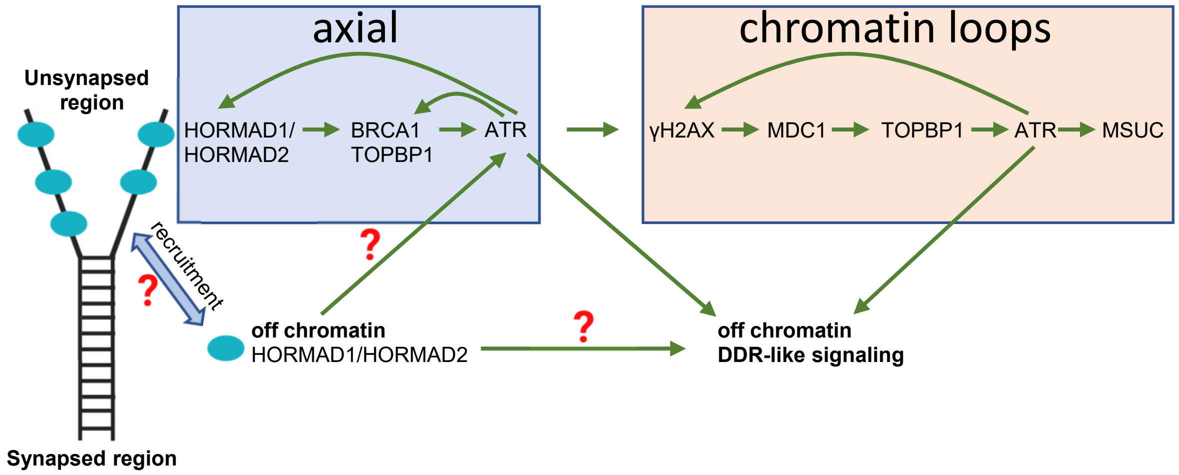

**Supplementary Figure 1. Current model and outstanding questions regarding the molecular network underlying asynaptic axis–triggered ATR activation and the resultant prophase checkpoint in mammalian meiosis.**

Green arrows indicate promoting effects; the blue double-headed arrow denotes inferred but undefined physical interactions that enable recruitment of HORMAD1 and HORMAD2 to unsynapsed axes. The prevailing model of the synapsis checkpoint proposes that preferential enrichment of HORMAD1 and HORMAD2 on unsynapsed chromosome axes forms the basis of asynapsis sensing [1–7]. Axis-bound HORMADs are thought to establish an ATR-mediated signalling cascade that is reinforced by positive feedback and feedforward mechanisms [8–18], resulting in robust ATR activity on both axes and associated chromatin loops within unsynapsed regions. This signalling drives meiotic silencing of unsynapsed chromatin (MSUC) and DNA damage response (DDR)–like signalling, both of which may contribute to the checkpoint mechanisms that eliminate persistently asynaptic meiocytes [8–10,13,15,17–23]. Key outstanding questions include: How are HORMAD1 and HORMAD2 recruited to chromosome axes? What is the functional significance of their axial localization? And do soluble, off-axis pools of HORMADs contribute to ATR activation and DDR-like responses during asynaptic meiosis?

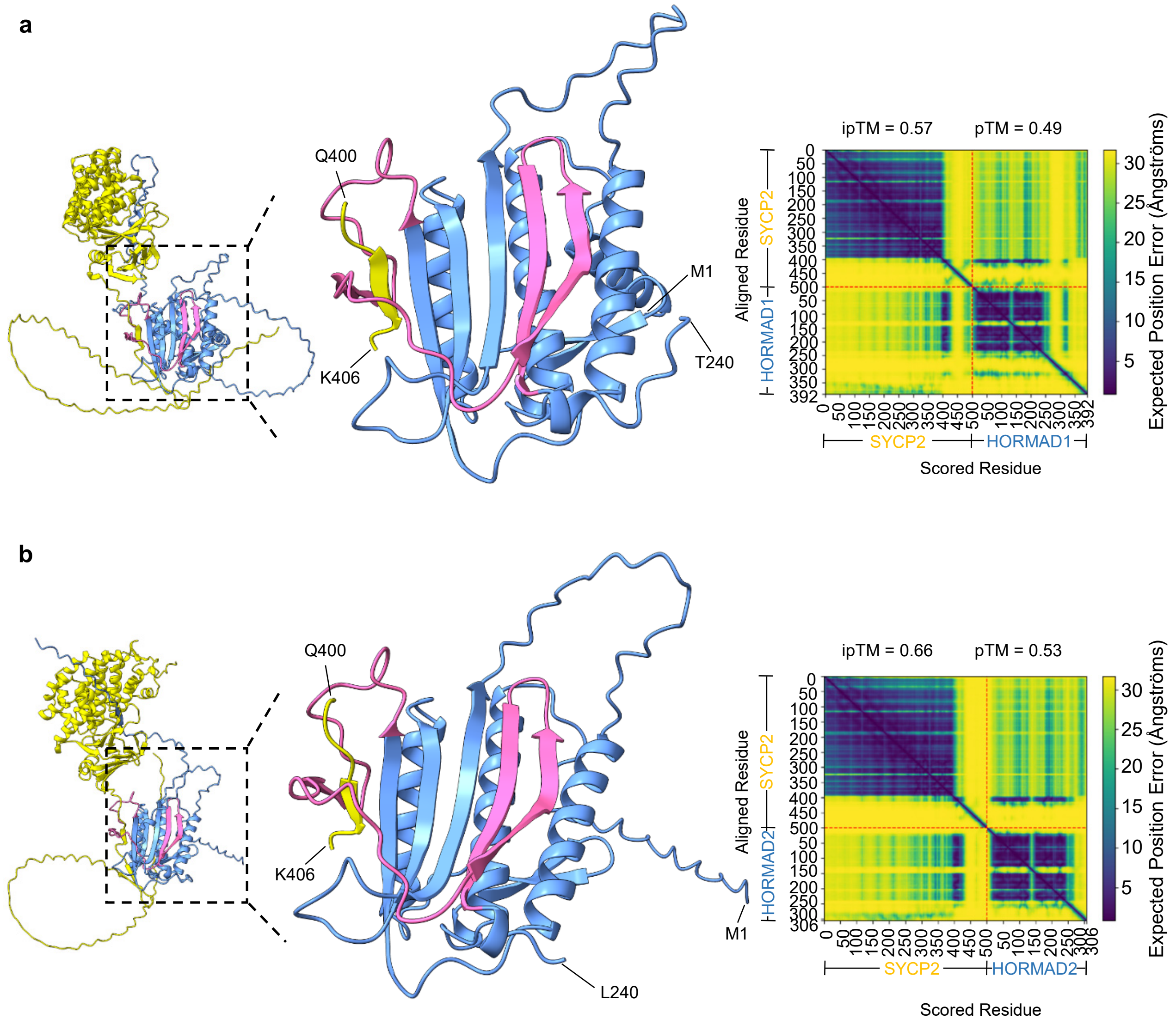

**Supplementary Figure 2. AlphaFold 3 models for HORMAD1 and HORMAD2 interactions with a predicted closure motif in SYCP2.**

**a-b** AlphaFold 3 model predictions of complexes between a fragment of mouse SYCP2 (amino acids (aa) 1-500) and full-length HORMAD1 (aa 1-392) **(a)** or HORMAD2 (aa 1-306) **(b)**. Colors: SYCP2 is yellow, HORMA domain safety belts (HORMAD1, aa 180-230; HORMAD2, aa 184-235) are pink, and the remaining HORMAD regions are blue. Insets of the full models (far left) are enlarged (middle), showing the predicted closure motif of SYCP2 (aa 400-406) wrapped by the safety belts on the surface of the HORMA domains. Right panels show predicted aligned error (PAE) plots with ipTM and pTM scores.

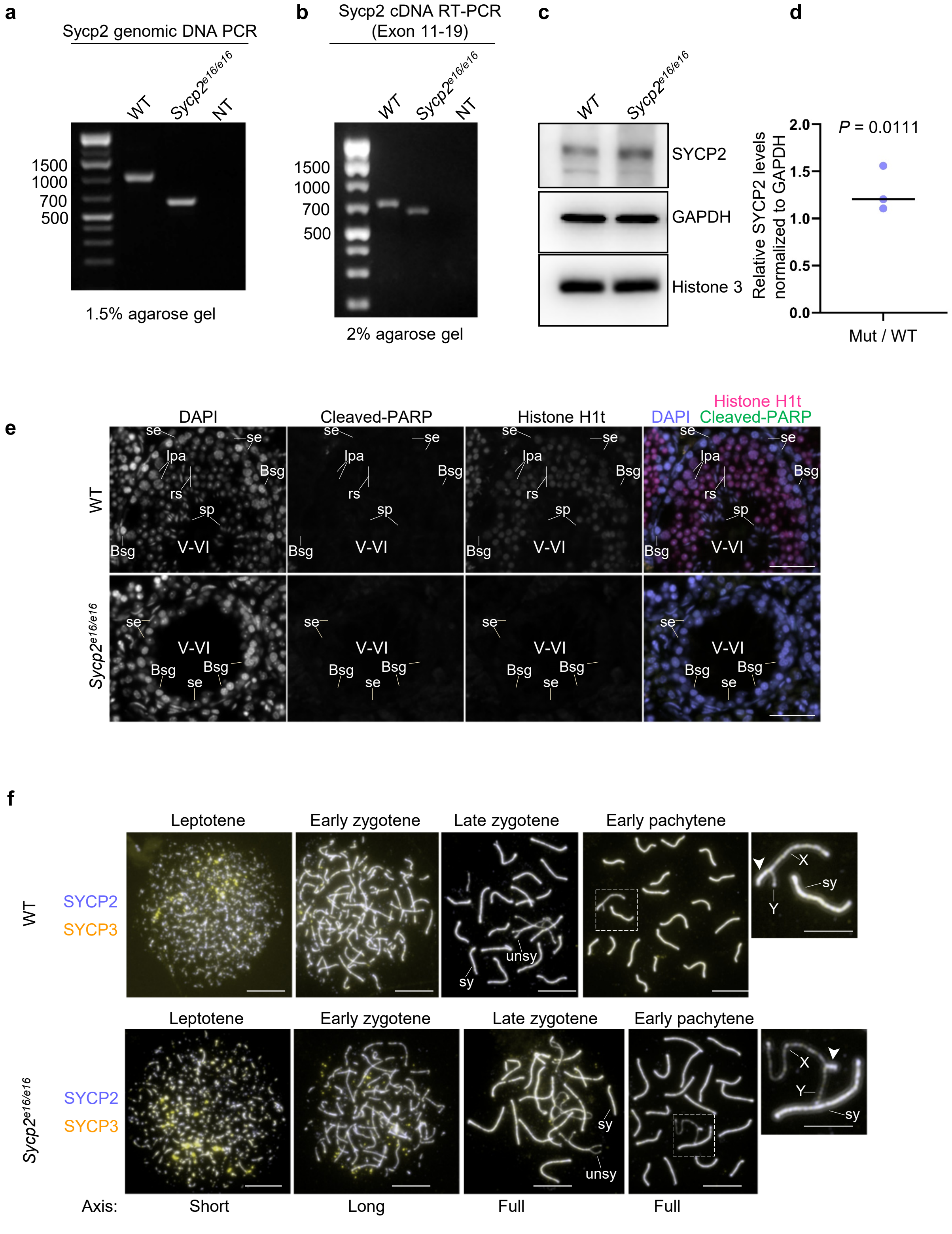

**Supplementary Figure 3. Spermatogenic failure despite efficient axis formation in *Sycp2$^{e16/e16}$*.**

**a** Agarose gel electrophoresis of PCRs using genomic DNA as a template and primers annealing to genomic loci flanking the 16th exon of *Sycp2*. Predicted PCR product sizes: 1057 bp in WT, and 643 bp in *Sycp2$^{e16/e16}$*. **b** Agarose gel electrophoresis of reverse transcription PCRs using total testis RNA as a template from 13 days post-partum (dpp) mice, amplifying sequences between the 11th and 19th exons of *Sycp2*. Predicted PCR product sizes: 713 bp in WT, and 638 bp in *Sycp2$^{e16/e16}$*. **a-b** No-template PCRs (NT) are shown. **c** SDS-PAGE immunoblots of total protein extracts from testes of WT and *Sycp2$^{e16/e16}$* mice at 13 dpp. GAPDH (cytoplasmic marker) and histone H3 (chromatin marker) serve as loading controls. **d** Quantification of total SYCP2 protein abundance in testes of 13 dpp mice. SYCP2 immunoblot signals from testis extracts of *Sycp2$^{e16/e16}$* mice were normalized to corresponding signals from wild-type controls. Bar indicates mean = 1.29 from three experiments; two-tailed one-sample *t*-test, $P$ = 0.0111. **e** DNA staining (DAPI) and immunostaining of cleaved PARP and histone H1t in testis cryosections from adult mice, showing seminiferous tubules in epithelial cycle stages V-VI (see Methods for staging). The images show the presence of pachytene spermatocytes and post-meiotic cells in the WT testis section and their absence in the *Sycp2$^{e16/e16}$* testis section, consistent with spermatocyte elimination at epithelial cycle stage IV in the mutant, as shown in Fig. 1d. Sertoli cells (Se), type B spermatogonia (Bsg), late pachytene spermatocytes (Lpa), round spermatids (rs), and sperm (sp) are marked. Scale bars, 50 µm. **f** Immunostained nuclear spreads of spermatocytes from adult mice. Morphology categories for axes (relevant to Fig. 1e) indicated at the bottom of the panel characterize the prophase stages indicated above each image. Unsynapsed (unsy) and synapsed (sy) regions of autosomes are marked in late zygotene. X and Y chromosomes and the pseudoautosomal region (PAR; white arrowheads) are marked in enlarged insets of early pachytene images. Scale bars, 10 µm (cell) and 5 µm (insets). Source data are provided as a Source Data file.

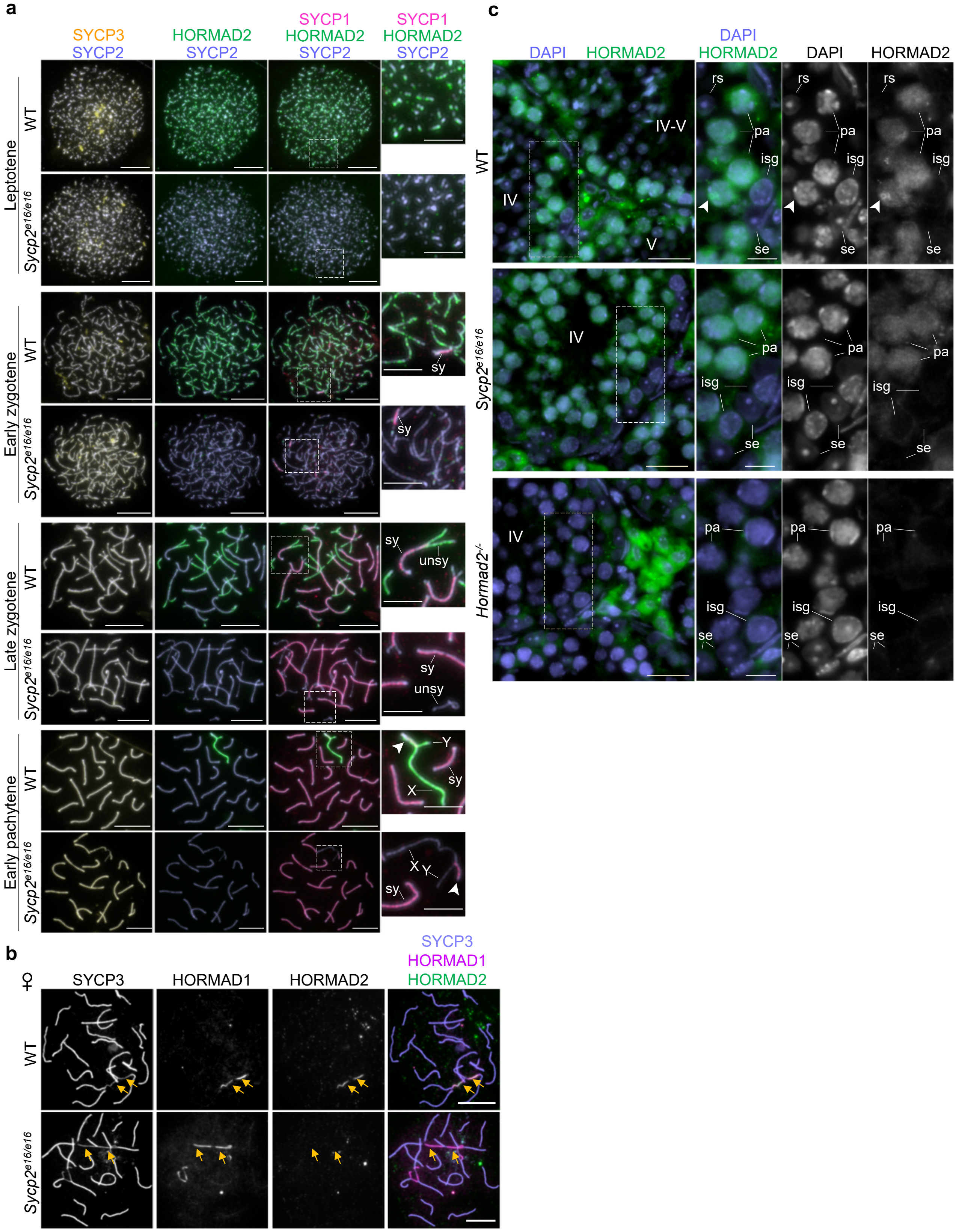

**Supplementary Figure 4. HORMAD2 is present in *Sycp2$^{e16/e16}$* spermatocytes but depleted from the chromosome axes.**

**a-b** Immunostaining in nuclear spreads of spermatocytes **(a)** from adult mice or oocytes **(b)** from 17 dpc foetuses. Scale bars, 10 µm (cell) and 5 µm (inset). Images with matched exposure and levelling are shown within each stage. The channels were differentially levelled between different stages to optimize viewing. **a** X and Y chromosomes, the pseudoautosomal region (PAR; white arrowheads), and examples of synapsed (sy) and unsynapsed (unsy) regions of autosomes are marked in enlarged insets. **b** Unsynapsed axes are marked by yellow arrows**. c** DNA staining (DAPI) and immunostaining of HORMAD2 in cryosections of testes from adult mice. Stages of the seminiferous epithelial cycle are indicated (see 'Methods' for staging). Sertoli cells (se), intermediate spermatogonia (isg), pachytene (pa) spermatocytes, and round spermatids (rs), are marked in the insets. White arrowheads mark the sex body, identified by enriched HORMAD2 signal in WT spermatocytes**.** Note the absence of this enrichment despite presence of nuclear HORMAD2 in pachytene spermatocytes in the *Sycp2$^{e16/e16}$* testis. Scale bars, 25 µm (section) and 10 µm (inset).

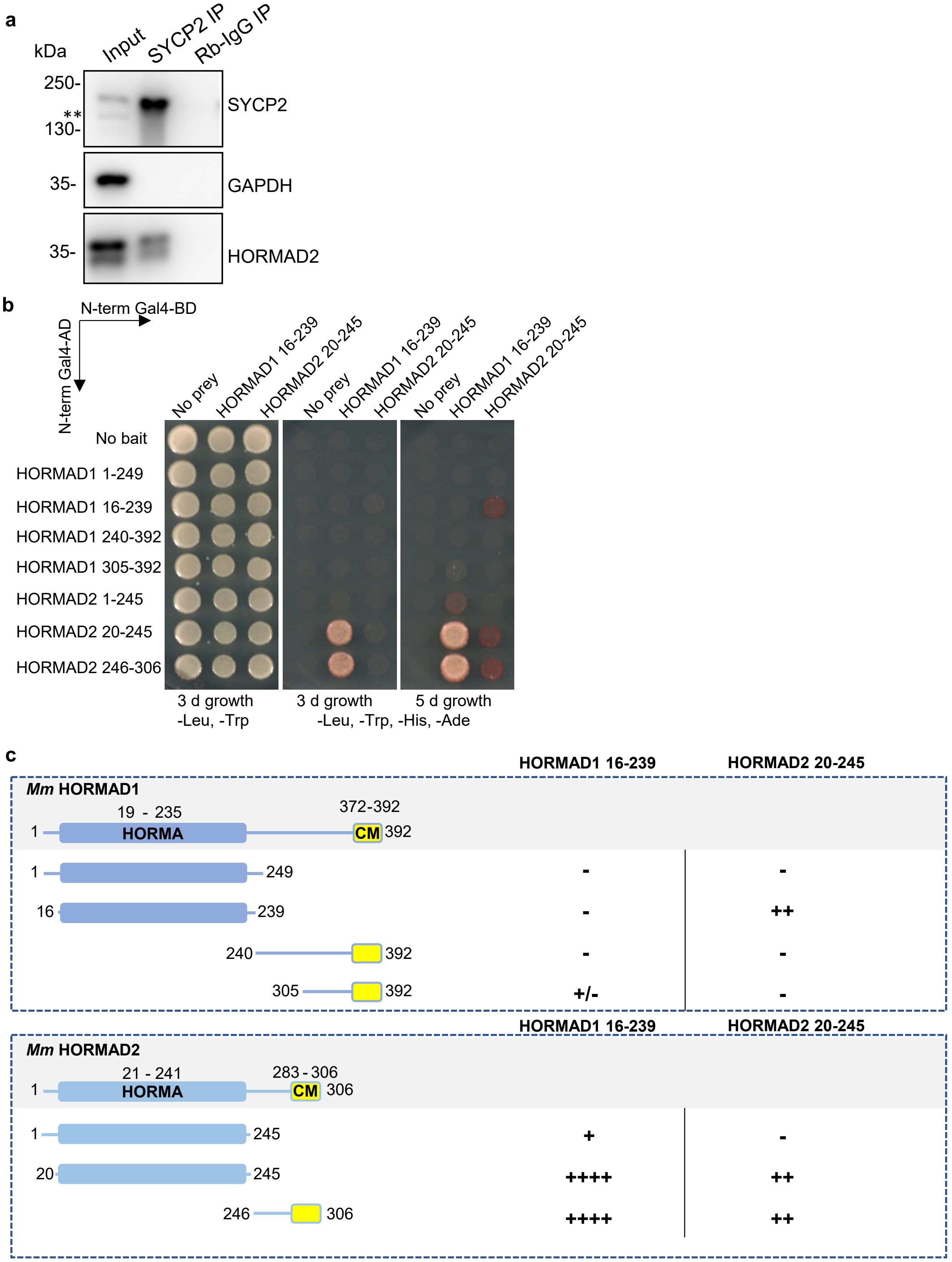

**Supplementary Figure 5. Y2H interactions between HORMADs and complex formation between SYCP2 and HORMAD2.**

**a** SDS-PAGE immunoblot analysis of protein extracts from testes of WT juvenile mice (11 dpp). Total lysate (Input), and immunoprecipitates with anti-SYCP2 (SYCP2 IP) or non-specific rabbit IgG (Rb-IgG IP) antibodies are shown. Asterisks (**) mark an aspecific band that is observed in total protein extracts when analysed in 10% PAGE (see also Fig. 2d-e) but undetectable in the immunoprecipitated product. **b** Yeast two-hybrid interaction assays testing interactions between fragments corresponding to distinct domains of HORMAD1 and HORMAD2. Amino acid (aa) positions of fragment ends are indicated. Yeast cultures are shown after 3 and 5 days of growth on dropout plates. For negative control, proteins of interest were tested in transformations where either the Gal4-binding domain (Gal4-BD) or the Gal4-activation domain (Gal4-AD) vectors were empty. **c** Schematics of HORMAD1 and HORMAD2 domain structures and summary of Y2H interactions between their HORMA domains and depicted protein fragments. Boxes mark the positions of HORMA domains and predicted closure motifs (CMs) as previously described [24]. Numbers represent amino acid positions. Strength of Y2H interactions is graded from no interaction (−) to very strong interaction (++++).

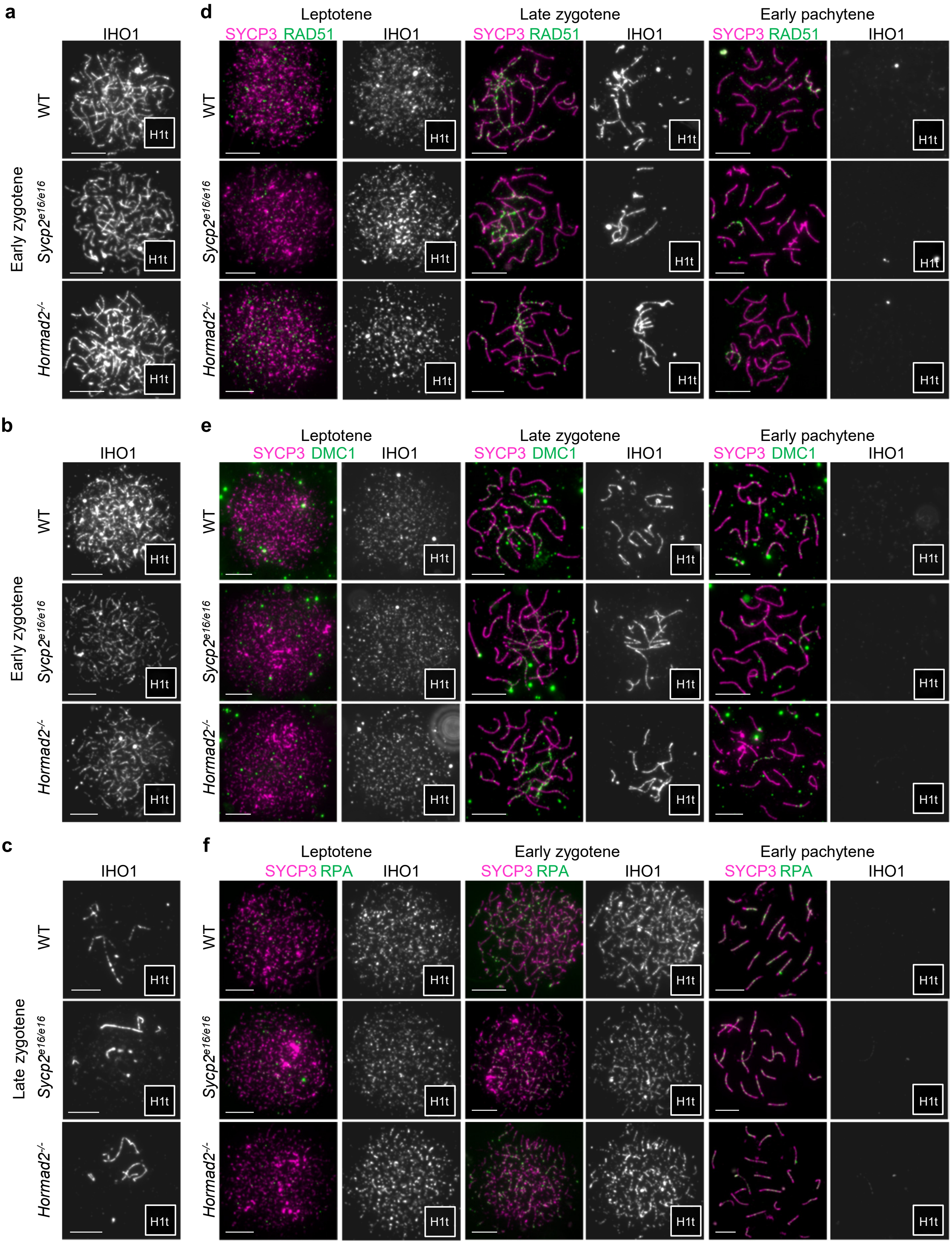

**Supplementary Figure 6. Recombination foci in WT, *Sycp2*$^{e16/e16}$ and *Hormad2*$^{-/-}$ spermatocytes.**

Immunostaining of nuclear surface spreads of spermatocytes from adult mice. For recombination markers (RAD51, DMC1, and RPA), images are shown with matched exposure and levelling across all genotypes and prophase stages (**d-f**). For stage markers (IHO1 and histone H1t), exposure and levelling are matched across genotypes for late zygotene and early pachytene; at other stages, exposure and levelling are matched across genotypes but differentially adjusted between stages for optimal viewing. SYCP3 was differentially levelled to optimize visualization. Histone H1t is shown in miniaturized images in the bottom right corners of full-size cell images. Panels **a-c** show IHO1 and histone H1t images in cells corresponding to Fig. 3d-f. Scale bars, 10 µm.

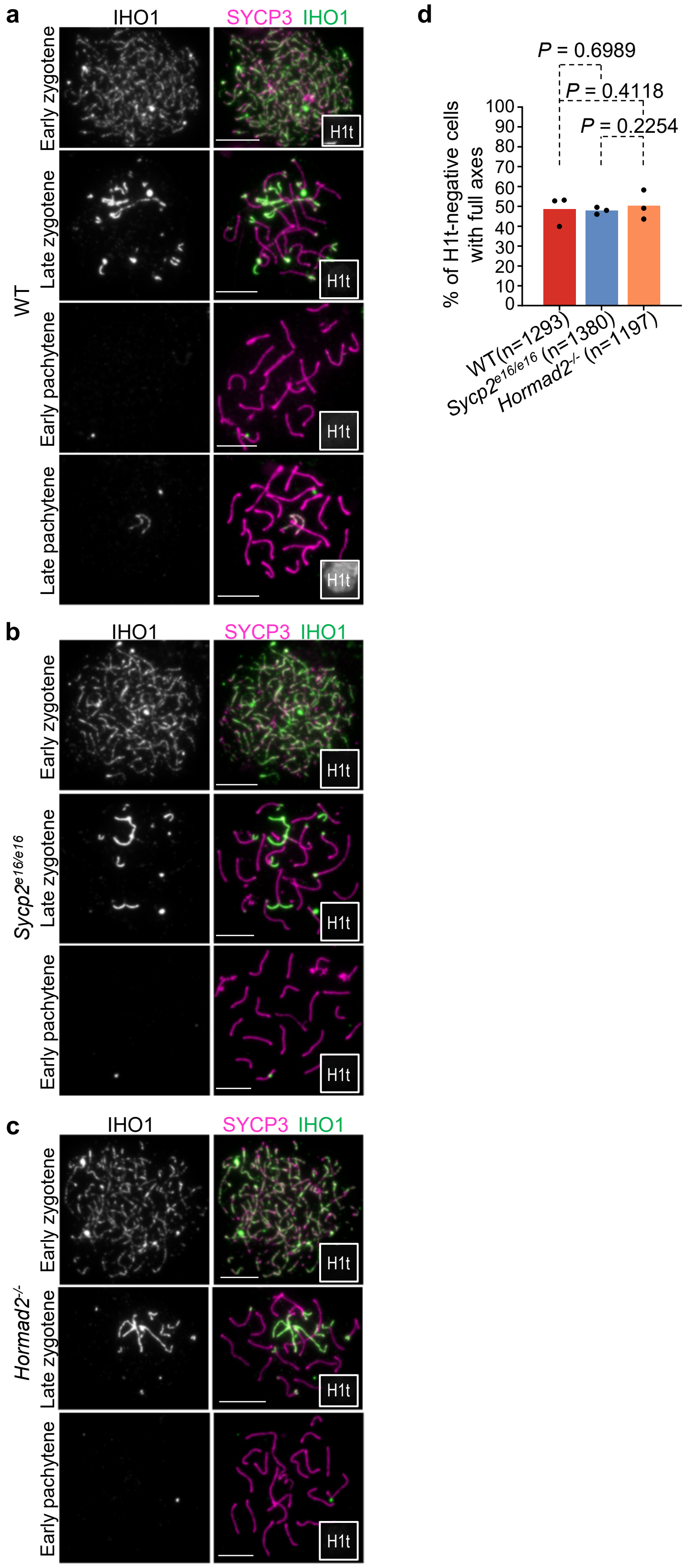

**Supplementary Figure 7. IHO1 localization in WT, *Sycp2$^{e16/e16}$* and *Hormad2$^{-/-}$*spermatocytes.**

**a-c** Immunostaining of nuclear surface spreads of spermatocytes from adult mice. Images with matched exposure and levelling are shown for stage markers IHO1 and histone H1t (miniaturized images in bottom right corner of full-size cell images) across genotypes and prophase stages. To illustrate IHO1 dynamics, early zygotene, late zygotene, and early pachytene are compared between WT **(a)**, *Sycp2*$^{e16/e16}$ **(b)**, and *Hormad2*$^{-/-}$ **(c)** genotypes; late pachytene is only shown in WT **(c)** because the mutants lack spermatocytes beyond mid-pachytene. SYCP3 was differentially levelled to optimize viewing. Scale bars, 10 µm. **d** Quantification of IHO1 localization on axes of spermatocytes that have fully developed axes and are negative for histone H1t (corresponding to late zygotene and early pachytene in wild-type spermatocytes). Graph shows data points and weighted averages (bars) of percentages of IHO1-positive spermatocytes (WT = 48.60%, *Sycp2*$^{e16/e16}$ = 47.90%, and *Hormad2*$^{-/-}$ = 50.33%) from three biological replicates. Number of analyzed cells (*n*) per genotype is indicated. Likelihood-ratio test (chi-squared distribution) was used to determine whether the proportion of IHO1-positive cells among histone H1t-negative, fully-axis-displaying spermatocytes differed significantly between mutants and wild type. Source data are provided as a Source Data file.

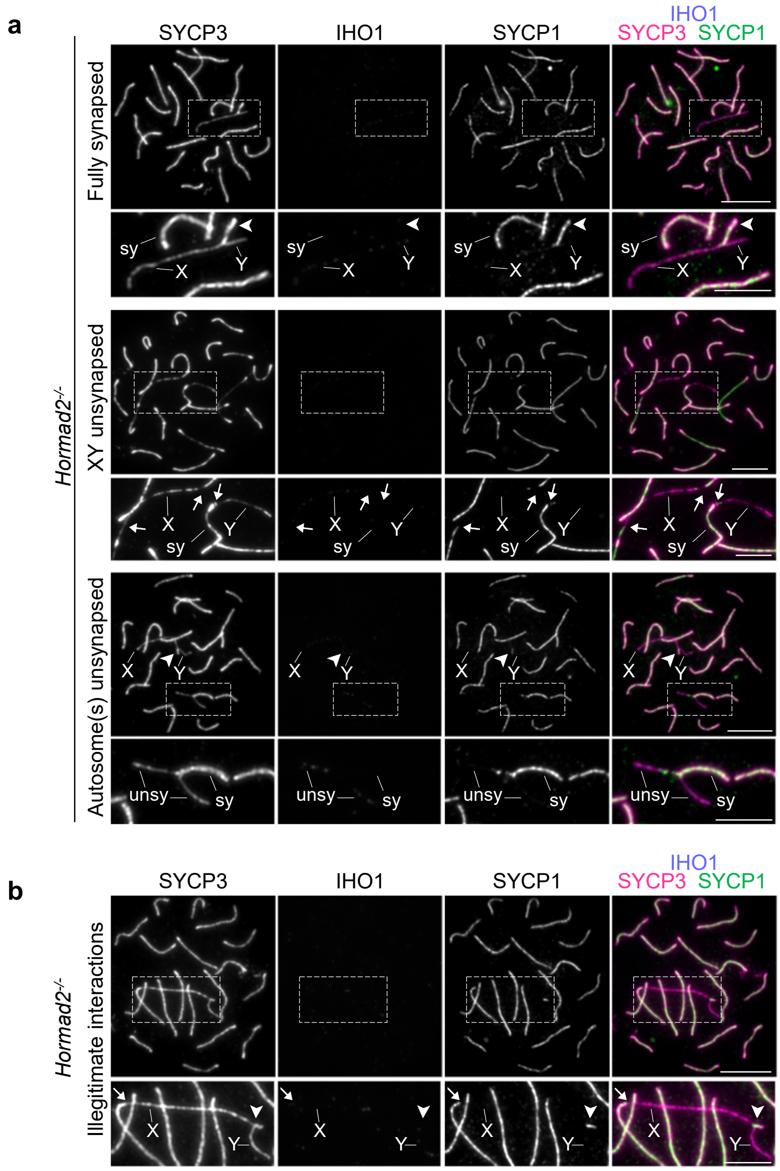

**Supplementary Figure 8.** ***Hormad2**$^{-/-}$* **spermatocytes display mild synapsis defects.**

**a-b** Immunostained nuclear spreads of early pachytene *Hormad2*$^{-/-}$ spermatocytes from adult mice illustrating synapsis categories quantified in Fig. 4c,d. SYCP1 marks synapsed regions; fully formed axes (SYCP3) and the absence of IHO1 identify the cells as early pachytene. Pseudoautosomal regions (PARs; white arrowheads in **a-b**), synapsed (sy) and unsynapsed (unsy) autosomal regions (**a**), and illegitimate interactions between the ends of the X or Y chromosomes and fully synapsed autosomes (white arrows in **a-b**) are indicated. Scale bars, 10 µm (whole cell) and 5 µm (insets).

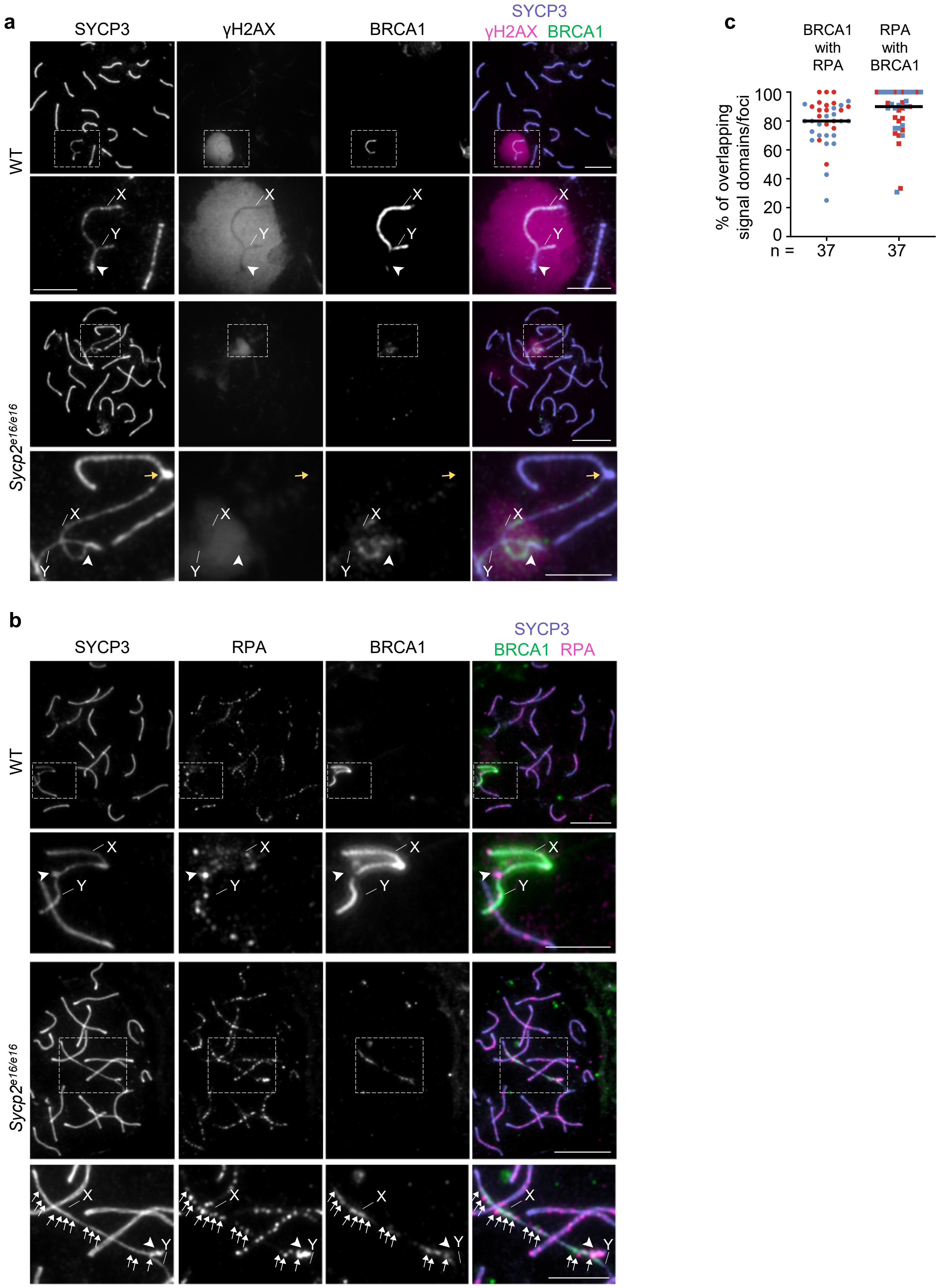

**Supplementary Figure 9. The 16th exon of *Sycp2* is required for efficient BRCA1 accumulation on sex chromosomes.**

**a-b** Immunostained nuclear surface spreads of early pachytene spermatocytes from adult mice. Images with matched exposure and levelling are shown for γH2AX, RPA and BRCA1 signals. X and Y chromosomes, PARs (white arrowheads in **a-b**), an illegitimate end-to-end association between the X chromosome and an autosome (yellow arrow in **a**), and sites of sex-chromosome-axis-associated RPA foci (arrows in **b**) are marked in enlarged insets. Scale bars, 10 µm (cells), 5 µm (insets). **c** Quantification of overlap between axis-associated RPA and BRCA1 signals on sex chromosomes in *Sycp2*$^{e16/e16}$ spermatocytes. The numbers of analysed cells (n) correspond to two experiments (differentiated by blue and red colors); medians (bars) are 80% (BRCA1 colocalizing with RPA), and 90% (RPA colocalizing with BRCA1).

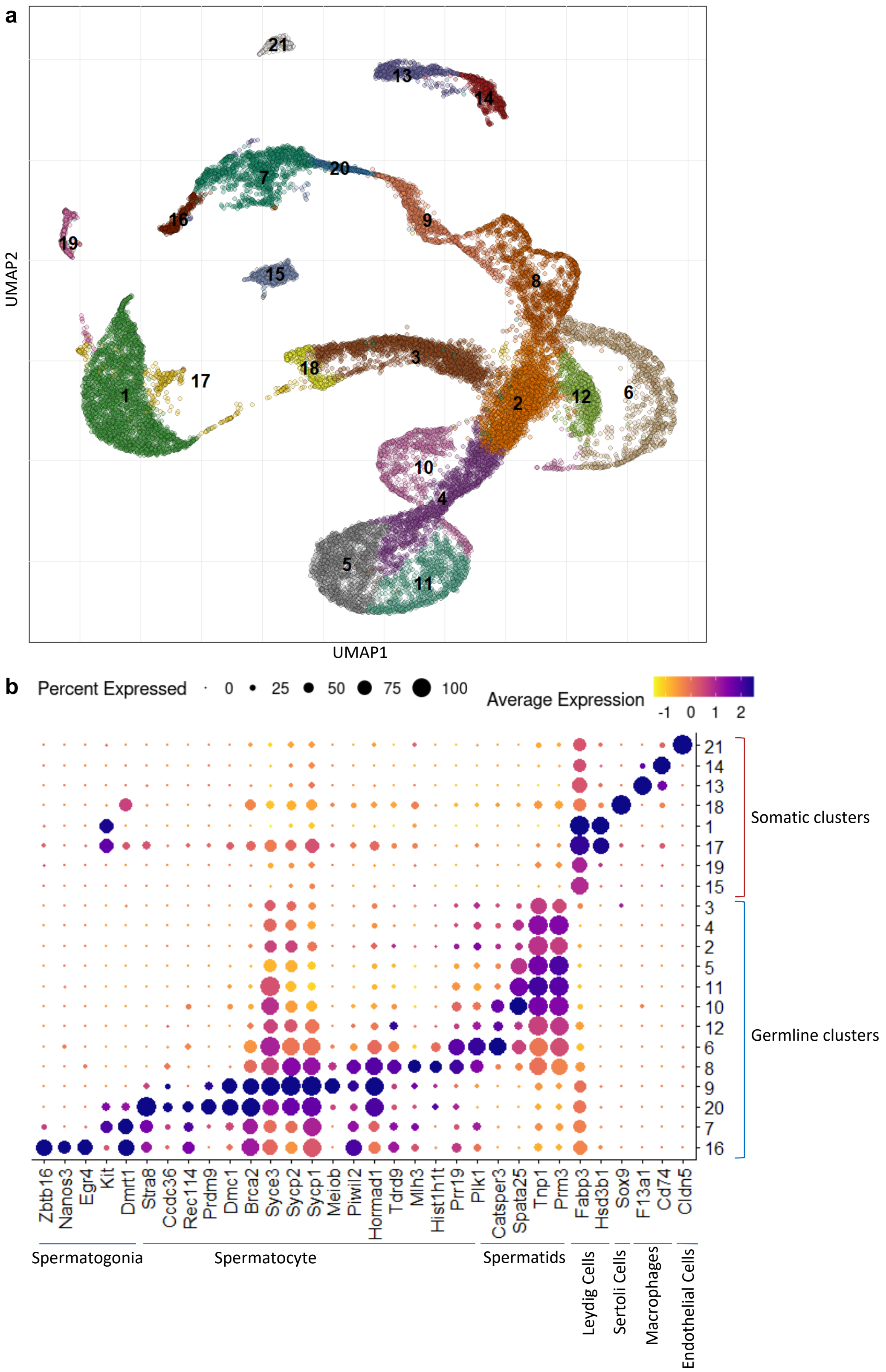

**Supplementary Figure 10. Unsupervised clustering of pooled scRNAseq data from testes of WT, *Sycp2*$^{e16/e16}$, and *Hormad2*$^{-/-}$ mice.**

**a** UMAP visualization of whole testis scRNA-seq data, showing cell populations identified by unsupervised Seurat-based clustering (cluster resolution parameter = 0.5). Pooled data are shown from testes of WT (four samples from two biological replicates), *Sycp2*$^{e16/e16}$ (two biological replicates), *Hormad2*$^{-/-}$ (two biological replicates) mice. The numbers represent clusters of distinct testicular cell populations as listed in **b**. **b** Dot plot heatmaps showing expression of selected marker genes. The x-axis lists representative marker genes and the cell types that primarily express them; the y-axis lists cluster numbers corresponding to **a**. Expression levels are represented as z-scores ranging from –1 to 2. The size and color intensity of each dot indicate the proportion of cells expressing the gene and the relative expression level, respectively. The full list of marker genes used for identifying spermatogenic and somatic clusters, and the distinguishing characteristics of the cell clusters are listed in Supplementary Table 3 and 4.

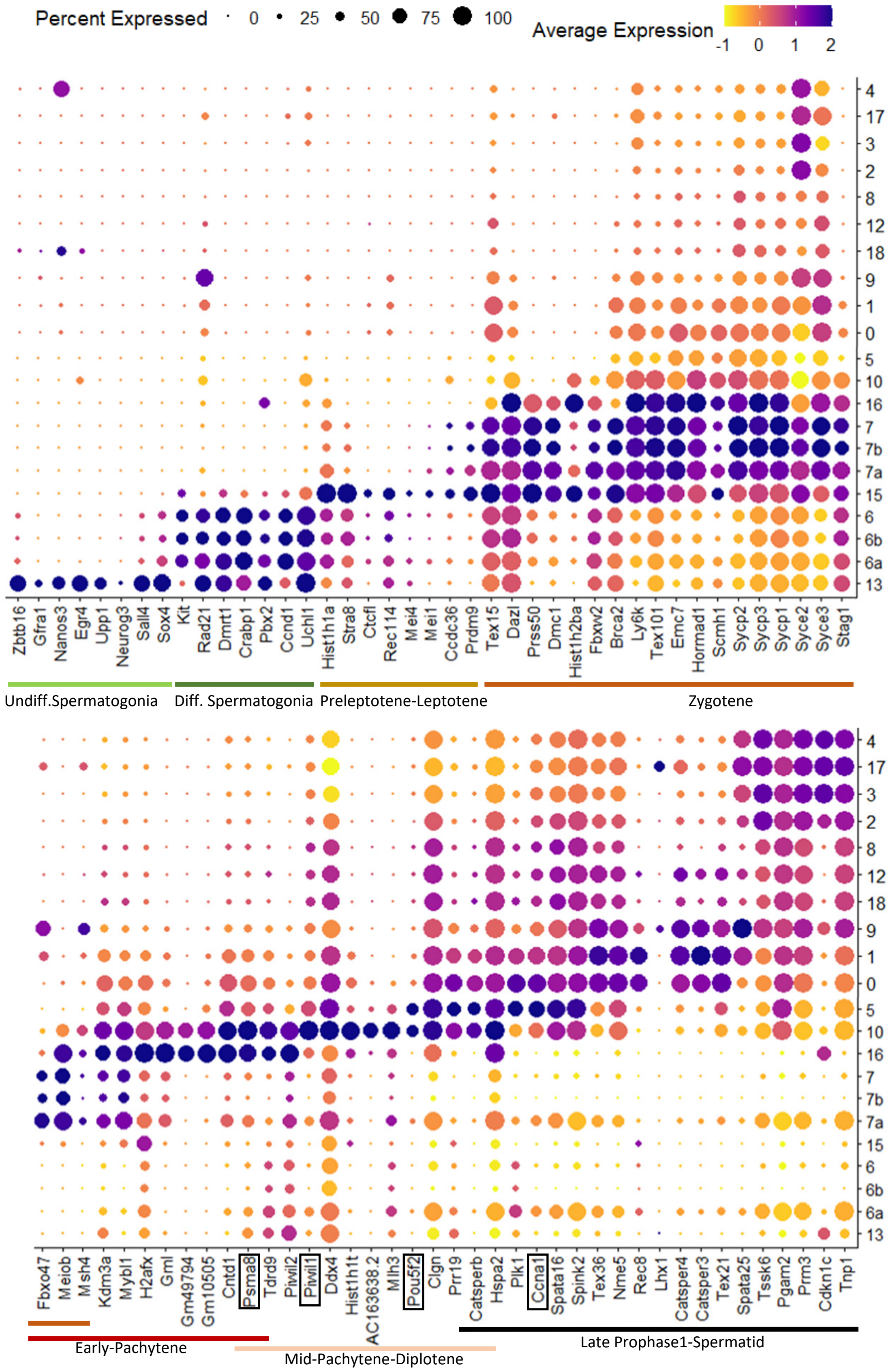

**Supplementary Figure 11. Expression of marker genes in spermatogenic populations identified by supervised clustering.**

Dot plot showing scRNA-seq–derived expression levels of all the marker genes used for supervised clustering and subsequent annotation of spermatogenic cell populations. Genes used exclusively for cell-type annotation are boxed. The x-axis lists marker genes; the y-axis lists cluster numbers corresponding to those in Fig. 6a–b. Clusters 6 and 7 are shown both as their constituent subclusters (6a/6b and 7a/7b) and as pooled clusters. In both cases, WT cells are preferentially enriched in subcluster "a" and mutant cells in subcluster "b," despite highly similar expression profiles of stage-defining markers. Expression values are represented as z-scores ranging from –1 to 2. Dot size and color intensity indicate the proportion of cells expressing each gene and the relative expression level, respectively. Colored bars below the x-axis denote the cell populations predominantly expressing the corresponding markers. A complete list of marker genes used for clustering and their defining characteristics is provided in Supplementary Tables 3 and 5.

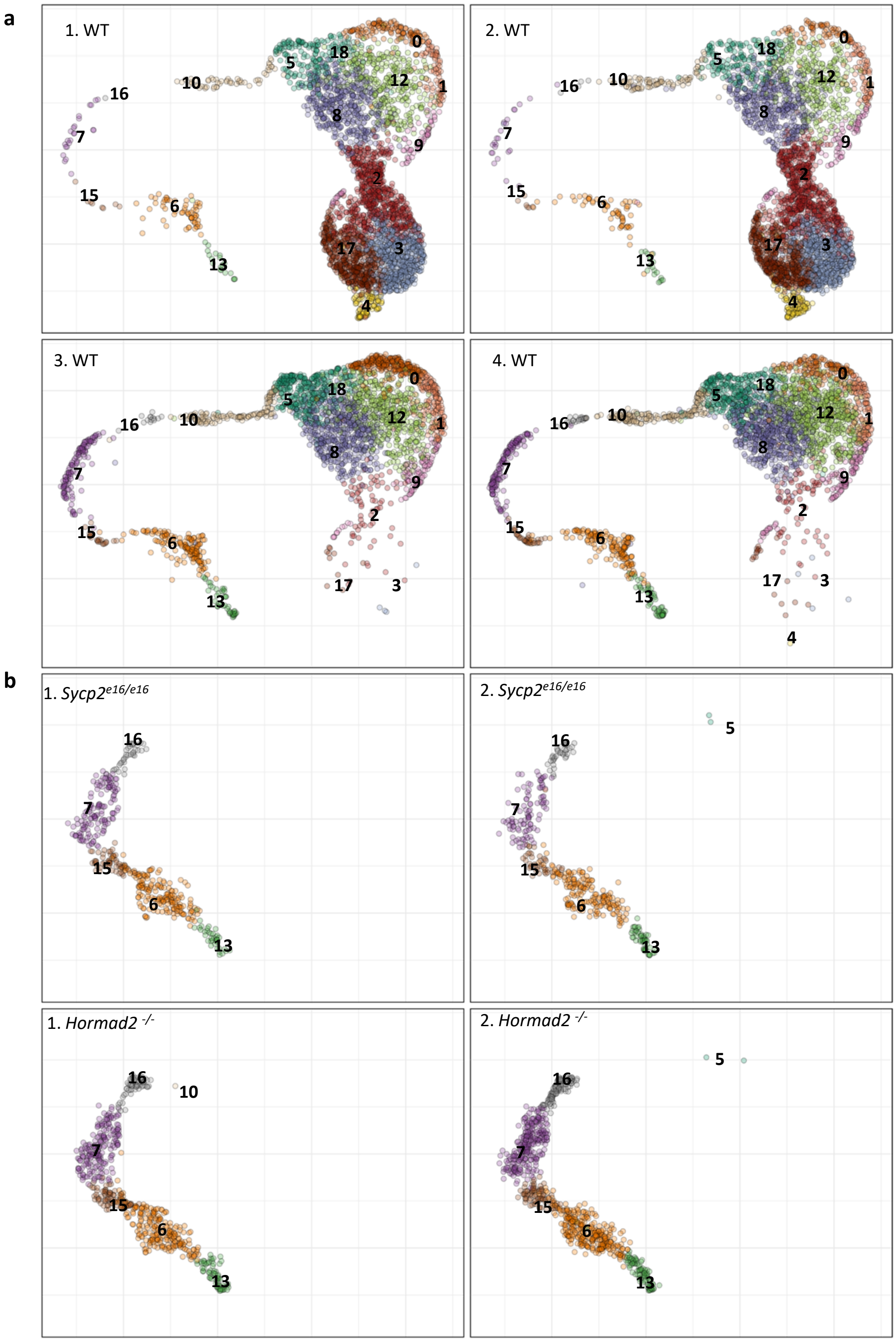

**Supplementary Figure 12. scRNA-seq UMAPs of spermatogenic populations from individual WT, *Sycp2*$^{e16/e16}$, and *Hormad2*$^{-/-}$ samples.**

UMAP visualization of single-cell RNA sequencing data showing spermatogenic cell populations identified by supervised clustering using germline marker genes listed in Supplementary Fig. 10 (see also methods). Cluster numbers correspond to spermatogenic stages listed in Fig. 6b. a UMAPs for two biological replicates (left and right columns) of WT samples processed either without (top row) or with (bottom row) sperm and dead cell depletion. UMAPs for two biological replicates (left and right columns) of *Sycp2*$^{e16/e16}$ (top row), and *Hormad2* $^{-/-}$ (bottom row) samples.

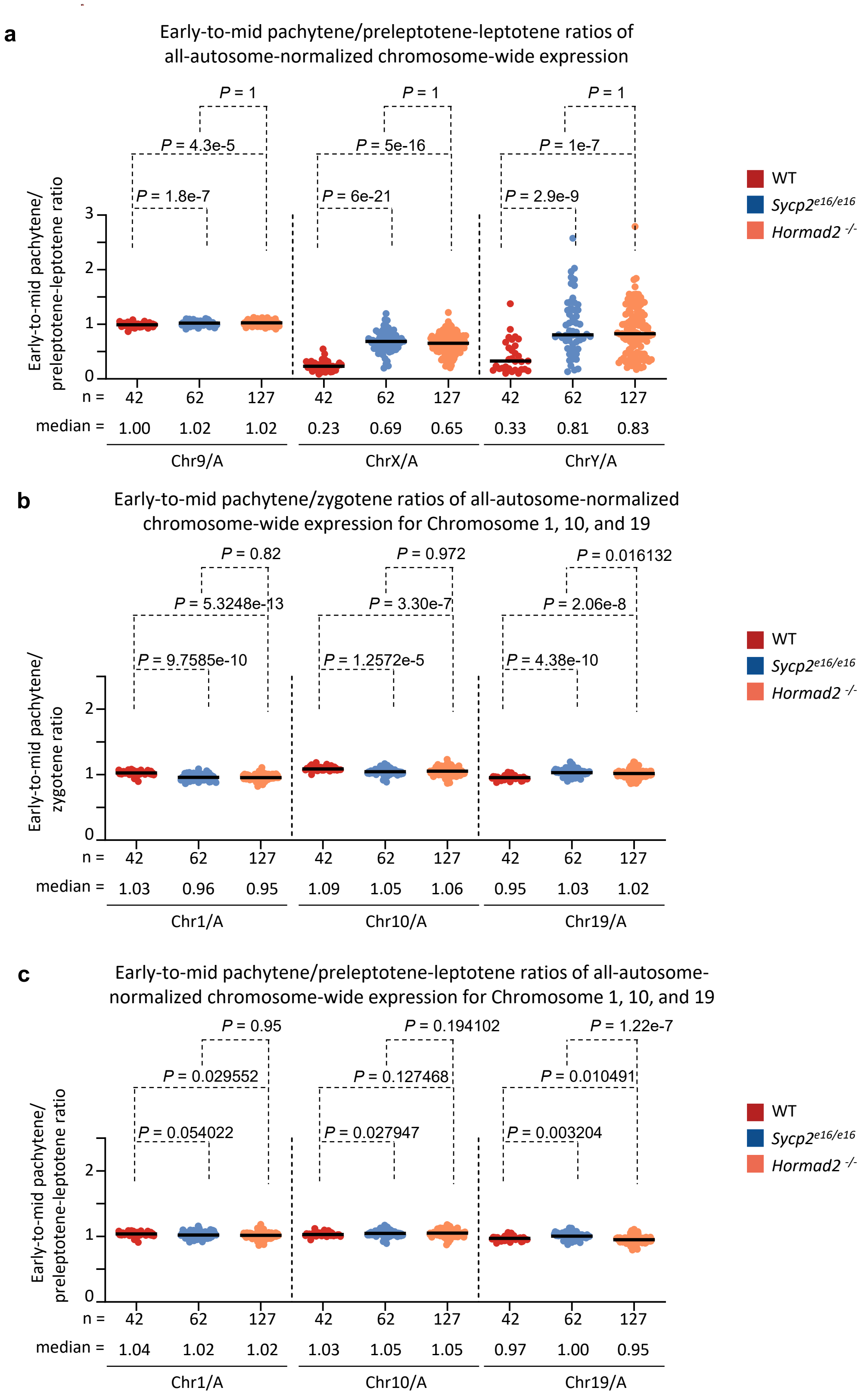

**Supplementary Figure 13. scRNA-seq reveals impaired sex chromosome silencing in *Hormad2*$^{-/-}$ and *Sycp2*$^{e16/e16}$ spermatocytes.**

**a** All-autosome-normalized chromosome-wide expression in early-to-mid pachytene cells relative to preleptotene-leptotene stages. Each data point represents normalized expression for chromosome 9 (Chr9/A), the X chromosome (ChrX/A), or the Y chromosome (ChrY/A) in early-mid pachytene cells divided by the median normalized expression of the same chromosome in preleptotene-leptotene cells (see Methods for details). *P* values were determined using the Wilcoxon rank-sum test. **b-c** All-autosome-normalized chromosome-wide expression in early-to-mid pachytene cells relative to zygotene (**b**) and preleptotene-leptotene (**c**) stages. Each data point represents all-autosome-normalized expression for chromosome 1 (Chr1/A), chromosome 10 (Chr10/A), or chromosome 19 (Chr19/A) in individual early-mid pachytene cells divided by the median normalized expression of the same chromosome in zygotene (**b**) or preleptotene-leptotene (**c**) cells. Data points represent individual cells; medians (black bars), and *P* values from the Wilcoxon rank-sum test are indicated. Source data are provided as a Source Data file.

**Supplementary Table 1. Fertility in female *Sycp2* $^{e16/e16}$ and *Hormad2* $^{-/-}$ mice**

Quantification of pup numbers from crosses of wild type (WT) male mice with wild-type, *Sycp2* $^{e16/e16}$ or *Hormad2* $^{-/-}$ female mice. Statistical significances were calculated by two-tailed unpaired t-test with Welch correction.

| female age | female < 35 weeks | | | female >35 weeks | | |
|---|---|---|---|---|---|---|
| female genotype | WT | *Sycp2* $^{e16/e16}$ | Hormad2$^{-/-}$ | WT | *Sycp2* $^{e16/e16}$ | Hormad2$^{-/-}$ |
| male genotype | WT | | | | | |
| breeding pairs | 4 | 4 | 4 | 4 | 4 | 4 |
| total breeding weeks | 91 | 93 | 96 | 61 | 55 | 51 |
| pups/breeding week | 1.759850823 | 1.628573834 | 1.526694757 | 0.802421534 | 0.895231729 | 0.944023638 |
| Comparison with WT (P values from unpaired t-test with Welch's correction) | x | 0.6979 | 0.2595 | x | 0.6699 | 0.5453 |

**Supplementary Table 2. Summary of single-cell RNA-seq quality metrics in wild-type and mutant testes samples.** Key sequencing and mapping statistics for four wild-type (WT), two *Hormad2* $^{-/-}$, and two *Sycp2* $^{e16/e16}$ samples. Two biological replicates of WT samples were processed either without (1 and 2) or with (3 and 4) sperm and dead cell depletion.

| | | Samples | | | | | | | |
|---|---|---|---|---|---|---|---|---|---|
| | | **1. WT** | **2. WT** | **3. WT** | **4. WT** | **1. *Hormad2*** $^{-/-}$ | **2. *Hormad2*** $^{-/-}$ | **1. *Sycp2*** $^{e16/e16}$ | **2. *Sycp2*** $^{e16/e16}$ |
| Metric | ID | CMO309_WT (no dead cell removal) | CMO312_WT (no dead cell removal) | CMO310_WT (with dead cell removal) | CMO311_WT (with dead cell removal) | CM0309 (no dead cell removal) | CM0310 (no dead cell removal) | CM0311 (no dead cell removal) | CM0312 (no dead cell removal) |
| | Number of sequencing rounds | 1 | 1 | 1 | 1 | 2 | 2 | 2 | 2 |
| | Cells numbers before quality control | 3860 | 4089 | 3678 | 4118 | 3163 | 3048 | 2784 | 2796 |
| | Confidently mapped antisense | 1.24% | 1.26% | 1.43% | 1.50% | 1.32% | 1.46% | 1.28% | 1.35% |
| | Confidently mapped to exonic regions | 79.87% | 79.59% | 72.50% | 70.05% | 75.91% | 71.88% | 77.82% | 75.60% |
| | Confidently mapped to genome | 93.27% | 93.28% | 92.20% | 92.05% | 92.21% | 91.64% | 92.46% | 92.35% |
| | Confidently mapped to intergenic regions | 5.92% | 5.82% | 7.16% | 7.58% | 3.27% | 3.80% | 3.03% | 3.30% |
| | Confidently mapped to intronic regions | 7.48% | 7.87% | 12.55% | 14.41% | 13.03% | 15.96% | 11.62% | 13.45% |
| | Confidently mapped to transcriptome | 76.25% | 76.01% | 68.73% | 66.38% | 65.31% | 64.17% | 65.82% | 64.98% |
| | Mapped to genome | 95.87% | 95.86% | 95.46% | 95.31% | 94.83% | 94.46% | 95.03% | 94.94% |
| | Median UMI counts per cell | 14631 | 12706 | 8555 | 6984 | 6956 | 5759 | 7879 | 7337 |
| | Median genes per cell | 3332 | 3230 | 3408 | 3073 | 2640 | 2441 | 2798 | 2700 |
| | Median reads per cell | 39845 | 36440 | 32869 | 28510 | 17692 | 15109 | 19995 | 19117 |
| | Number of reads from cells called | 273,559,970 | 261,150,602 | 292,705,166 | 260,357,842 | 69,938,676 | 59,008,196 | 69,298,650 | 65,718,657 |
| | Total number of genes detected | 26056 | 26009 | 26726 | 26745 | 21811 | 21998 | 21515 | 21973 |
| | Cell numbers in input for unsupervised clustering after quality control | 3703 | 3912 | 3402 | 3837 | 2355 | 2144 | 2162 | 2015 |
| | Cell numbers in spermatogenic populations subjected to supervised clustering | 3439 | 3501 | 2634 | 2872 | 455 | 417 | 643 | 881 |

**Supplementary Table 3. Germline and somatic marker genes used for cluster identification.**
Two sets of marker genes used to identify and separate germline and somatic cell populations in the single-cell RNA-seq data. Somatic markers (top section) were used to identify somatic cell-enriched clusters, which were excluded from further analysis. Germline markers (bottom section) were used to identify spermatogenic clusters and to perform supervised re-clustering of spermatogenic subpopulations.

| Gene Name | Predominant cell types of expression |
|---|---|
| somatic cells | |
| *Hsd3b1* | Leydig cells |
| *Star* | Leydig cells |
| *Fabp3* | Leydig cells |
| *Sncg* | Endothelial cells |
| *Cldn5* | Endothelial cells |
| *Mmrn2* | Endothelial cells |
| *F13a1* | Macrophages |
| *STAB1* | Macrophages |
| *Cd74* | Macrophages |
| *Nr5a1* | Sertoli cells |
| *sox9* | sertoli cells |
| spermatogenic cells | |
| *Zbtb16* | Undifferentiated Spermatogonia |
| *Gfra1* | Undifferentiated Spermatogonia |
| *Nanos3* | Undifferentiated Spermatogonia |
| *Egr4* | Undifferentiated Spermatogonia |
| *Upp1* | Undifferentiated Spermatogonia |
| *Dmrt1* | Undifferentiated Spermatogonia |
| *Neurog3* | Undifferentiated Spermatogonia |
| *Sall4* | Undifferentiated Spermatogonia |
| *Sox4* | Differentiated Spermatogonia |
| *Crabp1* | Differentiated Spermatogonia |
| *Hist1h1a* | Differentiated Spermatogonia |
| *Stra8* | Differentiated Spermatogonia |
| *Pbx2* | Differentiated Spermatogonia |
| *Ccnd1* | Differentiated Spermatogonia |
| *Uchl1* | Spermatogonia |
| *Rad21* | Spermatogonia and Spermatids |
| *Smc1b* | Spermatogonia and Early Prophase1 |
| *Dazl* | Spermatogonia and Early Prophase1 and Late Prophase1 |
| *Prss50* | Ealry Prophase 1 |
| *Tex15* | Ealry Prophase 1 |
| *Hist1h2ba* | Early prophase 1 |
| *Fbxo47* | Early prophase 1 |
| *Ccdc36* | Early prophase 1 |
| *Ctcfl* | Early prophase 1 |
| *Rec114* | Early prophase 1 |
| *Mei1* | Early prophase 1 |
| *Mei4* | Early prophase 1 |
| *Dmc1* | Early prophase 1 |
| *Prdm9* | Early prophase 1 |
| *Fbxw2* | Early prophase 1 |
| *Brca2* | Early prophase 1 |
| *Rec8* | Early prophase 1, Late prophase1, Early Spermatids |
| *Ly6k* | Early prophase 1 |
| *Sycp3* | Early prophase 1 |
| *Sycp1* | Early prophase 1 |
| *Tex101* | Early prophase 1 |

| | |
|---|---|
| *Emc7* | Early prophase 1 |
| *Syce2* | Early prophase1 and Late Spermatids |
| *Syce3* | Early prophase 1 |
| *Hormad1* | Early Prophase 1 |
| *H2afx* | Early prophase1 |
| *Sycp2* | Early prophase1 |
| *Meiob* | Early prophase1 |
| *Stag1* | Spermatogonia and Early Prophase1 |
| *Gml* | Early Prophase1 |
| *Ddx4* | Late Prophase1, broad from spermatogonia to early spermatids |
| *Plk1* | Late Prophase 1 |
| *Mlh3* | Late Prophase 1 |
| *Clgn* | Late Prophase 1 |
| *Cntd1* | Late Prophase 1 |
| *Prr19* | Late Prophase 1 |
| *Tdrd9* | Late Prophase 1 |
| *Piwil2* | Late Prophase 1 |
| *Scmh1* | Late Prophase 1 |
| *Catsperb* | Late Prophase 1 |
| *Kdm3a* | Late Prophase 1 |
| *Hist1h1t* | Late Prophase 1 |
| *Msh4* | Late Prophase 1 |
| *Spata16* | Late Prophase 1 |
| *Spink2* | Late Prophase 1 |
| *Pgam2* | Late Prophase 1 |
| *Hspa2* | Late Prophase 1 |
| *Mybl1* | Late Prophase 1 |
| *AC163638.2* | Late Prophase1 |
| *Gm49794* | Late Prophase1 |
| *Gm10505* | Late Prophase1 |
| *Nme5* | Early Spermatids |
| *Tex36* | Early Spermatids |
| *Cdkn1c* | Early Spermatids |
| *Spata25* | Early Spermatids |
| *Catsper4* | Early Spermatids |
| *Catsper3* | Early Spermatids |
| *Tex21* | Early Spermatids |
| *Prm3* | Late Spermatids |
| *Tssk6* | Late Spermatids |
| *Tnp1* | Late Spermatids |
| *Lhx1* | Early and Late Spermatids |

**Supplementary Table 4. Summary of unsupervised clusters and marker-based cell-type annotations.**
Clusters identified by unsupervised analysis of single-cell RNA-seq data and annotated based on expression of somatic and germline marker genes. Identified cell types include spermatogonia, spermatocytes, spermatids, Leydig cells, Sertoli cells, macrophages, and endothelial cells. Clusters with germline marker enrichment were subsequently selected for supervised re-clustering (see Supplementary Table 5). Key marker genes identifying dominant cell types in each cluster are listed.

| Cluster numer | Celltype | Selected for supervised clustering | Key marker genes |
|---|---|---|---|
| 13 | Macrophages | NO | Cd74, Stab1, F13a0 — macrophage markers. |
| 14 | Macrophages | NO | Cd74, Stab1, F13a1 — macrophage markers. |
| 21 | Endothelial cells | NO | Cldn5, Mmrn2 — endothelial cell markers. |
| 18 | Sertoli cells | NO | High Sox9 indicating dominant Sertoli cell contribution. Very low levels of spermatogenic markers. |
| 19 | Leydig cells | NO | Fabp3— Leydig cell marker. |
| 17 | Leydig cells | NO | Nr5a1, Star, Fabp3, trace Stra8, Sox4, Uchl1 —mostly Leydig markers, low level early spermatogenic markers. |
| 15 | Leydig cells | NO | Fabp3— Leydig cell marker. |
| 1 | Leydig cells | Yes | Nr5a1, Star, Fabp3 — strong Leydig-specific expression profile. |
| 3 | Mostly spermatid with contribution from late prophase spermatocytes | Yes | Spata16, Spink2, Pgam2, Prm3, Tssk, Prm1 — markers from spermatids and low levels of prophase spermatocyte markers. |
| 4 | Mostly spermatid with contribution from late prophase spermatocytes | Yes | Spata16, Spink2, Pgam2, Prm3, Tssk, Prm2— markers from spermatids and low levels of prophase spermatocyte markers (Sycp3, Sycp1). |
| 2 | Mostly spermatid with contribution from late prophase spermatocytes | Yes | Spata16, Spink2, Pgam2, Prm3, Tssk, Prm2— markers from spermatids and low levels of prophase spermatocyte markers (Sycp3, Sycp2). |
| 5 | Mostly spermatid with contribution from late prophase spermatocytes | Yes | Spata16, Spink2, Pgam2, Prm3, Tssk, Prm2— markers from spermatids and low levels of prophase spermatocyte markers (Sycp3). |
| 11 | Mostly spermatid with contribution from late prophase spermatocytes | Yes | Spata16, Spink2, Pgam2, Prm3, Tssk, Prm2; additional Sycp3, Prm3, Tssk6, Tnp1 — markers from spermatids and low levels of prophase spermatocyte markers. |
| 10 | Mostly spermatid with contribution from late prophase spermatocytes | Yes | Similar to cluster 11; enriched for Spata16, Tssk6, Tnp2 — markers from spermatids and low levels of prophase spermatocyte markers. |
| 12 | Mostly spermatid with contribution from late prophase spermatocytes | Yes | Spata16, Tssk6, Tnp3— markers from spermatids and low levels of prophase spermatocyte markers (Sycp3). |
| 6 | Late prophase spermatocyte to spermatid mixture | Yes | Plk1, Ddx4, Clgn, Rec8, Catsper3 — late prophase, meiotic division and early spermatid marker mix. |
| 8 | Late prophase spermatocytes | Yes | H1t, Brca2, Nme5, Gm49794, Gm10505 — late meiotic prophase spermatocyte markers. |
| 9 | Early and mid prophase spermatocytes | Yes | Stra8, Prss50, Tex15, Dmc1, Prdm9, Ddx4 — early and mid prophase spermatocyte markers. |
| 20 | Early prophase spermatocytes with contribution from differentiating spermatogonia | Yes | Stra8, Ddx4, Prdm9, Dmc1 — differentiating spermatogonia and early meiotic transition markers. |
| 7 | Differentiated spermatogonia and with contribution from early prophase spermatocytes | Yes | Similar to cluster 20; differentiating spermatogonia marker Kit and Dmrt1 expression stronger. |
| 16 | Spermatogonia | Yes | Zbtb16, Uchl1 — spermatogonial markers. |

**Supplementary table 5. Summary of supervised clusters and marker-based cell-type annotations.**

Clusters identified by supervised analysis of spermatogenic cells of single-cell RNA-seq data and annotated based on expression of germline marker genes. Key marker genes identifying dominant cell types in each cluster are listed.

| Cluster number | Celltype | Annotation Reasoning |
|---|---|---|
| 13 | Undifferentiated spermatogonia | **High expression:** undifferentiated spermatogonia markers e.g. *Zbtb16* , *Nanos3* , *Egr4,* and *Neurog3* . **Low expression:** markers of differentiated spermatogonia and later spermatogenic stages. |
| 6 (pool of subclusters 6a and 6b) | Differentiated spermatogonia | **High expression:** differentiated spermatogonia marker *Kit,* broad spermatogonial marker *Dmrt1.* **Low expression:** earliest undifferentiated spermatogonia markers (relative to cluster 13), early meiotic genes (relative to cluster 15). |
| 15 | Preleptotene / Leptotene | **High expression**: meiosis inducer *Stra8,* DSB machinery e.g. *Prdm9, Ccdc36,* and *Rec114,* and early recombination markers, e.g. *Dmc1.* **Low expression:** spermatogonia markers, intermediate (*Meiob* ) or late (*Mlh3* ) recombination markers. **Lower expression relative to clusters 7 and 16:** synaptonemal complex components e.g. *Syce3* and *Sycp1.* |
| 7 (pool of subclusters 7a and 7b) | Zygotene | **High expression:** both early (*Dmc1* ) and intermediate (*Meiob* ) recombination markers, synaptonemal complex components *Sycp2* and *Sycp1* (relative to Cluster 15). **Low expression:** meiosis meiosis inducer *Stra8* , DSB machinery e.g. *Prdm9* , *Ccdc36* , and *Rec114* , which are key markers of preleptotene/leptotene (cluster 15). |
| 16 | Early-to-mid Pachytene | **High expression**: intermediate (*Meiob* ) and some late (*Cntd1* ) recombination markers, synaptonemal complex components *Sycp2* and *Sycp1,* and asynapsis sensor for mid pachytene checkpoint (*Hormad1* ), piRNA pathway components *Tdrd9* and *Piwil2* . **Low expression:** early recombination markers (*Dmc1* and *Brca2* ) relative to clusters 7 and 15, and low but increasing levels of the mid-late pachytene marker *Hist1h1t* relative to cluster 7. |
| 10 | Late Pachytene-Diplotene | **High expression**: post mid achytene marker *Hist1h1t,* late recombination markers *Mlh3, Cntd1, Prr19,* piRNA pathway components *Tdrd9, Piwil2* and *Piwil1.* **Low expression:** early (*Dmc1* and *Brca2* ) and intermediate (*Meiob* ) recombination markers. |
| 5 | Late Prophase to Meiotic Divisions | **High expression:** some late prophase (*Pou5f2* and *Prr19* ) and cell division (*Plk1* and *Ccna1* ) markers. **Low expression**: most recombination markers (*Dmc1, Meiob,* and *Mlh3* ), synaptonemal complex components (*Sycp2* and *Sycp1* ), and piRNA pathway components (*Piwil1* and *Tdrd9* ) relative to clusters 10 and 16. |
| 0 | Meiotic Divisions to Early Spermatids | **High expression:** cell division markers *Ccna1, Plk1* , and round spermatid acrosomal marker *Spink2,* and spermiogenesis markers e.g. *Catsper3* and *Catsper4.* **Low expression:** most recombination markers (*Dmc1* , *Meiob* , and *Mlh3* ), synaptonemal complex components (*Sycp2* and *Sycp1* ), and piRNA pathway components (*Piwil1* and *Tdrd9* ) relative to clusters 16, 10 and 5. |
| 1 | Early Spermatids | **High/relatively high expression:** acrosomal markers *Spata16* and *Spink2,* and round spermatid marker *Spata25* relative to late spermatid clusters 2, 3, 17, 4. **Low expression**: elongating spermatid and sperm markers (*Tssk6, Tnp1, Tnp2, Prm1, Prm2, Prm3* ), relative to late spermatid clusters 2, 3, 17, 4. No or trace expression of meiotic prophase markers. |
| 9 | | |
| 12 | | |
| 18 | | |
| 8 | | |
| 2 | Late Spermatids | **High expression**: elongating spermatid and sperm markers (*Tssk6, Tnp1, Tnp2, Prm1, Prm2, Prm3* ). **Low expression:** early spermatid markers e.g. *Spata16, Spink2* , and *Spata25* (relative to cluster 1,9 ,12,18, 8) |
| 3 | | |
| 17 | | |
| 4 | | |
